# Hamster PIWI proteins bind to piRNAs with stage-specific size variations during oocyte maturation

**DOI:** 10.1101/2020.12.01.407411

**Authors:** Kyoko Ishino, Hidetoshi Hasuwa, Jun Yoshimura, Yuka W. Iwasaki, Hidenori Nishihara, Naomi M. Seki, Takamasa Hirano, Marie Tsuchiya, Hinako Ishizaki, Harumi Masuda, Tae Kuramoto, Kuniaki Saito, Yasubumi Sakakibara, Atsushi Toyoda, Takehiko Itoh, Mikiko C. Siomi, Shinichi Morishita, Haruhiko Siomi

**Affiliations:** Department of Molecular Biology, Keio University School of Medicine, Tokyo 160-8582, Japan; Department of Computational Biology and Medical Sciences, Graduate School of Frontier Sciences, The University of Tokyo, Tokyo 113-0032, Japan; School of Life Science and Technology, Tokyo Institute of Technology, Kanagawa 226-8501, Japan; Graduate School of Science, The University of Tokyo, Tokyo 113-0032, Japan; National Institute of Genetics, Mishima 411-8540, Japan; Department of Biosciences and Informatics, Keio University, Yokohama 223-8522, Japan; Japan Science and Technology Agency (JST), Precursory Research for Embryonic Science and Technology (PRESTO), Saitama, Japan

## Abstract

In animal gonads, transposable elements (TEs) are actively repressed to preserve genome integrity through the Piwi-interacting RNA (piRNA) pathway. In mice, piRNAs are most abundantly expressed in male germ cells, and form effector complexes with three distinct PIWI proteins. The depletion of individual *Piwi* genes causes male-specific sterility owing to severe defects in spermatogenesis with no discernible phenotype in female mice. Unlike mice, most other mammals have four PIWI genes, some of which are expressed in the ovary. Here, purification of PIWI complexes from oocytes of the golden hamster revealed that the size of the piRNAs loaded onto PIWIL1 changed during oocyte maturation. In contrast, PIWIL3, an ovary-specific PIWI in most mammals, associates with short piRNAs only in metaphase II oocytes, which coincides with intense phosphorylation of the protein. An improved high-quality genome assembly and annotation revealed that PIWIL1- and PIWIL3-associated piRNAs appear to share the 5′- ends of common piRNA precursors and are mostly derived from unannotated sequences with a diminished contribution from TE-derived sequences, most of which correspond to endogenous retroviruses (ERVs). Although binding sites for the transcription factor A-Myb are identified in the transcription start site regions of the testis piRNA clusters, the piRNA clusters in the ovary show no well-defined binding motifs in their upstream regions. These results show that hamster piRNA clusters are transcribed by different transcriptional factors in the ovary and testis, resulting in the generation of sex-specific piRNAs. Our findings show the complex and dynamic nature of biogenesis of piRNAs in hamster oocytes, and together with the new genome sequence generated, serve as the foundation for developing useful models to study the piRNA pathway in mammalian oocytes.

**Highlights:**

- The size of PIWIL1-associated piRNAs changes during oocyte maturation
- Phosphorylation of PIWIL3 in MII oocytes coincides with its association with small 19-nt piRNAs
- Improved high-quality genome assembly and annotation identifies young endogenous retroviruses as major targets of piRNAs in hamster oocytes
- PIWIL1- and PIWIL3-associated piRNAs share the 5′-ends of the common piRNA precursors in oocytes

## Introduction

Transposition of mobile DNA elements can cause severe damage by disrupting either the structural or regulatory regions on the host genome (Chuong et al., 2017; Han and Boeke, 2005; Hancks and Kazazian, 2016). To avoid such detrimental effects, many animals have a conserved adaptive immune system known as the piRNA pathway in gonads (Iwasaki et al., 2015; Ozata et al., 2019; Pillai and Chuma, 2012). piRNAs form effector complexes with PIWI proteins, a germline-specific class of Argonaute proteins, to guide recognition through complementary base-pairing, leading to silencing their target transposable elements (TEs) mainly in two ways: post-transcriptional silencing by PIWI-mediated cleavage of target transcripts in the cytoplasm and transcriptional silencing by PIWI-mediated chromatin modifications on target loci. Mutations in genes involved in the pathway can lead to sterility.

Although the mechanisms for generating piRNAs appear to largely differ among animals, the defined characteristics of piRNAs include a predominant length of 26–32 nucleotides (nt), a strong bias for uracil (U) at the 5′-ends (1U-bias), 2-O-methylation of the 3′-ends, and clustering of reads to distinct genomic locations (Iwasaki et al., 2015; Ozata et al., 2019). The characterization of the piRNA populations in *Drosophila* and mouse has led to two models for the biogenesis mechanisms: the ping-pong cycle (Brennecke et al., 2007; Gunawardane et al., 2007) and phased (Han et al., 2015; Mohn et al., 2015), which are intimately connected. Long single-stranded precursors, often more than 10 kb in size, are derived from discrete genomic loci that are now referred to as piRNA clusters or piRNA genes (Aravin et al., 2006; Brennecke et al., 2007; Girard et al., 2006; Lau et al., 2006; Vagin et al., 2006). Two major clusters exist to serve as genomic piRNA source loci: intergenic and genic clusters. Intergenic piRNA clusters are often located in the heterochromatic regions and comprise various types of TEs that tend to be young and potentially active, suggesting that intergenic piRNA clusters provide the host with acquired and heritable memory systems to repress TEs (Brennecke et al., 2007; Khurana et al., 2011). Genic piRNAs mainly arise from 3′ untranslated regions (UTRs) of the protein-coding genes (Robine et al., 2009; Saito et al., 2009). The function of the genic piRNAs is not well understood, but some genic piRNAs show significant complementarity to protein-coding genes (Saito et al., 2009; Gonzalez et al., 2015).

piRNA precursors are cleaved by either endonuclease Zucchini/mitoPLD or by the Slicer activity of PIWIs with pre-existing piRNAs to produce 5′ monophosphorylated piRNA intermediates that are loaded onto PIWI proteins (Brennecke et al., 2007; Gainetdinov et al., 2018; Gunawardane et al., 2007; Han et al., 2015; Ipsaro et al., 2012; Mohn et al., 2015; Nishimasu et al., 2012). PIWI-piRNA complexes then initiate piRNA production that is formed by an amplification loop termed the ping-pong cycle in which reciprocal cleavage of TE and cluster transcripts by PIWI proteins generates new piRNA 5′-ends and amplifies piRNA populations while destroying TE mRNAs in the cytoplasm. The ping-pong cycle produces two classes of piRNAs overlapped by precisely 10 nt at their 5′-ends: one class shows a strong preference for U at their 5′-ends (1U) and the second class shows a preference for adenine at nucleotide 10 (10A). The ping-pong cycle is then accompanied by the phased production of piRNAs downstream of the cleavage site, which further creates a sequence diversity of piRNAs. The 3′-ends of piRNAs are determined either by direct cleavage of Zucchini/mitoPLD (mouse *Zucchini* homolog) or PIWIs or by trimming piRNA intermediates by resecting enzymes (Nibbler in *Drosophila*, Trimmer in silkworm, and PNLDC1 in mouse) (Ding et al., 2017; Hayashi et al., 2016; Izumi et al., 2016; Nishida et al., 2018; Nishimura et al., 2018). piRNA biogenesis is then concluded with 2-O-methylation of the 3′-ends by HENMT1 methylase, which has been hypothesized to stabilize mature piRNAs (Gainetdinov et al., 2018; Horwich et al., 2007; Kirino and Mourelatos, 2007; Saito et al., 2007). The extent of 3′-end cleaving/trimming and consequently piRNA length is determined by the footprint of the PIWI protein, possibly explaining the different size profiles of piRNAs associated with distinct PIWI proteins. Structural studies have shown that the 5′- and 3′-ends of the guide small RNAs, including piRNAs, are recognized by the MID-PIWI and PAZ domains of Argonaute/PIWI proteins, respectively (Matsumoto et al., 2016; Wang et al., 2008; Yamaguchi et al., 2020).

Mammalian PIWI-piRNA pathways have been studied mainly in mice (Pillai and Chuma, 2012). Similar to *Drosophila*, mice express three PIWI proteins, MIWI (PIWIL1), MILI (PIWIL2), and MIWI2 (PIWIL4) in the testis, with varying spatiotemporal expression patterns during spermatogenesis. The non-redundant role of *Piwi* genes in the mouse testis is demonstrated by the fact that depletion of individual *Piwi* genes causes male-specific sterility owing to severe defects in sperm formation (Aravin et al., 2007; Aravin et al., 2008; Carmell et al., 2007; Deng and Lin, 2002; Ernst et al., 2017; Kuramochi-Miyagawa et al., 2008; Pillai and Chuma, 2012; Thomson and Lin, 2009). Each PIWI protein associates with distinct piRNA populations; fetal prepachytene piRNAs and pachytene piRNAs. Prepachytene piRNAs are formed from TE- and other repeat-derived sequences. In contrast, pachytene piRNAs have a higher proportion of unannotated sequences with the diminished contribution of TE sequences, though they still function in TE silencing by guiding MILI and MIWI to cleave TE transcripts (De Fazio et al., 2011; Reuter et al., 2011). A specific transcriptional factor, A-MYB, binds the promoter regions of pachytene piRNA clusters as well as core piRNA biogenesis factors, including MIWI, and initiates their transcription (Li et al., 2013). Some pachytene piRNA clusters are divergently transcribed from bidirectional A-Myb-binding promoters (Li et al., 2013).

Although PIWI genes in fly and zebrafish are expressed in both testes and ovaries, mouse *Piwi* genes are expressed only weakly, if not at all, in the ovary, and depletion of these *Piwi* genes does not affect the female germline. Thus, these findings led to the assumption that the piRNA pathway does not play a role in mammalian oogenesis. Unlike mice, many other mammals possess four distinct PIWI genes (*Piwil1*– *4*), suggesting that piRNA-mediated silencing may differ between mice and mammals with four PIWI genes (Hirano et al., 2014; Sasaki et al., 2003). However, except for mice, little is known about mammalian piRNA pathways, particularly their roles in ovaries. Although recent studies have reported the presence of piRNA-like molecules in mammalian female germ cells, including humans (Kabayama et al., 2017; Roovers et al., 2015; Williams et al., 2015; Yang et al., 2019), it is not yet known whether they play a role in the ovary because of the difficulty of technical and/or ethical application of genetic analysis. Although mice and rats belong to the *Muridae* family of rodents, both of which lack *Piwil3*, the golden Syrian hamster (golden hamster, *Mesocricetus auratus*) belongs to the *Cricetidae* family and has four PIWI genes. Golden hamsters have been used as an experimental rodent model for studying human diseases, particularly for cancer development and many different infectious diseases including COVID-19, because they display many features that resemble the physiology and pharmacological responses of humans (Hirose and Ogura, 2019; Sia et al., 2020). In addition, methods for manipulating gene expression, including the CRISPR/Cas9 system, have been recently employed in golden hamsters (Fan et al., 2014; Hirose et al., 2020). Herein, we analyzed PIWI-associated piRNAs in oocytes and early embryos of golden hamsters, in the hope of applying genetic analysis to the piRNA pathway in the ovary. Our analyses revealed that the size of PIWIL1-associated piRNAs changes during oocyte maturation and that PIWIL3 binds short piRNAs only at the metaphase II (MII) stage of the oocyte, which coincides with phosphorylation of the protein. With an improved high-quality genome assembly and annotation of golden hamster, we showed that PIWIL1- and PIWIL3-associated piRNAs appear to share their 5′-ends. Their contents are similar to those observed with pachytene piRNAs in the mouse testis, but their targets in oocytes are mostly endogenous retroviruses. We further identified ovarian piRNA clusters, and motif search for the transcription start site regions of the piRNA clusters revealed no distinct binding motifs in their upstream regions, although A-Myb-binding motifs were enriched in the upstream regions of the testis piRNA clusters. Our study provides a basis for understanding the roles of the piRNA pathway in mammalian oocytes.

## Results

### PIWI genes are expressed in the oocyte of the golden hamster

To examine the expression of PIWI genes in golden hamster gonads, we performed RNA-sequencing (RNA-seq) analysis using hamster ovary, oocyte, and testis samples and analyzed the expression level of PIWI genes by calculating transcripts per kilobase million mapped (TPM) (Figure 1A). The estimated expression levels of *Piwil1* and *Piwil2* were relatively high in the testis (TPM = 24.24 and TPM = 14.07, respectively). *Piwil1* was also expressed in the ovary (TPM = 4.63), while *PiwiL2* is not detected in the whole ovary and appeared to be only weakly expressed in the oocyte. *Piwil4* appeared to be expressed only in the testis, consistent with previous transcriptome analysis in human oocytes (Yang et al., 2019). Interestingly, *Piwil3* was highly expressed in the oocyte (TPM = 14.60). In sharp contrast, the expression of *Piwil3* could not be detected in the testis. These results are consistent with previous analyses of bovine and human PIWIL3 (Roovers et al., 2015; Yang et al., 2019). To analyze the expression levels of known PIWI-piRNA pathway factors other than PIWI genes, we also calculated the TPM values of the predicted homologous genes using RNA-seq data (Figure S1A). Most of the known PIWI-piRNA pathway factor homologs, including *Mov10l1*, *Mael*, *MVH* (mouse *Vasa* homolog), *MitoPLD*, *Gtsf1*, *Henmt1*, and Tudor domain families (Iwasaki et al., 2015), were expressed in both testes and oocytes, suggesting that both testes and oocytes of hamsters have similar biogenesis pathways to produce piRNAs.

**Figure 1.**
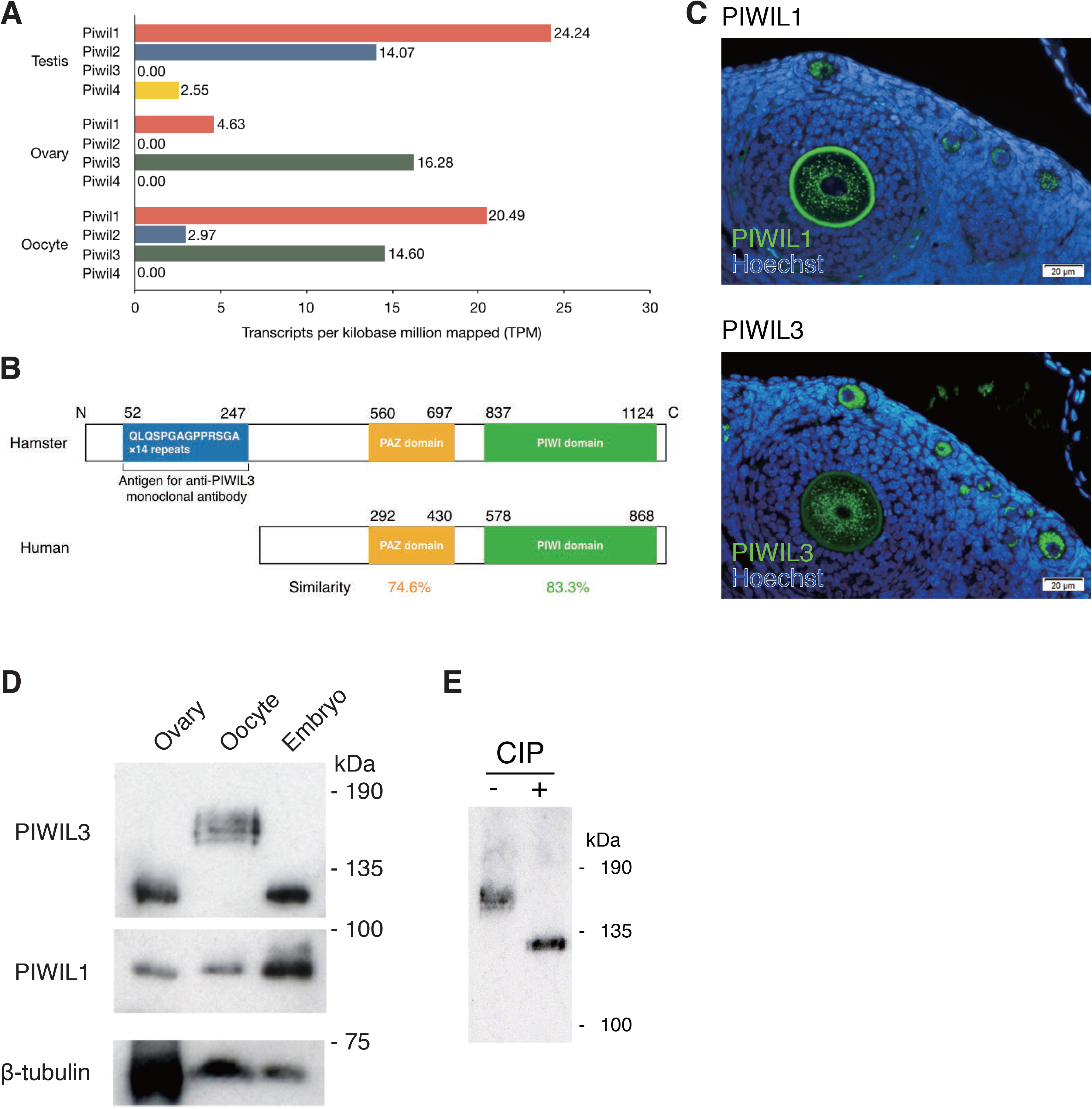
PIWIs are expressed in hamster oocytes. (A) Expression patterns of PIWI families in male and female hamsters. Transcriptome data were obtained from Illumina HiSeq2000. The x-axis suggests the normalized expression level (TPM). (B) Protein homology of PIWIL3 in hamster and human. Approximately 74.6% of the PAZ domain and 83.3% of the PIWI domain in hamster PIWIL3 are conserved in human PIWIL3. The large extension of the amino-terminal portion of the ORF, which contains 14 repeats of nucleotide sequences encoding the amino acid sequences QLQSPGAGPPRSGA is presented in hamster PIWIL3. (C) PIWIL1 (upper panel) and PIWIL3 (lower panel) levels from an ovary. Immunostaining with PIWIL1 and PIWIL3 are shown: (green) PIWIL1 and PIWIL3; (blue) Hoechst. The signal in the zona pellucida at upper panel is the autofluorescence of anti-PIWIL1 antibody. (D) Western blotting was performed from hamster whole ovaries, MII oocytes, and 2-cell embryos with anti-PIWIL3, anti-PIWIL1, and anti-β-TUBULIN antibodies. The size of the PIWIL3 protein largely shifted by approximately 40 kDa in MII oocytes. The anti-MARWI (PIWIL1) antibody (1A5) and anti-PIWIL3 antibody (3E12) detected a single band in each sample except for MII oocytes. (E) Western blotting was performed from hamster MII oocytes with (+) or without (-) CIP treatment. A discrete band migrated to the estimated molecular weight of 130 kDa in the CIP-treated sample, demonstrating that PIWIL3 is heavily phosphorylated in MII oocytes.

To confirm the expression of PIWIs in the ovary, we isolated their cDNAs from the ovary. Open reading frames (ORFs) of sequenced *Piwil1* and *Piwil2* cDNAs correspond well with the respective annotated gene products deposited in the Broad Institute database (MesAur1.0, Broad Institute data) (Figure S1B). However, to our surprise, during the cloning of the *Piwil3* cDNA, we found that the large extension of the amino-terminal portion of the ORF contains 14 repeats of nucleotide sequences encoding the amino acid sequences QLQSPGAGPPRSGA (Figure 1B). To further confirm the expression of PIWIs in the ovary at the protein level, we produced specific monoclonal antibodies against PIWIL3 (Figure S1C). We also found that a monoclonal antibody that recognizes marmoset PIWIL1 (Hirano et al., 2014) cross-reacts with hamster PIWIL1 specifically among hamster PIWI proteins (Figure S1C). Thus, we focused our analysis on PIWIL1 and PIWIL3. Immunostaining with the antibodies produced showed that both PIWIL1 and PIWIL3 were expressed in the cytoplasm of growing oocytes in the ovary (Figure 1C). Western blotting with anti-PIWIL1 antibody showed a discrete band at 90 kDa in the ovary, metaphase II (MII) oocytes, and 2-cell embryos (Figure 1D). Western blotting with anti-PIWIL3 antibody revealed a discrete band at 130 kDa in the ovary and 2-cell embryos, which is consistent with the calculated molecular weight of the protein with the large amino-terminal extension (Figure 1B and 1D). Intriguingly, the size of the protein largely shifted by approximately 40 kDa in MII oocytes (Figure 1D).

### PIWIL3 protein is highly phosphorylated in metaphase II oocytes

This large increase of the PIWIL3 protein in size, together with the broad and fuzzy nature of the band on the gel (Figure 1D), prompted us to test whether this size shift could be due to modification of the protein with phosphorylation. Treatment of MII oocytes with calf intestinal alkaline phosphatase (CIP), an enzyme known to dephosphorylate proteins (Siomi et al., 2002), caused a discrete band to migrate to the estimated molecular weight of 130 kDa, demonstrating that PIWIL3 is heavily phosphorylated in MII oocytes (Figure 1E). These results show that PIWIL1 and PIWIL3 are expressed in growing oocytes in the ovary as well as in early embryos and that PIWIL3 is modified with phosphorylation specifically at the MII stage of oocytes. Since the mouse genome lacks *Piwil3* and thus the characterization of PIWIL3 protein has been delayed, our findings indicate that *Piwil3* may have specific functions in female gonads.

### The size of piRNAs loaded onto PIWIL1 changes during oocyte maturation

To isolate PIWIL1- and PIWIL3-associated small RNAs from oocytes, we immunopurified the associated complexes from MII oocytes with specific monoclonal antibodies produced. We then isolated RNAs, ^32^P-labeled them, and analyzed them using a denaturing polyacrylamide gel (Figure 2A). Intriguingly, PIWIL1 was associated with two populations of small RNAs: one with 29–30 nt and the other with 22–23 nt in MII oocytes. We then immunopurified PIWIL1-associated complexes from whole ovaries (including growing oocytes) and 2-cell embryos. The sizes of PIWIL1-associated piRNAs in whole ovaries and 2-cell embryos were 29–30 nt and 22–23 nt, respectively (Figure 2B). The resistance of PIWIL1-associated piRNAs in MII oocytes to periodate oxidation (NaIO4) and β-elimination reactions show that they are modified at the 3′ terminal nucleotide with a 2′-O-methyl marker (Figure S2A). We also isolated PIWIL1-associated small RNAs from whole testes and found that small RNAs (29–30 nt) were loaded onto PIWIL1 in the testis (Figure S2B). These results show that piRNAs loaded onto PIWIL1 change their sizes during oocyte maturation, from the size equivalent to that observed in the testis to a mixture of long and short populations, and short piRNAs (22– 23-nt). To our knowledge, this is the first study to show that the size of piRNAs loaded onto a distinct PIWI protein changes during germline development.

**Figure 2.**
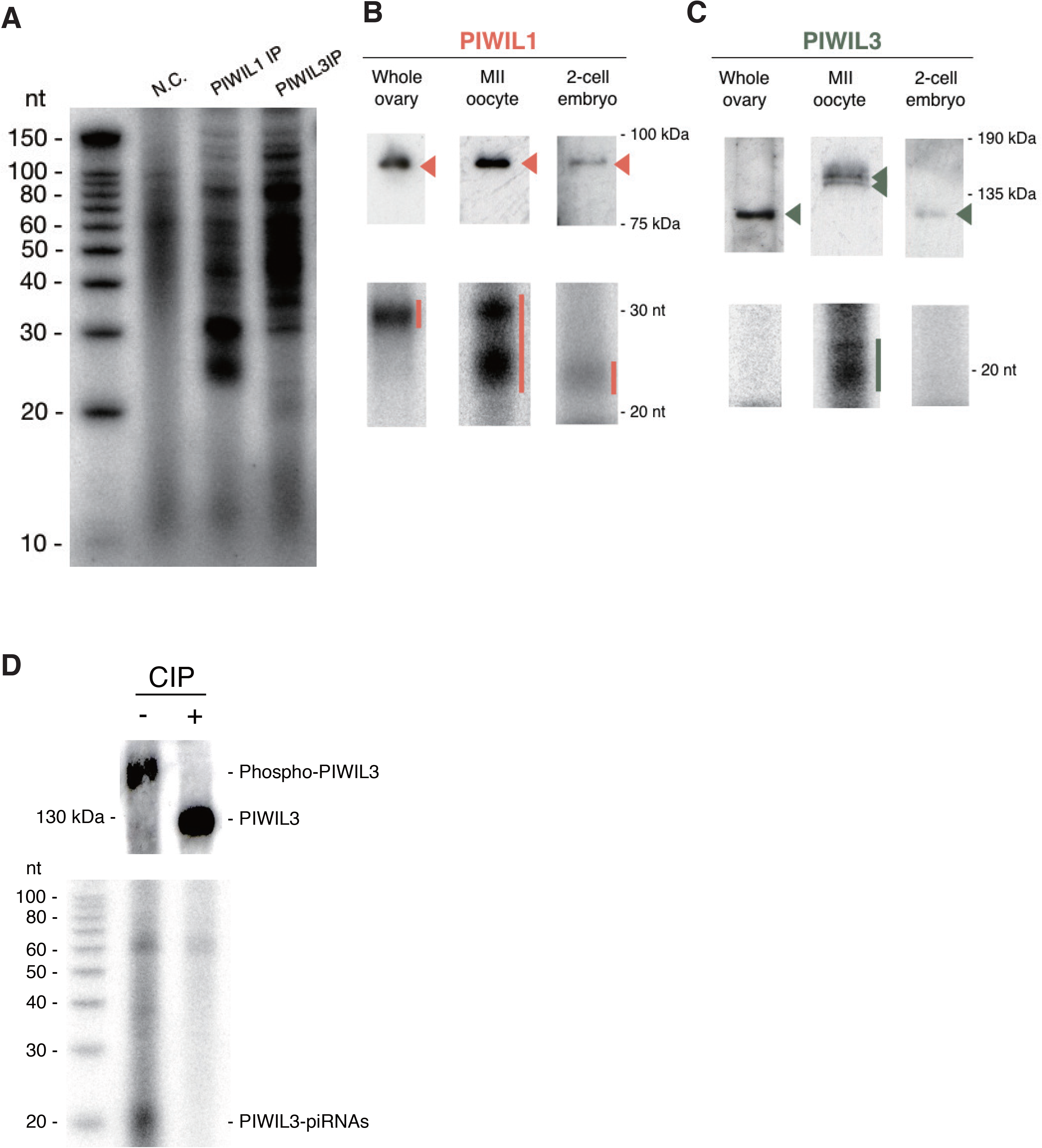
The different populations of piRNAs associate with each of the PIWI proteins at different developmental stages. (A) Isolated RNAs from PIWIL1 and PIWIL3 immunoprecipitates in MII oocytes were ^32^P-labeled and separated by a denaturing polyacrylamide gel. PIWIL1 and PIWIL3 proteins associate with ∼28∼31-nt and ∼18∼20-nt-long piRNAs, respectively. (B) Western blotting and ^32^P-labeling were performed from PIWIL1 immunoprecipitates in hamster whole ovaries, MII oocytes, and 2-cell embryos. Upper panel: Western blotting; lower panel: ^32^P-labeling. (C) The same experiments with (B) were performed from PIWIL3 immunoprecipitates. (D) Phosphorylation of the protein is required for PIWIL3 probably either to get loaded with piRNAs or hold them or both. PIWIL3 immunoprecipitates were treated with (+) or without (-) CIP. Upper panel: Western blotting; lower panel: ^32^P-labeling of RNAs, which were purified from each PIWIL3 immunoprecipitate.

### Phosphorylated PIWIL3 appears associated with short piRNAs only in MII oocytes

In sharp contrast, PIWIL3 was found to bind to a class of small RNAs with 19 nt (Figure 2A), which is consistent with the recent finding that human PIWIL3 associates with small RNAs (∼20 nt) in human oocytes (Yang et al., 2019). 19–20 nt RNAs in oocytes almost completely disappeared after β-elimination reactions, indicating that PIWIL3-associated piRNAs lack 2′-O-methylation at their 3′ terminal (Figure S2C). We failed to detect small RNAs associated with PIWIL3 in whole ovaries and 2-cell embryos (Figure 2C). With the caveat that this could be because of technical reasons for immunoprecipitation with the antibodies and/or the buffer conditions used, our findings suggest that PIWIL3 may bind piRNAs with 19 nt exclusively in MII oocytes but be freed from piRNAs as ‘empty’ PIWIL3 at the early stages of oocyte maturation and in early embryos. PIWIL3 is heavily phosphorylated only in MII oocytes, raising the possibility that phosphorylation of the protein may be required for the association with a class of short piRNAs. To test this, we performed immunoprecipitation with an anti-PIWIL3 antibody using MII oocyte lysate that had been treated with CIP and examined whether the CIP treatment affected the association of piRNAs with PIWIL3. Indeed, PIWIL3 treated with CIP was free from piRNAs (Figure 2D). This shows that phosphorylation of the protein is required for PIWIL3 probably either to get loaded with piRNAs or hold them or both.

### Generating the hamster genome assembled by resequencing the whole genome to accurately map piRNAs

Although the draft genome of the golden hamster has been sequenced (the MesAur1.0 genome), we soon came to realize that we needed much more accurate genome sequence data to further characterize these PIWI-associated piRNAs on the genome mainly because the MesAur1.0 genome sequence contains a large number of gaps (N) (17.58% of the genome; 420 Mb of the 2.4 Gb) and remains fragmented. Because TEs and other repeats in the genome are the main targets of piRNAs in many animals, the lack of accurate sequences of TEs and other repeats is a serious problem when mapping piRNAs on the genome. Accurate detection of TEs requires both full collection/classification of TE consensus sequences and high-quality genome assembly. Thus, we re-sequenced the golden hamster genome.

Details of the hamster genome assembly are shown in the Methods section. The final genome assembly is summarized in Table 1. The nucleotide difference between the DNA Zoo MesAur1.0_HiC assembly and our assembly was 0.140%. We assessed the completeness of the genome assemblies using the BUSCO tool (Waterhouse et al., 2018) and found that our golden hamster genome assembly included 3,991 complete genes (97.2%) and 37 fragmented genes among 4,104 single-copy genes. Our new golden hamster genome allowed us to resolve several issues that had remained ambiguous. For example, although putative ancestral karyotypes of rodents in the *Muridae* and *Cricetidae* families have been partly reconstructed by traditional chromosome painting (Romanenko et al., 2012), we compared our nearly complete genome with the mouse (*Mus musculus*) and rat (*Rattus norvegicus*) reference genomes (Methods) and identified conserved synteny blocks between the golden hamster, mouse, and rat genomes (Figure 3A). We inferred ancestral karyotypes by integrating synteny blocks shared between two or three species according to maximum parsimony, to minimize the amount of chromosomal rearrangement (Methods) and obtained a high-resolution ancestral karyotype of *Muridae* using the golden hamster genome as the outgroup as well as a precise ancestral *Cricetidae* karyotype (Figure 3B). Although our ancestral *Cricetidae* and *Muridae* karyotypes were consistent with most previous inference (Romanenko et al, 2012), we resolved some problems: for example, it was unclear whether mouse chromosomes 3 and 4 possessed synteny blocks from a common ancestral karyotype, and our analysis demonstrated the existence of a proto-chromosome in the ancestral *Cricetidae* karyotype (brown blocks in Figure 3B). We also confirmed previous speculation that mouse chromosomes 5 and 16 obtained blocks from a common *Muridae* proto-chromosome (light orange in Figure 3B), and that chromosomes 10, 15, and 17 obtained blocks from a common *Cricetidae* proto-chromosome (maroon blocks in Figure 3B). Finally, we identified two groups of mouse chromosomes (6, 8) and (8, 13) having large blocks from common *Muridae* proto-chromosomes.

**Figure 3.**
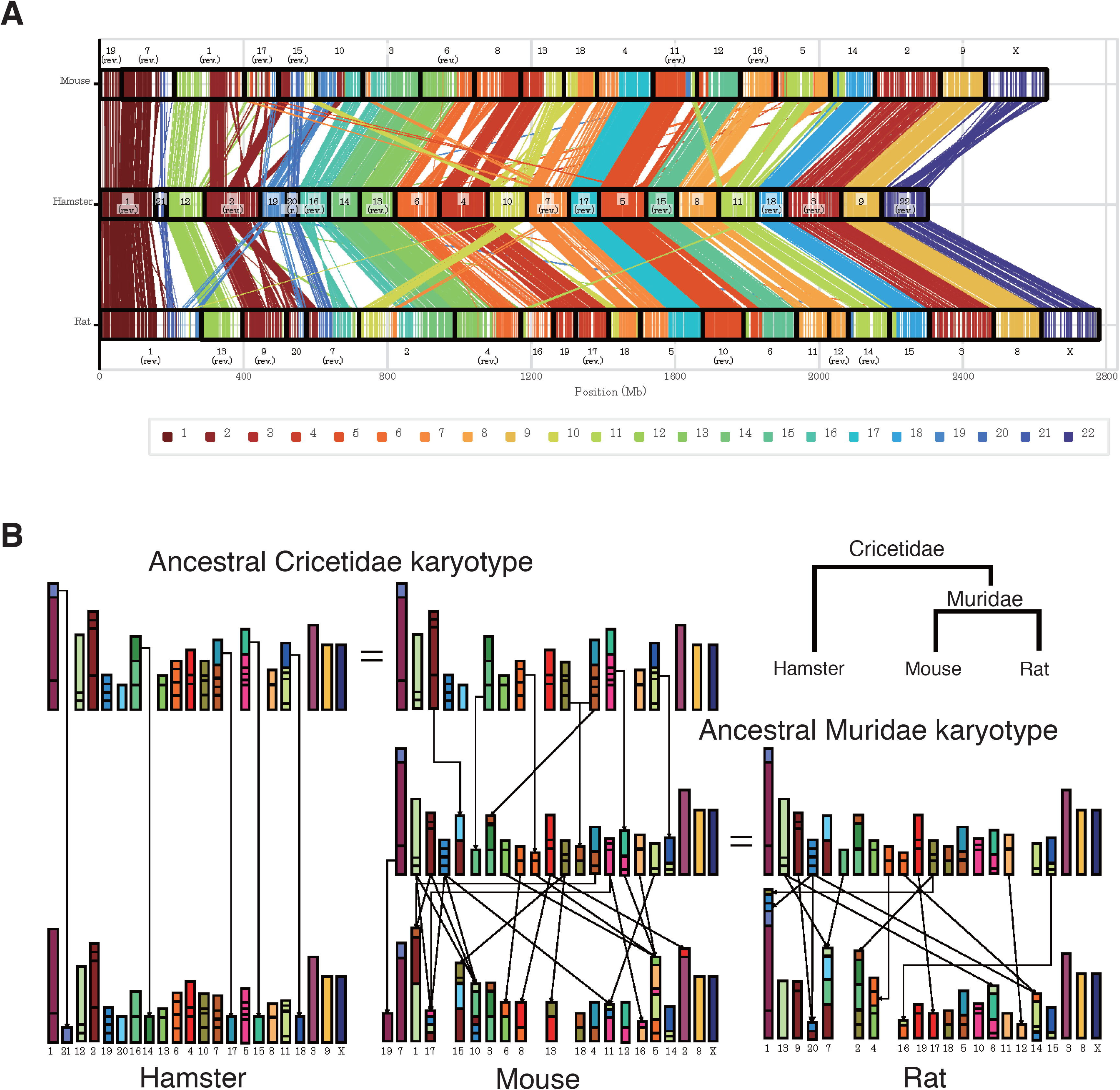
Chromosomal evolution in Rodentia. (A) Each line represents a reciprocally best-matching pair of positions in our hamster genome (middle) and the mouse (top) or rat (bottom) reference genome. In each line, the colored part represents the hamster chromosome, and the color palette displayed at the bottom shows the color-coding. Neighboring lines indicate synteny blocks conserved between two species. (B) Schematic showing the ancestral karyotypes of *Cricetidae* and *Muridae* inferred from conserved synteny blocks using maximum parsimony. The top two karyotypes are identical and represent the ancestral *Cricetidae* karyotype; the middle two show the ancestral *Muridae* karyotype. Each box represents a conserved synteny block among the three species; the colored part indicates the hamster chromosome as per Figure A. Small synteny blocks are enlarged. Lines from one proto-chromosome to the descendant chromosomes indicate rearrangements (fusion, fission, or translocation). To simplify the graph, we omitted lines between the identical chromosomes within the same columns. Brown blocks show the existence of a proto-chromosome in the ancestral *Cricetidae* karyotype. We confirmed the previous speculation that mouse chromosomes 5 and 16 obtained blocks from a common *Muridae* proto-chromosome (light-orange), and that chromosomes 10, 15, and 17 obtained blocks from a common *Cricetidat*e proto-chromosome (maroon blocks). We also identified two groups of mouse chromosomes (6, 8) and (8, 13) having large blocks from common *Muridae* proto-chromosomes.

**Table 1.**
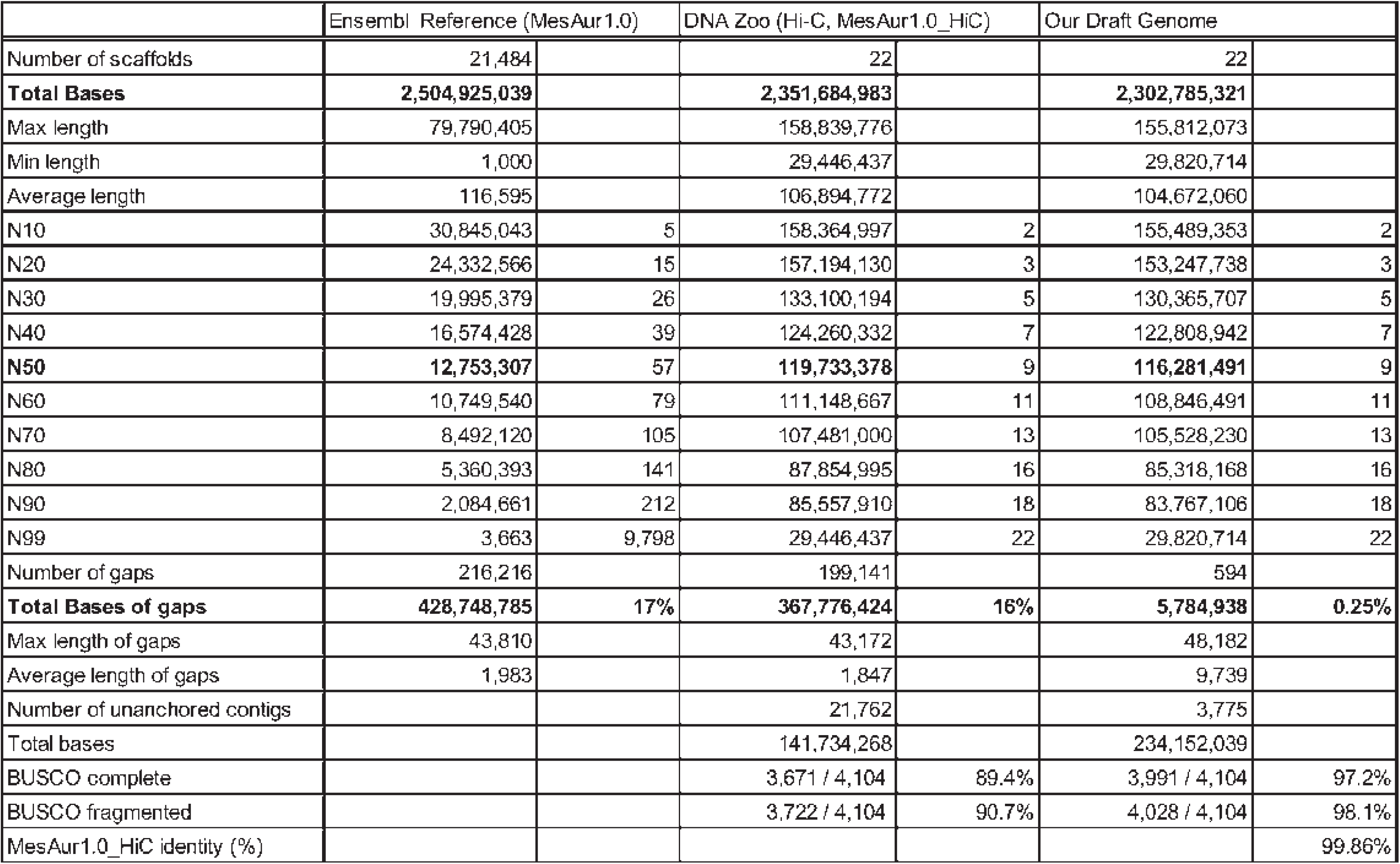
The statistics of the genome assemblies.

Using our genome assembly, we compared the hamster genomic locus containing the *Piwil3* gene with syntenic regions of the mouse and rat genomes. The *Piwil3* gene is flanked by *Wscd2* and *Sgsm1* in the hamster genome (Figure S3A). The order of the two genes is conserved in the syntenic regions of the mouse and rat genomes. We then extracted the genomic regions between these genes from our hamster genome and the reference genomes of mouse (mm10) and rat (rn6), compared the syntenic regions using dot plots (Figure S3B), and observed the absence of the *Piwil3* gene. The validity of the hamster genomic region with *Piwil3* was confirmed by checking each base in the region was covered by an ample number of long reads (Figure S3C). In addition, we performed a similarity search with blastn and ssearch36 using the protein-coding sequence (CDS) of hamster *Piwil3* as a query and found no hits in the corresponding regions of the mouse and rat genome. The PIWIL3 gene is conserved in most mammals, including humans, suggesting that *Piwil3* might have been deleted after speciation in mice and rats.

With the new golden hamster genome sequence generated, we also conducted a *de novo* repeat characterization and identified 177 consensus sequences of repetitive elements at the subfamily level, including 3 SINEs, 12 LINEs, 156 long terminal repeat (LTR) retrotransposons, and 2 DNA transposons. RepeatMasker analysis using our custom repeat library (RepeatMasker rodent library with newly identified 177 consensus sequences) revealed that SINEs, LINEs, LTR retrotransposons, and DNA transposons occupy 9.2%, 16.9%, 12.1%, and 1.3% of the hamster genome, respectively. The contents of TEs are equivalent to those found in mice and rats, but the fractions of SINE and LINE are, respectively, higher and slightly lower in the hamster than those observed in mice and rats (Table 2). The contents of the MesAur1.0 genome (MesAur1.0_HiC) were similarly analyzed. In contrast to our assembly, a much lower proportion of young TEs were detected even using our custom repeat library (Figure S4A, Table S4). This is mostly because of the high frequency of gaps (Ns) in the MesAur1.0_HiC assembly (Table 1), which resulted in apparently less similarity between TEs and their consensus sequences.

**Table 2.** Proportion (%) of transposable elements in rodents.

| Class | Clade/Superfamily | Hamster | Mouse | Rat |
| --- | --- | --- | --- | --- |
| SINE | B1 | 3.30 | 2.32 | 1.56 |
|  | B2 | 1.96 | 2.16 | 2.10 |
|  | B4 | 2.60 | 2.13 | 2.12 |
|  | ID | 0.34 | 0.17 | 0.72 |
|  | MIR | 0.72 | 0.56 | 0.53 |
|  | other SINE | 0.24 | 0.01 | 0.01 |
| LINE | L1 | 16.22 | 19.75 | 18.94 |
|  | L2 | 0.57 | 0.43 | 0.41 |
|  | other LINE | 0.13 | 0.10 | 0.10 |
| LTR | ERV1 | 0.64 | 0.74 | 0.99 |
|  | ERVK | 5.02 | 5.02 | 3.81 |
|  | ERVL | 5.55 | 5.90 | 5.26 |
|  | other LTR | 0.87 | 0.44 | 0.04 |
| DNA | hAT | 0.98 | 0.82 | 0.81 |
|  | TcMar | 0.27 | 0.22 | 0.21 |
|  | other DNA | 0.03 | 0.03 | 0.03 |
| Other TE |  | 0.38 | 0.29 | 0.27 |
| Unknown TE |  | 0.09 | 0.12 | 0.79 |
| **Total** |  | **39.90** | **41.22** | **38.70** |

The custom library substantially improved the detection ability of recently active TEs, as represented by a higher proportion of young elements, for example, those with low (< 5.0) Kimura two-parameter (K2P) divergence from the consensus (Figure 4A). Hamster-specific subfamilies of B1 SINE were recently active, and 36,000 copies of young full-length B1s are present in the assembly. We identified three groups of the LINE-1 (L1) family, two of which were recently highly active (L1-4_MAu and L1-5_MAu), and the genome harbors at least 1,000 young full-length copies (Figure 4B). The most active LTR retrotransposons belong to the ERV2/ERVK superfamily, including IAP (for ERV classification, see Kojima 2018; Vargiu et al., 2016). We identified 20 families of ERV2 that contain an internal portion between LTRs, and 11 of them possess a clear reverse transcriptase domain. There are over 13,000 LTR sequences and 1,600 internal portions of the recently active elements in the genome. Among them, we found three recently expanded families: IAP1E_MAu, ERV2-5_MAu, and ERV2-7_MAu, which accounted for 29.9%, 14.7%, and 26.8% of the very young (<2.0 K2P divergence) LTR retrotransposons, respectively (Figure 4C). It is likely that not only IAP but also other ERV2/ERVK elements are currently active in the hamster genome. Together, these results show that our effort to re-sequence the golden hamster genome significantly improved annotations, especially for recently active TEs, which are potential piRNA targets.

**Figure 4.**
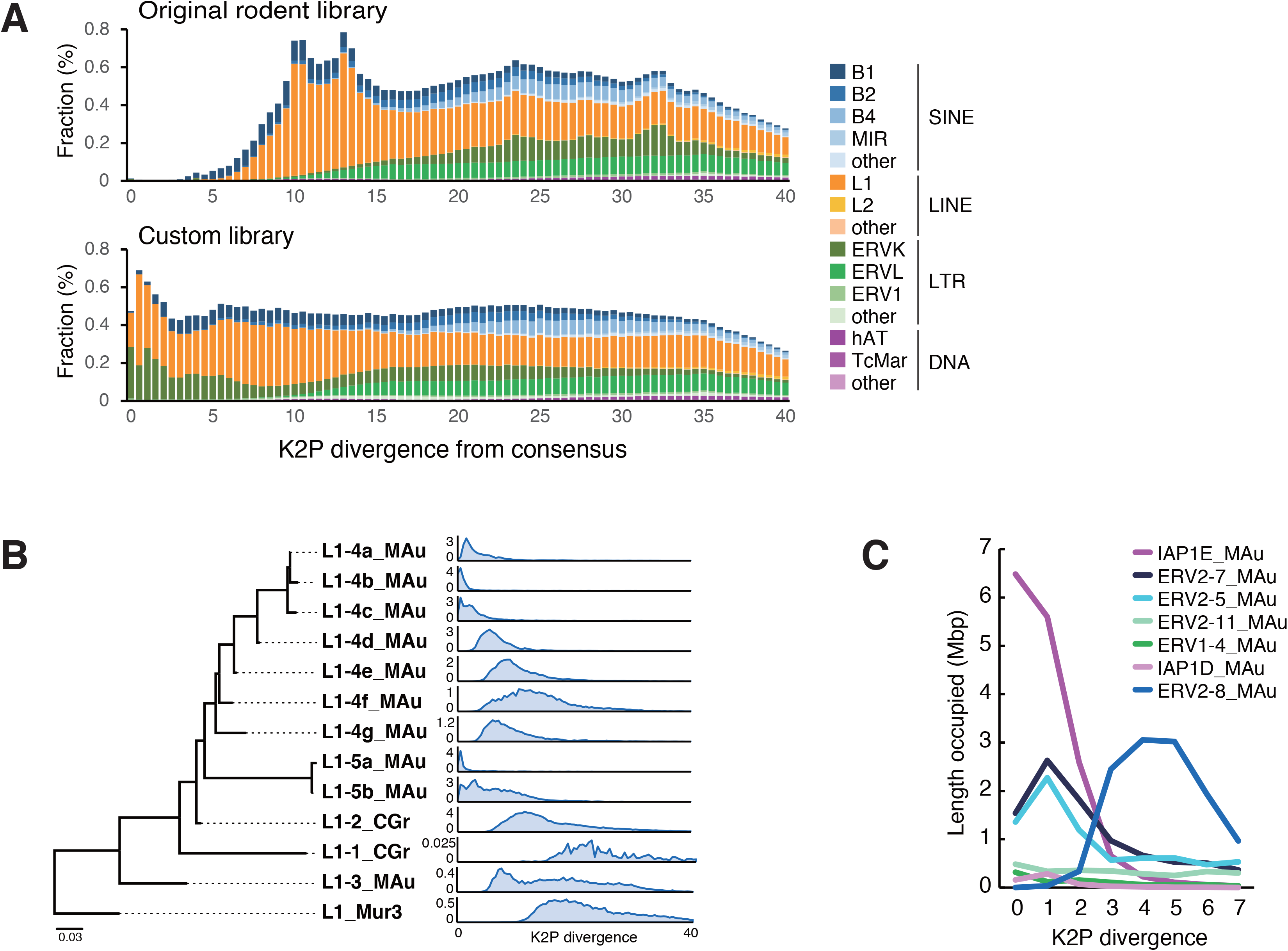
Identification of active transposable elements in the hamster genome. (A) Age distribution of the hamster TEs compared between the original and customized repeat libraries. The proportion of TEs is shown for 0.5 bins of Kimura 2-parameter (K2P) distance (CpG-corrected) from each consensus sequence. (B) A Maximum-likelihood tree of the L1 ORF2 subfamilies and their age distribution (0.5 bins of K2P distance). The TE amount is the length occupied in Mbp. (C) Amount of recently-active LTR retrotransposons compared among representative IAP and other ERV2 families.

### Characterization of piRNAs loaded onto PIWIL1 or PIWIL3

To characterize piRNAs loaded onto PIWIL1 or PIWIL3 in oocytes, we performed small RNA sequencing using piRNA samples immunopurified with an anti-PIWIL1 or an anti-PIWIL3 antibody from oocytes. For PIWIL1-associated piRNAs, we also sequenced piRNA samples immunopurified with an anti-PIWIL1 antibody from testes. Replicates were highly correlated with each other (R^2^ > 0.9) (data not shown); therefore, we merged the reads. First, we performed a length distribution analysis of the obtained reads (Figure 5A-E). We observed peak signals at the size range of 29–30 nt in the testes and ovaries, 29 nt and 23 nt in MII oocytes, and 23 nt in 2-cell embryos of PIWIL1-associated piRNAs and 19 nt in MII oocytes of PIWIL3-associated piRNAs, confirming that the length of PIWIL1- and PIWIL3-associated piRNAs sequenced is consistent with that observed on the gels. Then we divided into two groups using the length of PIWIL1-associated piRNAs in MII oocyte; Oocyte long piRNAs (L1 OoL-piRNAs) and Oocyte short piRNAs (L1 OoS-piRNAs) for further analysis. Sequencing also revealed that the piRNA populations in all samples showed a strong 1U bias, a conserved characteristic for piRNAs (Figure 5A–E).

**Figure 5.**
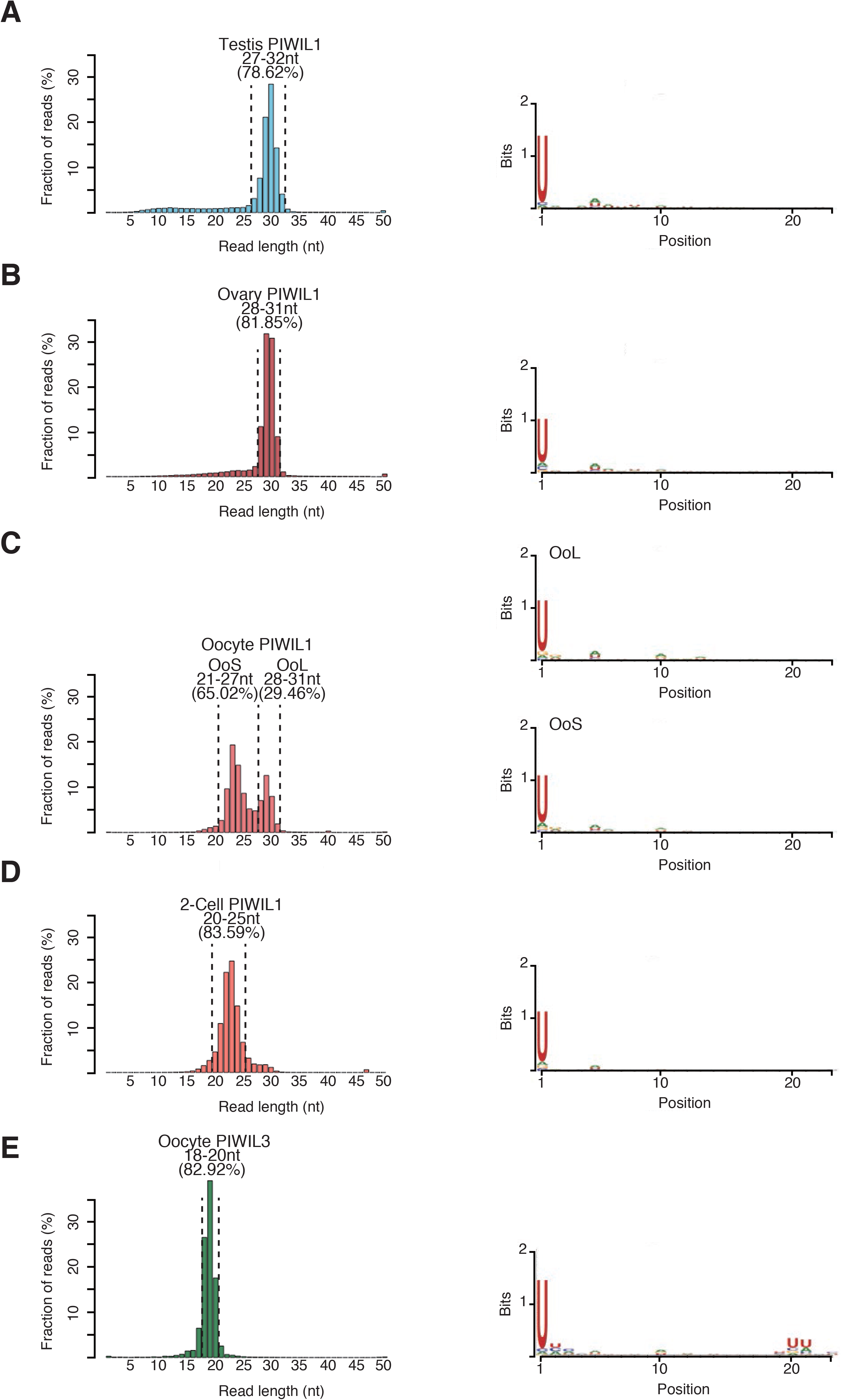
Characterization of PIWI-associated piRNAs in hamster male and female gonads. Size distribution and nucleotide bias in PIWIL1-associated piRNAs in hamster (A) testis, (B) ovary, (C) MII oocyte, (D) 2-cell embryo, and (E) PIWIL3-associated piRNAs in hamster MII oocytes. Left panel: size distribution; peaks of size distribution are shown at 29–30 nt in testis and ovary PIWIL1-piRNAs, 29–30 nt and 23–24 nt in MII oocyte PIWIL1-piRNAs, 22–23 nt in 2-cell embryo PIWIL1-piRNAs and 19 nt in MII oocyte PIWIL3-piRNAs, respectively. The right panel shows nucleotide bias. All piRNA populations have a strong uridine (U) bias at their 5′-end.

We then mapped piRNAs to the new hamster genome (hamster.sequel.draft-20200302.arrow.fasta). Among the PIWI-associated piRNA reads, 50.0∼67.4% of the reads were uniquely mapped to the genome and 6.2∼43.3% were mapped multiple times (Figure 6A, upper panel). Then, we annotated each piRNA read to analyze the genomic region from which the piRNA reads were generated. Approximately 79.05∼89.84% of the reads mapped to unannotated intergenic regions, and only 7.07∼13.19% originated from TEs (Figure 6A, lower panel). This profile is similar to that of pachytene piRNAs associated with mouse MIWI, in which ∼70% is derived from intergenic regions and ∼24% from TEs (Reuter et al., 2011).

**Figure 6.**
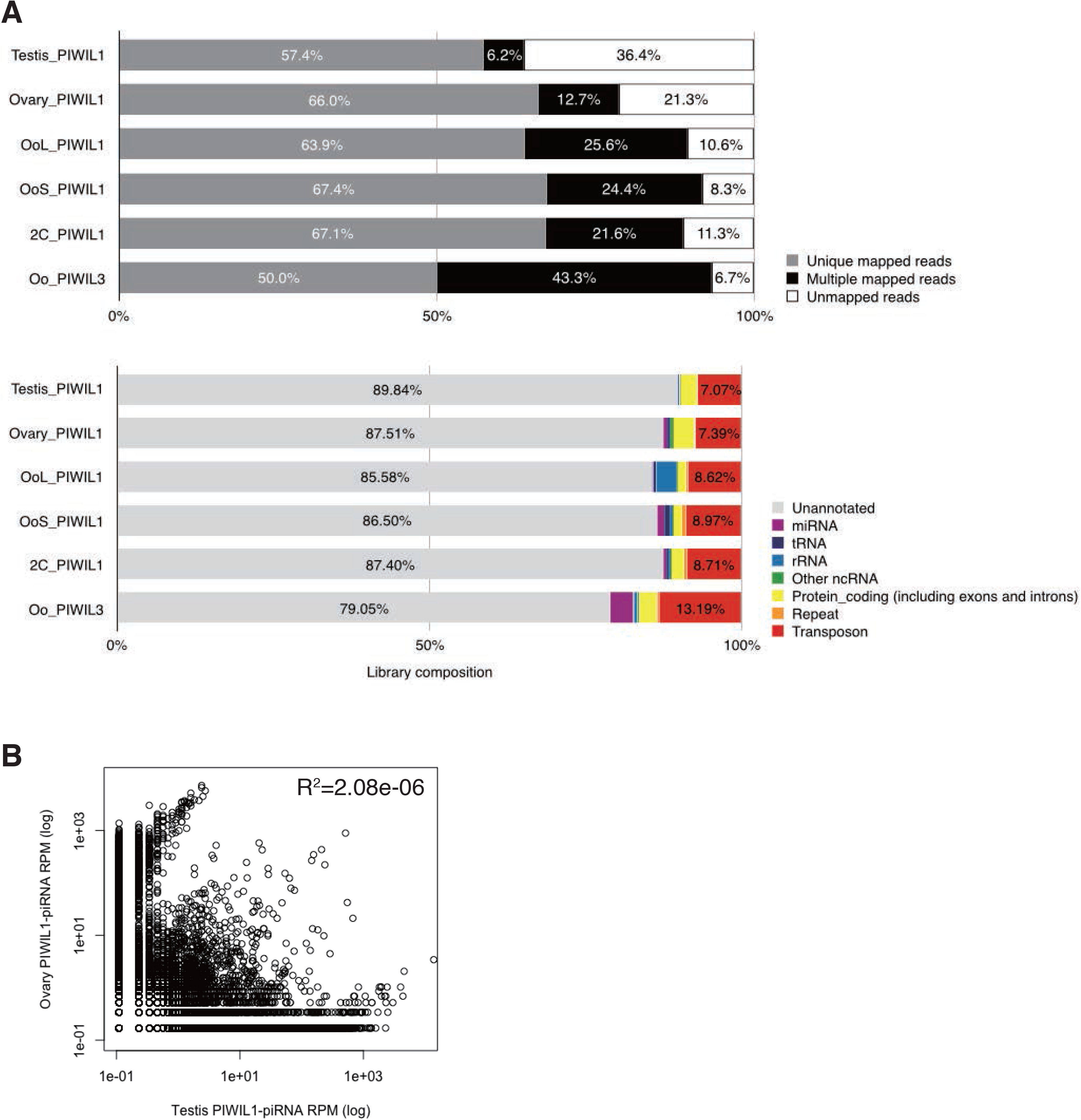
Characterization of genome mapped PIWI-associated piRNA reads in hamster male and female gonads. (A) Genome mapping ratio and annotation of PIWIL1- and PIWIL3-associated piRNA reads in hamster testis, ovary, MII oocyte, and 2-cell embryo. The upper panel exhibits the genome mapped ratio of each PIWI-associated piRNA reads: (gray) unique mapped reads; (black) multiple mapped reads; (white) unmapped reads. Lower panel exhibits the results of annotation of genome mapped reads: (red) transposon; (orange) repeat; (yellow) protein-coding gene, including exon and intron; (purple) miRNA; (blue) tRNA; (skyblue) rRNA; (green) other ncRNA, including snoRNA and snRNA. (B) Comparison between testis and ovary PIWIL1-associated piRNAs. Pearson correlation coefficient was calculated using R. Each dot exhibits the specific piRNA reads.

The characteristics of PIWIL1-piRNAs were virtually indistinguishable between the testis and ovary. To determine whether this was due to the common piRNA sequences, we calculated reads per million mapped reads (RPM) for each piRNA sequence and compared the correlation between testis and ovary samples. The RPM of piRNAs greatly varied between the testis and ovary samples (R^2^ = 2.08e-06). In addition, when we checked the top ten piRNA sequences with the most abundant read counts, none of the sequences were common between the testis and ovary (data not shown). These results show that testis and ovary PIWIL1-piRNAs possess distinct sequences (Figure 6B).

We found that most piRNAs in hamster female gonads were derived from LTR retrotransposons, including endogenous retroviruses (ERVs), compared to PIWIL1-piRNAs in the hamster testis, which corresponded to both LINEs and LTR retrotransposons (Figure S5). This observation is consistent with the fact that the main targets of the piRNA pathway in mouse testes are L1 and IAP elements. In the mouse genome, L1 is the most active TE family, as represented by an increasing accumulation of their young copies (Figure S4B). There are also young LTR retrotransposons in mice, among which IAP and MERVL (ERV3/ERVL) families are highly active, while ERV2/ERVK families, except IAPs, showed much lower proportions among the young elements (Figure S4C). Interestingly, most LTR retrotransposons from which piRNAs in hamster female gonads were derived were evolutionally younger judged by the K2P divergence (Figure 4C). This suggests that hamster oocyte piRNAs were generated from the higher activity of recent transposition. In contrast, testis PIWIL1-piRNAs target both LINEs and LTR retrotransposons. Together with the diversity in PIWI-associated piRNA sequences of the oocyte and testis (Figure 6B), this supports the notion that PIWI-piRNA target TEs are different for male and female gonads.

### Relationships among PIWIL1- and PIWIL3-bound piRNAs in hamster oocytes

Figure 2 and 5 show that piRNAs loaded onto PIWIL1 in MII oocytes consist of two distinct populations, short piRNAs (L1 OoS-piRNAs) and long piRNAs (L1 OoL-piRNAs). Shorter piRNAs (18–20 nt) were loaded onto PIWIL3 in MII oocytes. These findings prompted us to test whether these piRNAs are derived from the same genomic loci and whether L1 OoS-piRNAs and/or PIWIL3-associated piRNAs are processed products of L1 OoL-piRNAs by cleaving and/or trimming their 3′-ends. To this end, we asked whether the genomic positions of the 5′- or 3′-ends of the L1 OoL-piRNAs differ from those of L1 OoS-piRNAs or PIWIL3-piRNAs and calculated the probabilities of the 5′- or 3′-ends of L1 OoL-piRNAs coinciding with the 5′- or 3′-ends of L1 OoS-piRNAs or PIWIL3-piRNAs (Gainetdinov et al., 2018). In MII oocytes, L1 OoL-piRNAs were much more likely to share 5′-ends with L1 OoS-piRNAs and PIWIL3-piRNAs (Oo PIWIL3) than with 3′-ends (0.65 for 5′-ends versus 0.03 for 3′-ends and 0.54 for 5′-ends versus 0.02 for 3′-ends, respectively; Figure 7A). These results suggest that approximately half of these three piRNA populations share the same 5′ cleaved piRNA precursors, thereby sharing the same piRNA clusters as piRNA sources. Thus the size differences among these piRNAs account for their 3′-end formation of these populations that may be determined by the footprint of piRNA-binding PIWI proteins, probably because of either the structures of partner PIWI proteins (in particular, the distance between MID-PIWI and PAZ domains) or conformational changes in the partner proteins (see Discussion).

**Figure 7.**
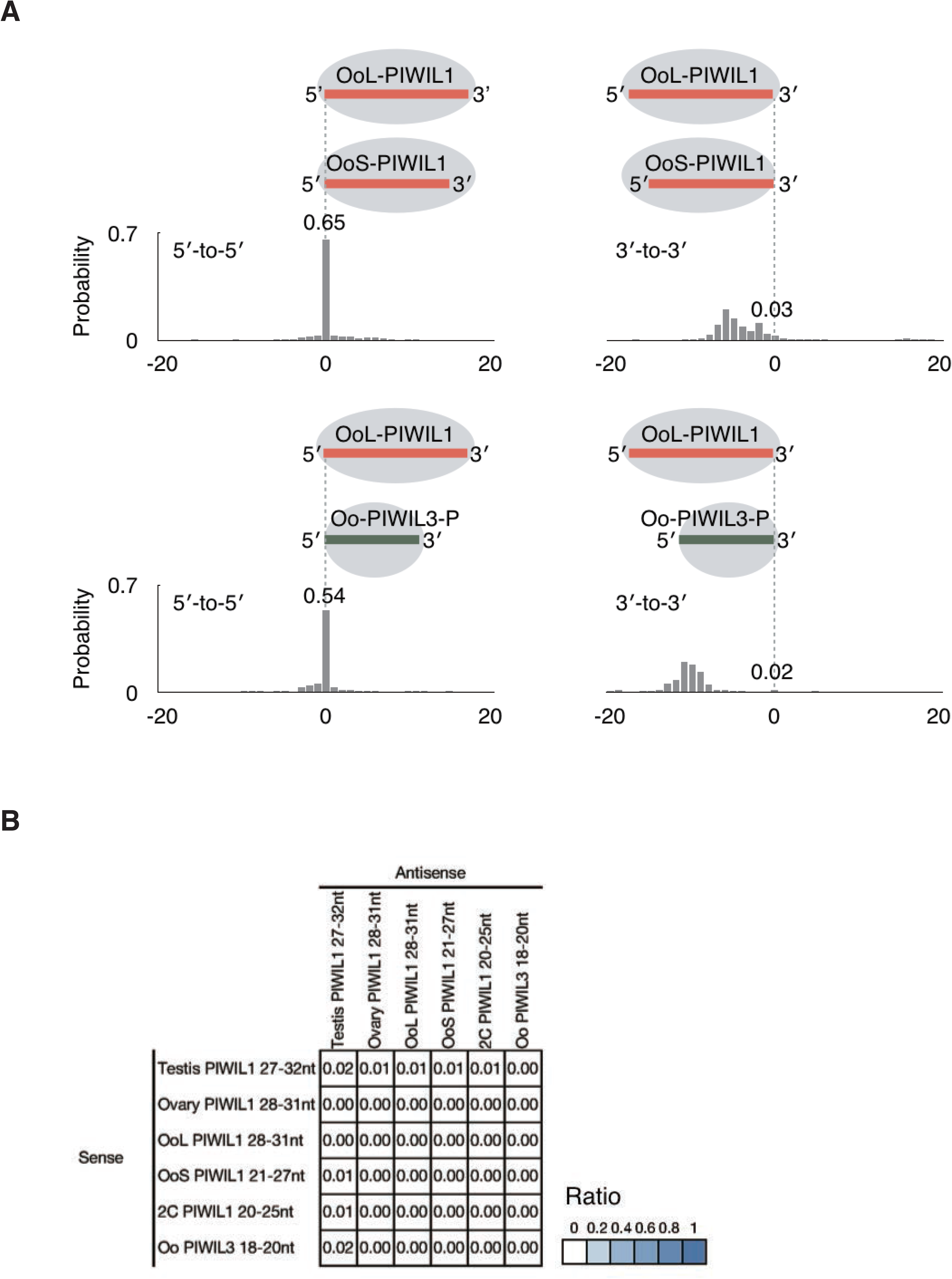
PIWIL1- and PIWIL3-associated piRNA populations are likely to share the same 5′ cleaved piRNA precursors. (A) Probability of distances between the 5′- or 3′-ends of OoL PIWIL1 (longer)- and OoS PIWIL1- or PIWIL3 (shorter)-piRNAs (Oo PIWIL3). The numbers indicate the total frequency of the 5′- or 3′-ends of shorter piRNAs residing before, after, or coinciding with the 5′- or 3′-ends of the longer piRNAs. This indicates that piRNA populations are likely to share the same 5′ cleaved piRNA precursors, but the 3′-end formation of these populations may be determined by the footprint of piRNA-binding PIWI proteins probably because of either structure. (B) A heat map showing the probability that the first base of the PIWI-associated piRNA reads mapped to the antisense strand overlap the 10th base of the reads, which mapped to the sense strand. This result shows that they do not have any ping-pong signals with each other.

The finding that piRNA populations in oocytes share the same 5′ cleaved piRNA precursors suggests little ping-pong amplification among them. We calculated the ping-pong signature of each piRNA and found that they had little or no ping-pong signatures (Figure 7B). This result indicates that most of the piRNAs expressed in oocytes are not involved in the ping-pong cycle. Recently, a novel class of small RNAs with 21– 23 nt, termed short PIWI-interacting RNAs (spiRNAs), was identified in mouse oocytes (Kabayama et al., 2017). They are produced by the ping-pong cycle driven by the MILI slicer activity, and thus, small piRNAs found in hamster oocytes are distinct from spiRNAs.

### piRNA clusters in the oocytes are distinct from those observed in the testis

We found that more than half of PIWIL1- and PIWIL3-associated piRNAs are likely to share 5′-ends of precursor transcripts, suggesting that the primary source of piRNAs is also shared among them. To test the possibility that PIWIL1- and PIWIL3-associated piRNAs share piRNA clusters from which they are derived, we next focused on the identification of piRNA clusters in hamster testes, ovaries, MII oocytes, and 2-cell embryos. We identified piRNA clusters using proTRAC (version 2.4.3) under the following conditions as described (Yang et al., 2019) with some modifications: (1) more than 75% of the reads were appropriate to the length of each piRNA; (2) more than 75% of the reads exhibited the 1 U or 10 A preference; (3) more than 75% of reads were derived from the main strand; and (4) -pimin option with 21, 28, and 18 for oocyte PIWIL1- piRNAs and PIWIL3-piRNAs, respectively. We identified 101, 89, 74, 55, and 61 piRNA clusters in testis PIWIL1-piRNAs, ovary PIWIL1-piRNAs, MII oocyte PIWIL1-piRNAs, 2-cell PIWIL1-piRNAs, and oocyte PIWIL3-piRNAs, respectively. In the testis, both unidirectional clusters, in which piRNAs map to only one strand, and bidirectional clusters, in which the polarity of piRNA production switches between plus and minus strands, were identified, although unidirectional clusters were dominant (Figure 8A and Figure S7A). We found that the majority of the piRNA clusters identified in female gonads were unidirectional (Figure 8B and Figure S6A).

**Figure 8.**
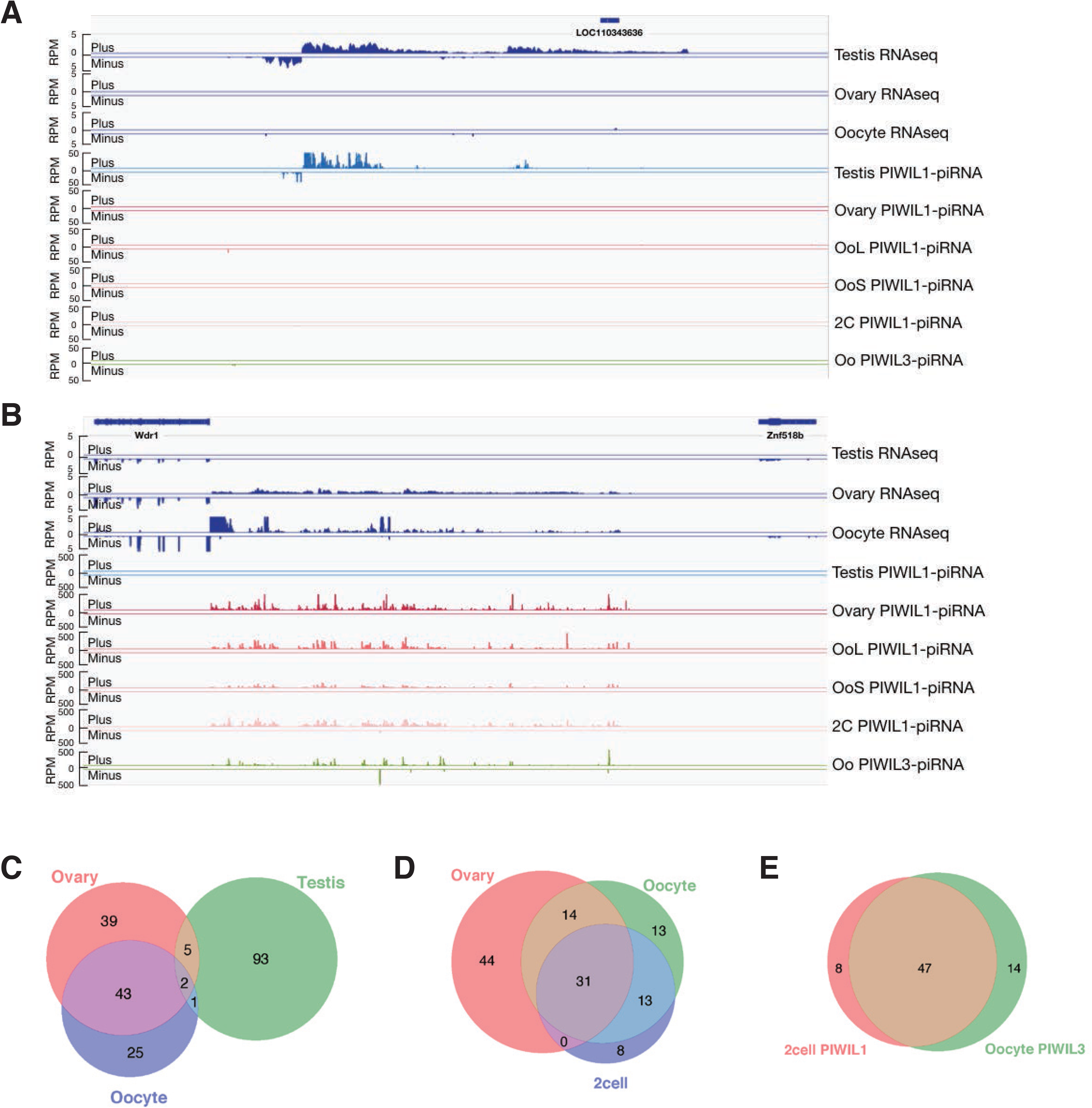
piRNA clusters in hamster male and female gonads. (A) Example of male-specific piRNA cluster in hamster. Distributions of uniquely mapped piRNAs and RNA-seq reads located in piRNA clusters are shown. (B) Example of female-specific piRNA cluster in hamster. Distributions of uniquely mapped piRNAs and RNA-seq reads located in piRNA clusters are shown. (C) The Venn diagram shows the amount of overlap among PIWIL1-piRNA clusters, which are consistent with 28∼30 nt piRNAs in hamster testis, ovary, and MII oocyte. (D) The Venn diagram shows the amount of overlap among PIWIL1-piRNA clusters in hamster ovary, MII oocyte and 2-cell embryo. (E) The Venn diagram shows the amount of overlap among PIWIL1-piRNA clusters in hamster 2-cell embryo and PIWIL3-piRNA clusters in hamster MII oocyte.

Interestingly, approximately 90% of the piRNA clusters corresponding to PIWIL1-associated piRNAs in testis, ovary, and MII oocytes were sex-specific, and only eight piRNA clusters were identified commonly in both male and female gonads (Figure 8C). These results support the idea that transcription of the primary source of piRNA (piRNA clusters) plays a key role in the production of sex-specific piRNAs (Figure 6B and 8C). We next examined overlaps among ovaries, MII oocytes, and 2-cell embryo PIWIL1 piRNA clusters and found that they shared 31 clusters (Figure 8D). PIWIL1-piRNAs in 2-cell embryos shared approximately 85% of the source clusters with those in MII oocytes, while they shared approximately 60% of the source clusters with those in the ovary, which suggests that the genomic regions where piRNAs are derived are altered during oocyte maturation. PIWIL3-associated piRNAs shared 77% of the clusters with PIWIL1-associated piRNAs in MII oocytes. These results are consistent with the findings that PIWIL1- and PIWIL3-associated piRNA populations are likely to share the same 5′ cleaved piRNA precursors (Figure 7A). Finally, we found that piRNAs loaded onto PIWIL3 in MII oocytes shared a large number of piRNA clusters with piRNAs loaded onto PIWIL1 in 2-cell embryos, suggesting that they may have common targets (Figure 8E).

Given that the transcription of piRNA clusters greatly differs in male and female gonads, we analyzed the motif sites surrounding the potential transcription start sites of the testis and ovary piRNA clusters. We first extracted sequences surrounding the transcriptional start sites of unidirectional and bidirectional piRNA clusters, predicted from small RNA-seq and RNA-seq mapping data. We used these sequences and performed motif searches using MEME v.5.1.0, which can discover motifs enriched in the given dataset. The results of the MEME analysis show that the A-Myb binding site is significantly represented in the bidirectional piRNA clusters in the testis transcriptional start site surrounding regions (Figure S6B), where it was under-represented in the ovary transcriptional start site surrounding regions (data not shown). Moreover, we calculated the expression level of A-Myb family homolog genes in hamster testis, ovary, and oocyte using RNA-seq data, and found that A-Myb was especially highly expressed in the testis (TPM = 66.33), whereas it was hardly expressed in the ovary and oocyte (TPM = 2.02 in the ovary and was not detected in MII oocytes, respectively) (Figure S6C), consistent with previous findings that in mice, A-Myb is expressed only in the testis, but not in the female gonads (Li et al., 2013). It has been previously shown that A-Myb regulates transcription of piRNA clusters in mice and roosters (Li et al., 2013). These results suggest that piRNA clusters in hamster testes may also be regulated by the transcription factor A-Myb.

In contrast, we failed to detect clear binding consensus sequences for oocyte-expressing transcriptional factors in the upstream regions of unidirectional piRNA clusters in oocytes. Although we also identified some bidirectional clusters in oocytes (Figure S7A), clear binding consensus sequences for oocyte-expressing transcriptional factors between regions of piRNA clusters could not be detected. This could be due to a lack of information on the exact 5′-ends of the piRNA precursors. Alternatively, the transcription of piRNA clusters in oocytes may differ from the testis piRNA clusters in their modes in which, for example, multiple transcriptional factors, but not a few master transcriptional factors such as A-Myb, may drive the transcription of their loci.

## Discussion

In this study, we generated a new golden hamster genome, which revealed the presence of young and possibly transposition-competent TEs in the genome. This also allowed us to characterize piRNAs in golden hamster oocytes. Intriguingly, the size of PIWIL1-associated piRNAs changes during oocyte maturation. In sharp contrast, PIWIL3 binds to piRNAs only in MII oocytes, and the size of loaded piRNAs is shorter (19 nt).

### PIWI proteins and piRNAs in hamster oocytes and early embryos

The change in the size of PIWIL1-associated piRNAs during oocyte maturation may necessitate unloading of the 3′-ends of the long piRNAs from PIWIL1 to shorten the long PIWIL1-associated piRNAs to short 22–23 nt either by trimming or by direct cleavage to determine their new 3′-ends. Alternatively, PIWIL1-associated short piRNAs may not be the processed products of loaded long piRNAs, but they may be produced by a mechanism similar to that produced by long piRNAs and then loaded onto either newly produced PIWIL1 or PIWIL1, which has unloaded piRNAs. Because it is thought that the size of small guide RNAs is determined by the footprint of small RNA-binding Argonaute/PIWI proteins (Wang et al., 2008), the change in the size of PIWIL1-associated piRNAs implies a change in the structure of the protein that accommodates short piRNAs. The question is how the change in the structure of PIWIL1 is induced to either unload long piRNAs or reload short piRNAs or to conclude the production of short piRNAs, but not long piRNAs, from intermediates, of which the 5′-ends are likely anchored within the MID-PIWI domain of PIWIL1. It is known that the release of the 3′-end of the guide small RNA from the PAZ domain of some bacterial Argonaute proteins occurs during target recognition, which is accompanied by conformational changes in the PAZ domain (Kaya et al., 2016; Sheng et al., 2014). A recent study has also shown that disengagement of the small RNA 3′-end from the PAZ domain occurs during the mammalian Argonaute activity cycle (Baronti et al., 2020). Thus, it is tempting to speculate that conformational changes in PIWIL1, either upon target recognition of long piRNA-loaded PIWIL1 or by hitherto unknown mechanisms, may occur to conclude the production of short piRNAs. In other words, PIWIL1 could switch between states of structural rearrangements to produce two types of piRNAs. In this context, it is interesting that short piRNAs are loaded onto PIWIL3 when the protein is heavily phosphorylated. Indeed, we demonstrated that phosphorylation is required for PIWIL3 to associate with piRNAs. It is known that the loading of small guide RNAs onto Argonaute proteins appears to be affected by phosphorylation, although the underlying mechanisms are poorly understood (Treiber et al., 2019). Phosphorylation could induce changes in the structure of PIWIL3 so that the protein is loaded with processed intermediates of piRNA precursors and the footprint of the protein determines the size of loaded piRNAs. Alternatively, but not mutually exclusive, PIWIL3 may need to be phosphorylated to stably hold piRNAs. Our findings suggest that hamster PIWI proteins in the oocyte can alternate between states (allostery). It will be interesting to model the structural changes in the PIWI protein using published data on structures of PIWI proteins and other Argonaute proteins.

We found that PIWIL3 binds piRNAs only in MII oocytes but appears to be free from piRNAs at other stages of oocyte maturation. There are precedents for piRNA-independent functions of PIWI proteins. Mouse PIWIL1 (MIWI) was found to bind spermatogenic mRNAs directly, without using piRNAs as guides, to form mRNP complexes that stabilize mRNAs essential for spermatogenesis (Vourekas et al., 2012). Recent studies have also shown that human PIWIL1 (HIWI) functions as an oncoprotein via piRNA-independent mechanisms (Li et al., 2020; Shi et al., 2020). Although Argonaute/Piwi proteins tend to be destabilized when small RNAs are not loaded onto them (Elkayam et al., 2012; Haase et al., 2010; Kobayashi et al., 2019; Smibert et al., 2013), these studies suggest that PIWI proteins may be stable in some conditions in the absence of piRNAs to play a role in some cellular functions.

### piRNA-target TEs in the hamster oocyte

In mouse testes, the main target TEs in the piRNA pathway are those of LINE1 and IAP family members. Two distinct piRNA populations are present in mouse testes: pre-pachytene piRNAs are enriched in TE-derived sequences (approximately 80% of the total) (Aravin et al., 2008) and pachytene piRNAs have a higher proportion of unannotated sequences, with diminished contribution from TE-derived sequences (approximately 25%) (Aravin et al., 2006; Girard et al., 2006). These piRNAs guide PIWI proteins to target LINE1 and IAP elements and repress them either by cleaving their transcripts in the cytoplasm or by modifying their chromatin loci in the nucleus (Iwasaki et al., 2015; Ozata et al., 2019). Recent studies also support a model in which TE-derived piRNAs serve as a guide to selectively target young L1 families for *de novo* DNA methylation (Molaro et al., 2014) or H3K9me3 modification (Pezic et al., 2014) in fetal testes. However, the Slicer activity of both MIWI and MILI is still required to maintain TE silencing in the mouse testis after birth, suggesting that continuous cleavage of TE transcripts by the Slicer activity is essential for repressing TEs in mouse testes (Reuter et al., 2011; De Fazio et al., 2011). We found that the contents of PIWIL1- and PIWIL3-associated piRNAs in hamster oocytes are similar to those observed in mouse pachytene piRNAs. However, the major target TEs in the piRNA pathway in hamster oocytes appear as endogenous retroviruses (ERVs), including ERV2 families. Indeed, our results are concordant with the fact that 41.5% of recently active LTR retrotransposons are accounted for by ERV2 families such as ERV2-7_MAu and ERV2-5_MAu. The differences in piRNA targets between testes and oocytes suggest the possibility that the activity of IAP and these ERV2 elements might be distinctively controlled and their young copies in the genome might have jumped in different types of germline cells. Recent studies have shown that ERVs, which constitute a large portion (8∼10%) of mammalian genomes (Mandal and Kazazian, 2008), have a significant impact on shaping transcriptomes and DNA methylation patterns (methylomes) in mammalian oocytes in a species-specific manner (Brind’Amour et al., 2018; Franke et al., 2017). These oocyte transcriptomes and methylomes in mammals are defined by a balance between activation and repression of ERVs. Thus, the piRNA pathway is likely to contribute to the formation of species-specific transcriptomes and methylomes in oocytes. Indeed, it was recently shown that unmethylated IAP promoters in *Miwi*-deficient mouse testes rewire the transcriptome by driving the expression of neighboring genes (Vasiliauskaite et al., 2018). This also implies that spermatogonial dysfunction, observed in *PIWI*-deficient mice, may not simply be due to deregulation of TEs but due to transcriptional rewriting. It will be interesting to see if the lack of PIWIL1 or PIWIL3 leads to dysfunctions in hamster oocytes.

### piRNA clusters in hamster oocytes and early embryos

We found that nearly 80% of piRNA clusters corresponding to PIWIL1-associated piRNAs in testis, ovary, and MII oocytes of hamsters were sex-specific. This is in agreement with previous studies with total small RNA-seq of ovaries, indicating that piRNAs in human and bovine ovaries mostly share common piRNA clusters with piRNAs in testes (Roovers et al., 2015; Williams et al., 2015). However, a recent study showed that only about 30% of human oocyte piRNA clusters overlapped with the human testis-piRNA clusters, proving that testes and oocytes differentially express piRNA clusters (Yang et al., 2019). Our study, together with that of human oocyte piRNAs, suggest a model in which the expression of oocyte and testis piRNA clusters have different functions with distinct targets. Transcriptional factors that regulate their expression are also distinct, though further analysis, including ATAC-seq to define transcription start sites of piRNA precursors, will be needed to identify the transcriptional factors responsible for piRNA clusters in hamster oocytes. In addition, we demonstrated that nearly half of the two populations of PIWIL1-associated piRNAs in oocytes share common clusters and that nearly half of PIWIL3-associated piRNAs in MII oocytes map to clusters from which PIWIL1-associated piRNAs are derived. These results suggest the possibility that a considerable portion of oocyte piRNA cluster transcripts are processed by a common biogenesis pathway but are loaded onto distinct PIWI proteins.

In summary, we have demonstrated that piRNAs are abundantly expressed in hamster oocytes and their size changes during oocyte maturation. Given the recent development of methods to produce genome-edited hamsters (Fan et al., 2014; Hirose et al., 2020), our findings have set the stage for understanding the role that the piRNA pathway plays in mammalian oocytes. Our newly assembled golden hamster genome will also greatly promote the use of golden hamster as a model to understand human disease.

## Materials and methods

### Animals and tissue collection

Ovaries were collected from 4-week-old golden hamsters that were obtained from the Sankyo Labo Service Corporation, Inc. MII oocytes were collected from 8-week-old golden hamsters that were injected with 40 U of equine chorionic gonadotropin (Serotropin; ASUKA Pharmaceutical Co., Ltd., Tokyo, Japan) at estrus. Two-cells were collected from 8-week-old golden hamsters that were injected with equine chorionic gonadotropin at estrus and mated with adult male hamsters.

### 5′ RACE

Total RNA for *PIWIL3* 5′ RACE was extracted from the ovaries of 8-week-old golden hamsters using ISOGEN (Nippon Gene) and RNeasy (Qiagen). 5′ RACE was performed using the SMARTer RACE 5/3 kit (Takara Bio) according to the manufacturer’s instructions. The PCR-amplified PIWIL3 5′ RACE fragments were cloned into the pRACE vector and sequenced.

### Construction of golden hamster PIWI expression vector

To construct the golden hamster PIWI gene expression vectors, PIWI genes were amplified by PCR using gene specific primers and 3-week-old hamster testis and ovary cDNA. The PCR products were digested with *SpeI* and *NotI*, and then cloned into pEF4-3xFlag vector.

### Generation of anti-Hamster PIWIL3 monoclonal antibodies

Anti-PIWI monoclonal antibodies were produced as described previously (Ishizuka et al. 2002; Nishida et al. 2007) with some modifications. Monoclonal mouse antibodies against hamster PIWIL3 were generated by injecting mice with glutathione S-transferase (GST)-Hamster PIWIL3 (20–260 amino acids from the N-terminal where contains repeat region) and then fusing lymph node and spleen cells with the myeloma cell line P3U1 by GenomONE™-CF EX Sendai virus (HVJ) Envelope Cell Fusion Kit (Cosmo Bio) to produce hybridomas. Polyclonal antibodies were selected using ELISA, immunostaining, western blotting, and immunoprecipitation. The hybridoma clone 3E12 used in this study is available for all these applications.

### Western blotting

Western blot analysis was performed as described previously (Miyoshi et al. 2010). One-tenth of protein from ovaries, proteins from 15 oocytes and 2-cell embryos, and one-tenth of protein from the testes were subjected to SDS-PAGE. Culture supernatants of anti-marmoset PIWIL1 (MARWI) (1A5-A7) hybridoma cells (Hirano et al. 2014) and anti-hamster PIWIL3 (3E12) hybridoma cells (1:500) and mouse monoclonal anti-β-tubulin (1:5000, DSHB, E7) were used.

### Immunofluorescence

The ovaries from 8-week-old wild type female golden hamsters were embedded without fixation for frozen sections. Freezing blocks were sliced to 8 µL and treated with an anti-PIWIL1 monoclonal antibody (1A5) or anti-PIWIL3 monoclonal antibody (3E12). An Alexa488-conjugated anti-mouse IgG (Molecular Probes) was used as the secondary antibody. Fluorescence was observed with an IX72 fluorescence microscope (Olympus).

### Immunoprecipitation

Immunoprecipitation details have been described previously (Saito et al. 2009). Briefly, the ovaries were homogenized using TissueLyser II (QIAGEN) and oocytes and 2-cell embryos in which the zona pellucida were eliminated using polyvinyl acetate with acetic tyrode solution and homogenized by pipetting in binding buffer (30 mM HEPES, pH 7.3, 150 mM potassium acetate, 2 mM magnesium acetate, 5 mM dithiothreitol (DTT), 1% Nonidet P-40 (Calbiochem, 492016), 2 mg/mL Leupeptin (Sigma, L9783), 2 mg/mL Pepstatin (Sigma, P5718), and 0.5% Aprotinin (Sigma, A6279)). PIWIL1 and PIWIL3 proteins were immunopurified using anti-MARWI (1A5) and anti-Hamster PIWIL3 (3E12) immobilized on Dynabeads protein G (Life Technologies, 10004D) and anti-Mouse IgG (IBL, 17314) were used as non-immune, negative controls. The reaction mixtures were incubated at 4°C overnight and the beads were rinsed three times with binding buffer.

### β-elimination

Periodate oxidation and β-elimination were performed as described previously (Hirano et al. 2014; Kirino and Mourelatos 2007; Ohara et al. 2007; Saito et al. 2007; Simon et al. 2011). A 100 µL mixture consisting of purified RNAs and 10 mM NaIO4 (Wako, 199-02401) was incubated at 4°C for 40 min in the dark. The oxidized RNAs were then incubated with 1 M Lys-HCl at 45°C for 90 min to achieve β-elimination.

### 32P-labeling

The 5′-end of the RNAs were labeled with [gamma-^32^P] ATP (Perkin Elmer) and T4 polynucleotide kinase (ATP: 5-dephosphopolynucleotide 5’-phosphotransferase, EC 2.7.1.78). The labeled RNAs were cleaned using Micro Bio-Spin™ column 30 (Bio-Rad) and ethanol precipitation. The precipitated RNAs were subjected to SDS-PAGE with 15% urea.

### Summary of genome sequence and assembly of the golden hamster

Raw PacBio reads (Table S1) were assembled into contigs using the FALCON genome assembler, which is widely used for processing long reads (Chin et al., 2016). To correct assembly errors in the FALCON contigs, we applied the Racon module (Vaser et al., 2017) three times. To obtain a chromosome-scale genome assembly, we aligned all contigs to the 22 golden hamster chromosomes (MesAur1.0_HiC, DNA Zoo) (Figure S7). We attempted to fill 3,797 gaps in the reference chromosomes using the FALCON contigs and error-corrected reads. We generated error-corrected reads using the FALCON assembler, which aligned PacBio raw reads to each other, obtained the consensus sequence of aligned reads using multiple alignments, and then output the consensus sequences as error-corrected reads. To fill unsettled gaps, we aligned error-corrected reads to the gaps using the minimap2 program (Li, 2018) and manually inspected the results to determine whether the gaps were spanned by more than one error-corrected read (Table S2). Finally, we polished the draft assembly using the PacBio raw reads and Arrow program.

### PacBio read assembly

We used the FALCON genome assembler version 2018.08.08-21.41-py2.7-ucs4-beta (Chin et al. 2016) with default parameter settings to assemble PacBio reads. To correct errors in the assembly, we applied the RACON module (version 1.4.10; Vaser et al. 2017) three times with default parameter settings.

### Genome assembly alignment

We aligned all contigs in the assemblies to the golden hamster reference assembly (DNA Zoo release MesAur1.0_HiC.fasta.gz) using MUMmer 4.0.0beta2 software (Marçais et al. 2018) with the nucmer program, and the following parameters: mum min cluster = 100, max gap = 300. We also used the minimap2 version 2.13 (Li, 2018) with default parameter settings.

### Effects of the Arrow program on the draft genome

We polished the draft genome using the Arrow program (version 2.3.3; https://github.com/PacificBiosciences/GenomicConsensus) with default parameter settings. We used the QUAST 5.0.2 tool (Gurevich et al. 2013) to calculate mismatch and indel ratios for our golden hamster genome with respect to the DNA Zoo Hi-C genome, both before and after genome polishing (Table S3).

### Gene lift over from N2 to VC2010

We lifted gene structures and other genome annotations from the golden hamster reference assembly (MesAur1.0 release 100) to our golden hamster assembly. Such cross-assembly mapping typically requires an annotation file in a standard format (e.g., GFF3; https://github.com/The-Sequence-Ontology/Specifications/blob/master/gff3.md), chain alignment (Kent et al. 2003), and a program capable of mapping annotations based on chain alignment (e.g., liftOver) (Speir et al. 2016). The reference genome sequence was downloaded (ftp://ftp.ensembl.org/pub/release-100/fasta/mesocricetus_auratus/dna/Mesocricetus_auratus.MesAur1.0.dna.toplevel.fa.gz) as were the annotations of canonical golden hamster genes (ftp://ftp.ensembl.org/pub/release-100/gff3/mesocricetus_auratus/Mesocricetus_auratus.MesAur1.0.100.gff3.gz). Both the genome sequence and its annotations were obtained from the Ensembl database (release-100). To chain-align our golden hamster assembly (as the query) to the reference assembly (as the target), we used methods almost identical to those described by the University of California Santa Cruz (UCSC) for same-species genomic chain alignment (http://genomewiki.ucsc.edu/index.php/DoSameSpeciesLiftOver.pl). The liftOver protocol required several utility programs from UCSC, some of which were downloaded as precompiled binaries (http://hgdownload.cse.ucsc.edu/admin/exe/linux.x86_64). To map genome annotations, we used liftOver with the parameter -gff -minMatch = 0.90.

### Comparative genome analysis

We compared our golden hamster genome with the mouse reference genome (*Mus musculus* GRCm38.p6 release-100 in the Ensembl database; ftp://ftp.ensembl.org/pub/release-100/fasta/mus_musculus/dna/) and the rat reference genome (*Rattus norvegicus* Rnor_6.0 release-100 in the Ensembl database; ftp://ftp.ensembl.org/pub/release-100/fasta/rattus_norvegicus/dna/) using the Murasaki program (ver. 1.6.8) with a seed pattern weight of 76 and a length of 110 (Popendorf et al. 2010).

### Chromosomal evolution in Rodentia

First, we identified synteny blocks that were shared between two or three species, that is, hamster, mouse, and/or rat (details provided in Figure S8). Next, we inferred ancestral karyotypes of Muridae and *Cricetidae* (Figure 3B) by integrating synteny blocks according to maximum parsimony, minimizing the total number of chromosomal rearrangements such as chromosome fusions, chromosome fissions, and translocations. We excluded inversions from chromosomal rearrangements, which were markedly more prevalent than the other rearrangements. We compared our nearly complete golden hamster genome with the mouse (*Mus musculus*) sand rat (*Rattus norvegicus*) reference genomes. Figure 4A shows conserved synteny blocks between the golden hamster, mouse, and rat genomes. We inferred ancestral karyotypes by integrating synteny blocks shared between two or three species according to maximum parsimony to minimize the amount of chromosomal rearrangement. We obtained a high-resolution ancestral karyotype of Muridae, including mice and rats, using the golden hamster genome as the outgroup as well as a species ancestral *Cricetidae* karyotype (Figure 3B).

### Characterization of TEs in the hamster genome

We first used the RepeatModeler ver. 2.0 (Flynn et al, 2020) coupled with LTR_retriever ver. 2.8 (Ou and Jiang, 2018) for *de novo* identification of repetitive elements in the hamster genome. Among the initial repeat candidates obtained, 282 elements, covering ∼37% of the genome in total, were used for subsequent in-depth characterization. In the accurate identification process, we conducted a BLASTN search and collected 80–100 sequences along with the 6-kbp flanking sequences of each element. The sequences were aligned with MAFFT ver 7.427 (Katoh and Standley, 2013) followed by manual curation, and a refined consensus sequence was constructed to be used for the next round of blastn search. This process was repeated for three rounds at maximum until the consensus sequence reached the 5’ and 3’ termini. We finally constructed 177 consensus sequences at the subfamily level and classified them based on the sequence structure and a comparison with known elements using CENSOR (Jurka et al. 2005) and RepeatMasker ver. 4.1.0. We finally constructed a custom repeat library by adding 177 novel consensus sequences to the original rodent repeat library. RepeatMasker analysis was conducted for genome annotation using the cross-match engine with the sensitive mode (-s). Young (*i.e.*, recently active) complete TEs were selected based on the <5.0 K2P divergence from the consensus sequence and the full-length insertions, although ignoring the lack of sequence homology in up to 50 bp of the 5-terminal of LINEs and 3 bp at the edge of other TEs.

### Small RNA-seq library preparation

PIWIL1 and PIWIL3-associated piRNAs were prepared as described previously (Hirano et al. 2014). PIWI family associated small RNAs were extracted from immunopurified proteins using ISOGEN (Nippon Gene, 311-02501). Libraries were prepared using NEBNext Small RNA Library Sample Prep Set (New England BioLabs (NEB), E7330) with slight modifications. Small RNAs obtained and purified using SPRIselect (Beckman Coulter, B23317) underwent 3 ligation at 16°C for 18 h, then free 3 adaptors were degraded using 5 deadenylase (NEB, M0331) and RecJf (NEB, M0264). The 3′-ligated RNAs underwent 5 adaptor ligation at 25°C for 1 h. The RNAs were reverse-transcribed using SuperScriptIII (Life Technologies, 18080-044) and were PCR-amplified using Q5 Hot Start High-Fidelity DNA Polymerase (NEB, M0493) with 18 cycles.

### Small RNA sequencing and data processing

PIWIL1 and PIWIL3-associated small RNAs were sequenced using MiSeq (Illumina) with three replicates from different samples. A total of 32,478,862 reads were obtained and processed as described previously (Iwasaki et al. 2017). The adaptors were trimmed from the reads using Cutadapt version 2.10 (Martin 2011). The replicates were highly correlated (R^2^ ≧ 0.9), so the reads were merged. Each read was mapped to the reference genome (hamster.sequel.draft-20200302.arrow.fasta) using Bowtie version 1.2.3 (Langmead et al. 2009) with the -v 0 option, which extracts the small RNA reads that were perfectly mapped. Genome mapped reads were selected by size using Seqkit (Shen et al. 2016). The PIWIL1-piRNAs were divided into two groups: Oocyte Short (OoS) group, which included 21–27 nt RNAs, and Oocyte Long (OoL) group with 28–31 nt RNAs. PIWIL3-piRNAs with 18–20 nt were selected for further analysis.

### RNA-seq data processing

The RNA-seq library in oocytes was prepared using the SMART-Seq® Stranded Kit (TaKaRa, 634442). Total RNAs obtained using NucleoSpin® RNA Plus XS (TaKaRa, 740990) from approximately 100 oocytes were fragmented at 85℃ for 6 min. Sheared RNAs were processed under the “Low input category.” PCR1 was performed with 5 cycles, followed by PCR2 with 13 cycles, and the final cleanup was performed once. RNA-seq libraries in the ovary and testis were prepared using the TruSeq Stranded mRNA Sample Prep kit. The libraries were sequenced using HiSeq2000 (Illumina) and the obtained reads were combined, resulting in a total of 79,152,300 pair-end reads for hamster testis and 81,979,150 pair-end reads for hamster ovary, respectively. Reads with trimming adaptors and quality filtering were mapped to the hamster reference genome (hamster.sequel.draft-20200302.arrow.fasta) using hisat2 version 2.2.0 (Kim et al. 2015) with the strandness option (--strandness FR). To calculate transcripts per kilobase million mapped (TPM) as expression levels of genes, we used StringTie version 2.1.3 (Pertea et al. 2015).

### Sequence logo

Sequence logos were generated using the motifStack R package (http://www.bioconductor.org/packages/release/bioc/html/motifStack.html). Small RNA sequences were aligned at the 5′-end, and nucleotide bias was calculated per position.

### Annotation of reads

Annotation of genome mapped reads was determined as described previously (Iwasaki et al., 2017) with some modifications. We examined the overlap between read-mapped genomic regions and feature track data. Each feature data was obtained from RpeatMasker (Smit et al. http://www.repeatmasker.org/) for transposons, repeats, tRNAs, rRNAs and snoRNAs, miRDeep2 version 2.0.1.2 (Friedländer et al. 2012) for miRNAs and UCSC LiftOff pipeline for protein-coding genes. Reads were assigned to a feature when the length of its overlap was longer than 90% of the small RNA. The priority of the feature assignment was defined to avoid any conflict of the assignment. For this study, the priority was in the following order: transposon, repeat, miRNA, rRNA, tRNA, snRNA, snoRNA, and protein-coding genes (exons and introns). Any unassigned regions were regarded as unannotated regions.

### piRNA target TE prediction

Prediction of PIWI-piRNA targets derived from TE regions was determined as described previously (Hirano et al. 2014; Iwasaki et al. 2017) with some modifications. First, we eliminated piRNA reads that mapped to tRNAs or rRNAs. The extracted reads were aligned to consensus sequences of transposons (Rodentia custom library), allowing two mismatches. The alignment was performed using Bowtie because of the large number of obtained sequences.

### piRNA cluster prediction

Prediction of piRNA clusters was performed using proTRAC version 2.4.3 (Rosenkranz et al. 2012) under the following conditions as described previously (Yang et al., 2019): (1) more than 75% of the reads that were appropriate to the length of each piRNA; (2) more than 75% of the reads exhibited the 1 U or 10 A preference; (3) more than 75% of reads were derived from the main strand; and (4) -pimin option with 21, 28, and 18 for oocyte PIWIL1-piRNAs and PIWIL3-piRNAs, respectively.

### Motif search at the transcriptional start site of piRNA clusters

We analyzed the motif sites surrounding the transcription start sites of the testis and ovary piRNA clusters. We first extracted sequences surrounding transcriptional start sites of unidirectional and bidirectional piRNA clusters predicted from small RNA-seq and RNA-seq mapping data. For the detection of bidirectional piRNA clusters, we first determined the number of reads per base in the cluster based on the results of RNA-seq with TopHat version 2.1.1 (Kim et al., 2013). We next detected the direction of each base site and a region in which the same direction was contiguous by more than 200 bp was identified. If (+) or (-) occupies more than 75% of the cluster, the cluster is designated as the direction. If the number of reads was less than five, the direction was the same as the previous base site. The region between the switch of transcriptional direction was extracted along with 200bp upstream and downstream regions, as transcriptional start site of bidirectional clusters. The bidirectional cluster with multiple switching regions identified using these criteria was omitted. For the detection of unidirectional piRNA clusters, we extracted 300 bp upstream and 200 bp downstream of genomic regions where piRNA clusters overlapped with the transcript regions detected from Cufflinks version 2.2.1 (Trapnell et al., 2010). The direction of the sequence strands was the same as in the transcripts. We then used these sequences and performed motif searches using MEME version.5.1.0 (Bailey et al., 2009). Tomtom version 5.1.1 (Bailey et al., 2009) was used for motif comparison.

### Visualization of sequenced reads

To visualize the read density obtained from smRNA-seq snd RNA-seq, we created a BigWig file by using HOMER version 4.11 (Heinz et al. 2010) and displayed the Integrative Genomics Viewer (IGV) version 2.4.1. (Robinson et al., 2011). The normalized expression level of each sample was calculated using reads per million reads (RPM).

### Accession number

The accession number for the deep-sequencing datasets reported in this paper is PRJDB10770 in DDBJ.

## Acknowledgments

We thank all members of the Siomi laboratory, especially Drs. Hirotsugu Ishizu and Akihiko Sakashita, for the discussions and comments on this study. We are also grateful to Yukiteru Ono for assisting the bioinformatics. S. Morishita thanks Drs. Erez Lieberman-Aiden and Olga Dudchenko for making the chromosome-scale DNA-ZOO hamster genome publicly available, Koko Saito and Dr. Chie Owa for PacBio Sequel sequencing, and Dr. Yoichiro Nakatani for stimulating discussion on chromosomal evolution in Rodentia. K.I. was supported by a Grant-in-Aid for JSPS Fellows (18J22025) from the Japan Society for the Promotion of Science (JSPS) and is the Keio University Doctoral Student Grant-in-Aid Program 2018 and 2019. This study was supported by Grants-in-Aid for Scientific Research on Innovative Areas (16H06279) (PAGS) to K.I., J.Y., Y.S., A.T., T.I., and S.M. from JSPS. Y.W.I. was supported by funding from JSPS KAKENHI Grant Numbers 19H05268 and 18H02421. S. Morishita was supported by the Japan Agency for Medical Research and Development (AMED) GRIFIN program. H. Siomi was supported by a Grant-in-Aid for Scientific Research (S) (<u>25221003</u>) from JSPS and is a recipient of funding for the Project for Elucidating and Controlling Mechanisms of Aging and Longevity (<u>1005442</u>) from AMED and the ‘Program of totipotency: from decoding to designing’ from JSPS (<u>19H05753</u>).

## Author Contributions

KI and HH designed and performed most of the experiments with YWI, HM, TH, NMS, MT, KS, and MCS. JY, HI, YS, AT, TI, and SM participated in the *de novo* genome assembly of the golden hamster, and JY, YS, and SM in the chromosomal evolution in Rodentia. HN and TK detected transposable elements. HS conceived the study, and KI, SM, and HS wrote the paper with input from the other authors.

## Supplemental Figures

**Figure S1. Related to Figure 1.**
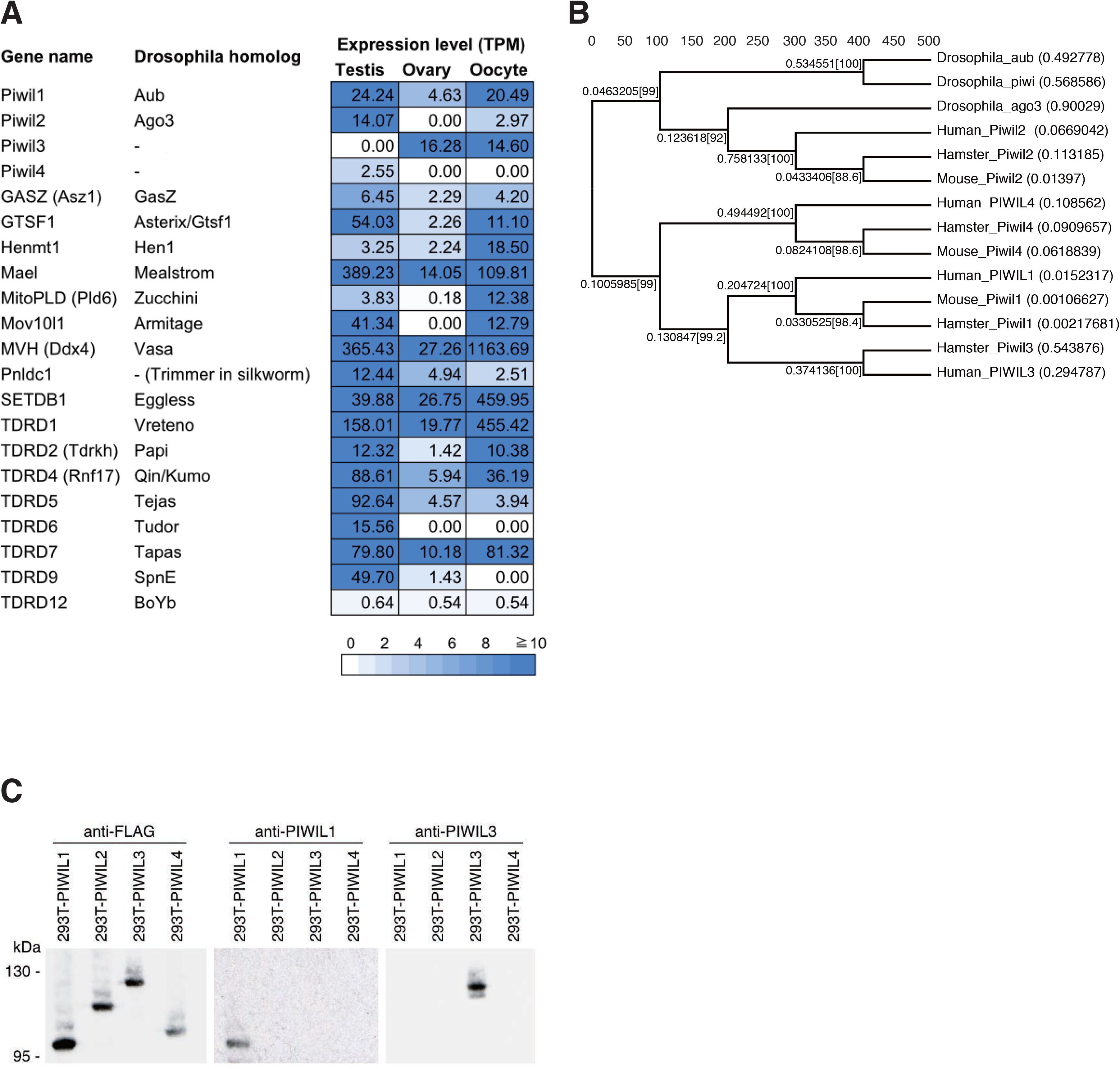
(A) Expression levels of homolog genes in hamster testis, ovary, and MII oocyte, which is known to correspond to the PIWI-piRNA pathway in mouse and *Drosophila.* Expression levels are normalized by transcripts per kilobase million mapped (TPM). (B) A phylogenetic tree of PIWI genes in hamster, human, mouse, and *Drosophila*. Their evolutionary distances were calculated using the maximum likelihood method. (C) Western blotting was performed on 293T cells, which were transfected with *Piwil1* and *Piwil3* genes, respectively. The results show that anti-MARWI (marmoset PIWIL1) and hamster PIWIL3 antibodies specifically recognize their targets.

**Figure S2. Related to Figure 2.**
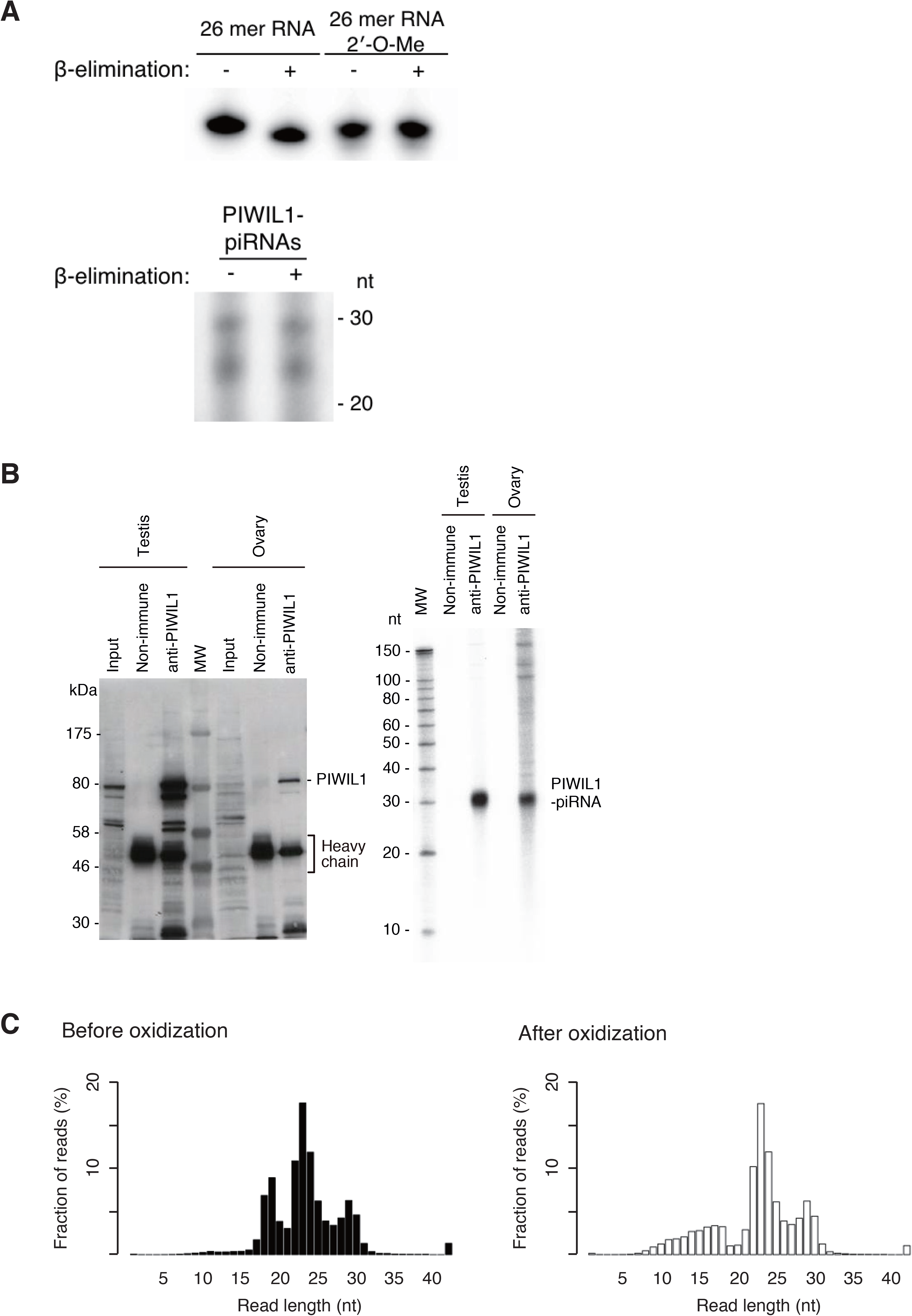
(A) PIWIL1-piRNAs were subjected to periodate oxidation and β-elimination treatment. Synthetic RNAs with or without 2′-*O*-methyl modification were used as controls (upper panel). The unchanged PIWIL1-piRNA signals indicate that the 3′-termini of PIWIL1-piRNAs is 2′-*O*-methylated, as the characteristic of piRNAs (lower panel). (B) Western blotting (left panel) and ^32^P labeling (right panel) were performed from PIWIL1 immunoprecipitates in hamster testis and ovary. Non-immune means negative control, using an anti-mouse IgG antibody. (C) Composition of small RNAs in the hamster oocytes before and after NaIO4 oxidization. The average result of three biological replicates is shown.

**Figure S3. Related to Figure 3.**
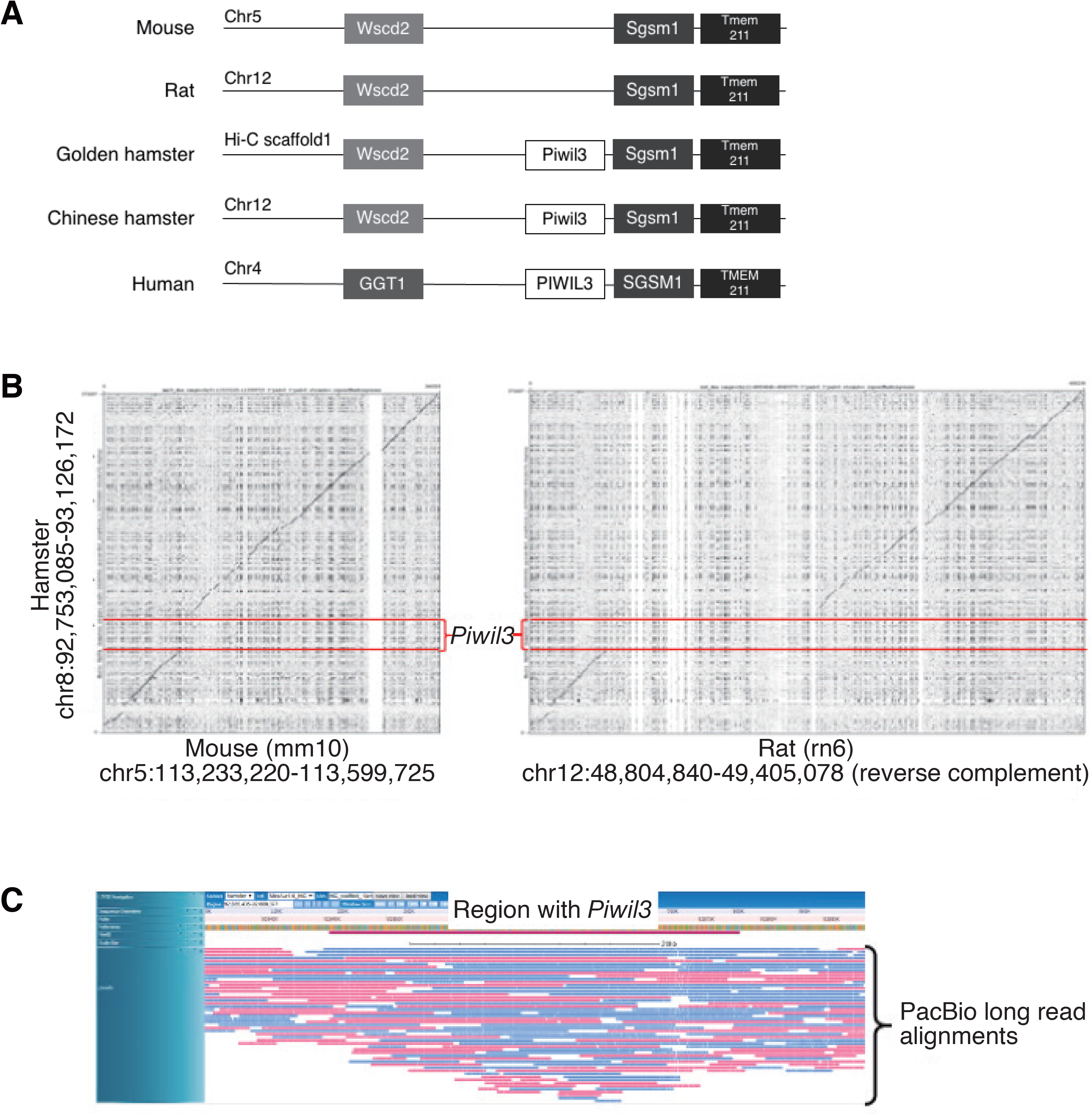
(A) The synteny of the *Piwil3* gene. The synteny of *Wscd2* and *Sgsm1* genes was conserved between hamsters, mice, and rats. Otherwise, a synteny block of *Piwil3*, *Sgsm1*, and *Tmem1* was conserved between hamsters and humans. (B) From our hamster genome and the reference genomes of mouse (mm10) and rat (rn6), we retrieved and compared the three genomic regions between *Wscd2* and *Sgsm1*. The *Piwil3* encoding region is present in the focal region of the hamster genome but is absent in the mouse and rat genomes. The *Piwil3* region is deleted in the mouse genome and is replaced with another sequencing in the rat genome. (C) We aligned to PacBio long reads to the assembled contig with *Piwil3* (the red-colored region). In the lower portion, the respective blue and red colored lines show the alignments of reads in the plus and minus strands. We observed that each base position was covered by an ample number (ten or more) of long reads, and the read coverage was nearly even, thereby confirming the accuracy of the assembled contig.

**Figure S4. Related to Figure 4.**
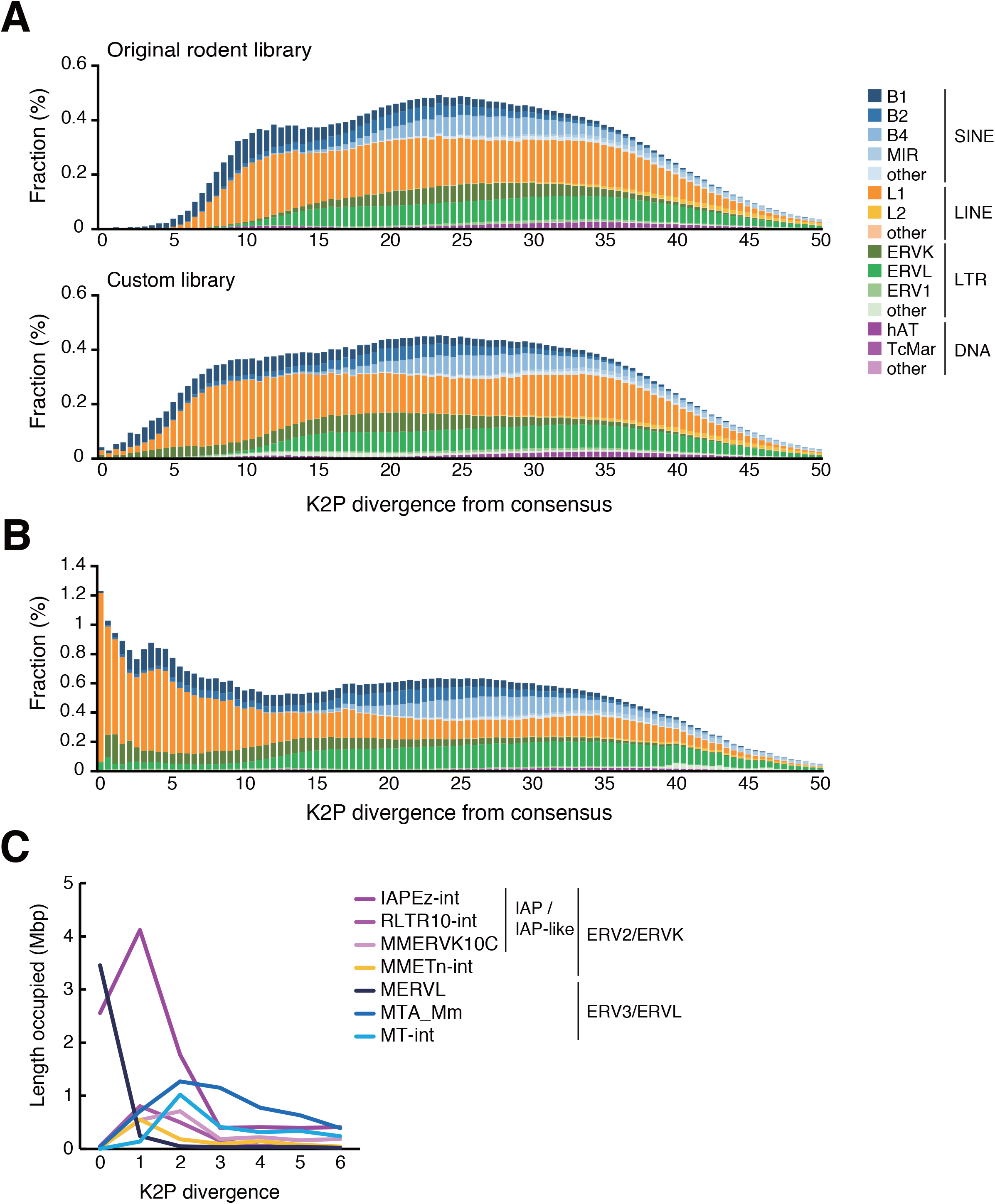
Age distribution of TEs in another hamster assembly (DNA ZOO MesAur1.0_HiC) and mouse genome. The proportion of TEs is shown for 0.5 bins of K2P distance (CpG-corrected) from each consensus sequence. (A) TEs in the MesAur1.0_HiC assembly was analyzed with the original (top) and customized (bottom) repeat libraries. (B) TEs in the mouse genome was analyzed using the mouse repeat library. The color code is the same as in (A). (C) Representative young LTR retrotransposons in the mouse genome.

**Figure S5. Related to Figure 6.**
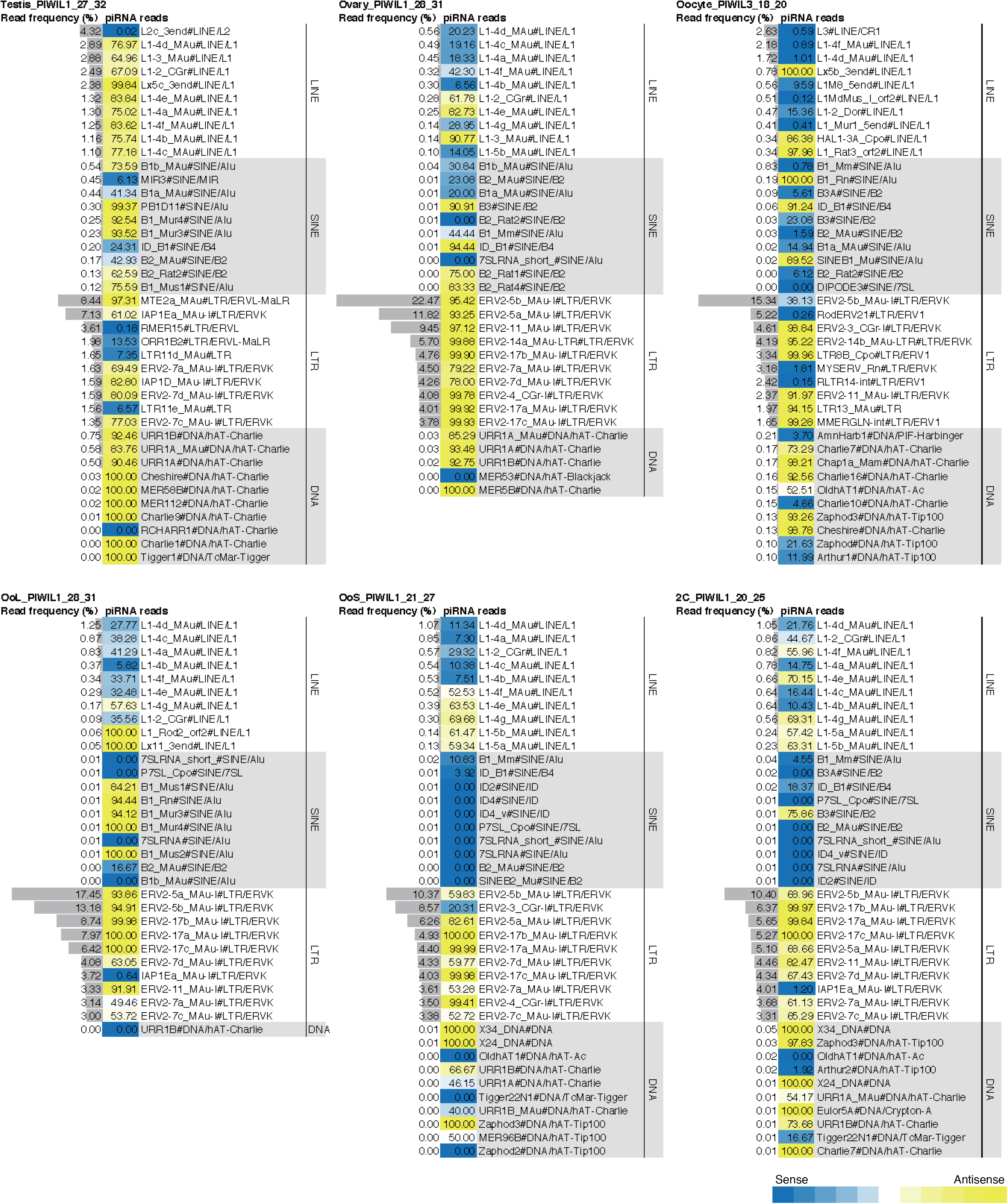
Heat maps show strand bias of transposon-derived piRNAs corresponding to each PIWIs in hamster testis, ovary, MII oocyte and 2-cell embryo. Transposons are grouped into LINEs, SINEs, LTRs, and DNA transposons. Color intensities indicate the degree of strand bias: (blue) sense; (yellow) antisense; (white) unbiased. The frequencies of piRNAs mapped to each TE subfamily over the total TE-mapped piRNAs are shown in the bar graph.

**Figure S6. Related to Figure 8.**
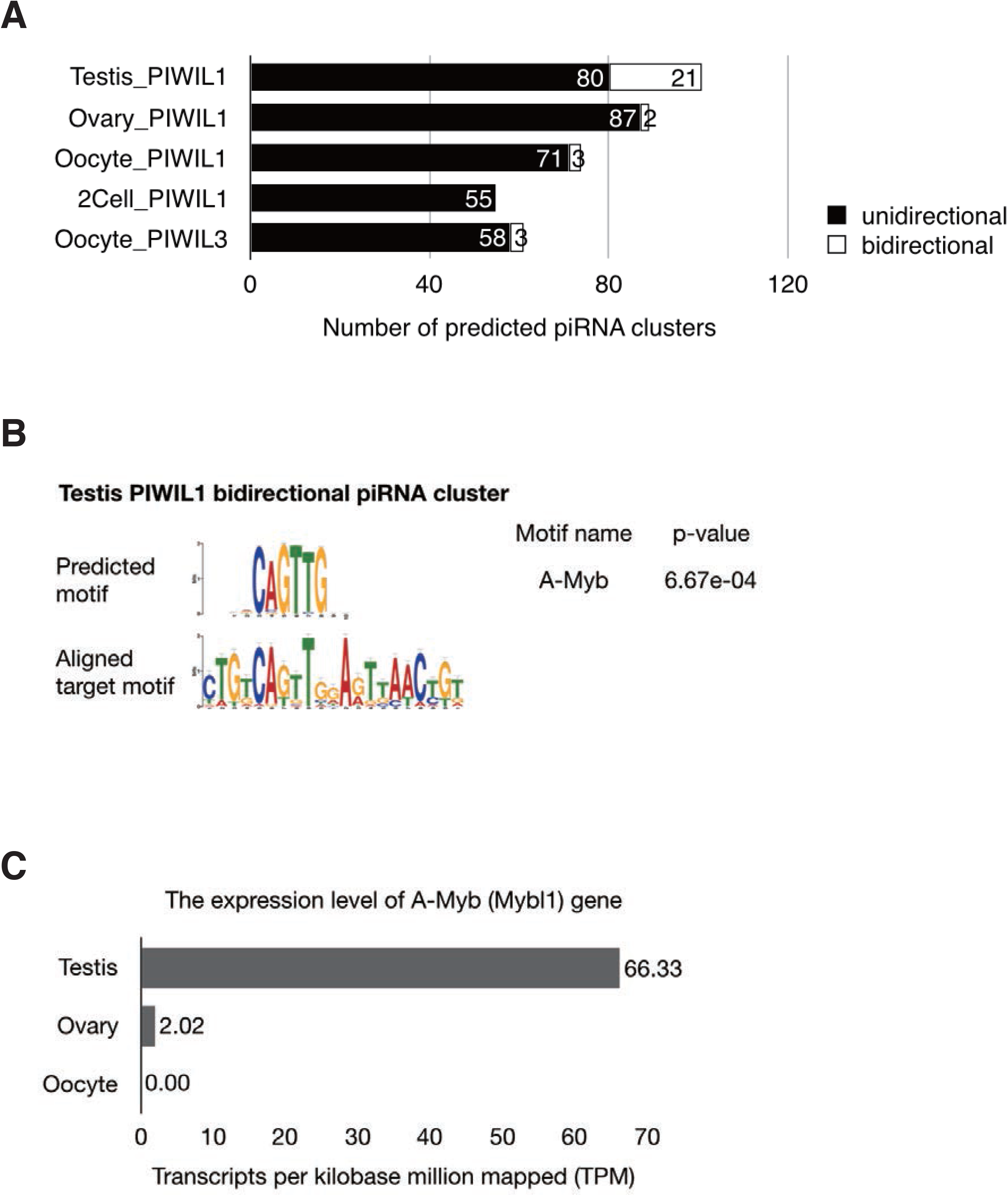
(A) The number of predicted piRNA clusters in hamster testis, ovary, MII oocyte and 2-cell embryo. The black bar shows the unidirectional piRNA clusters and white shows the bidirectional piRNA clutsters, respectively. (B) A motif search was performed for each unidirectional and bidirectional piRNA cluster in the hamster testis using MEME. The results suggest that the A-Myb binding site is significantly represented in the bidirectional piRNA clusters in testis transcriptional start site surrounding regions. (C) Expression levels of transcription factor candidates detected using MEME. The expression pattern of the A-Myb gene in male and female hamsters. RNA-seq data were obtained using Illumina HiSeq2000. The x-axis suggests the normalized expression level (TPM).

**Figure S7.**
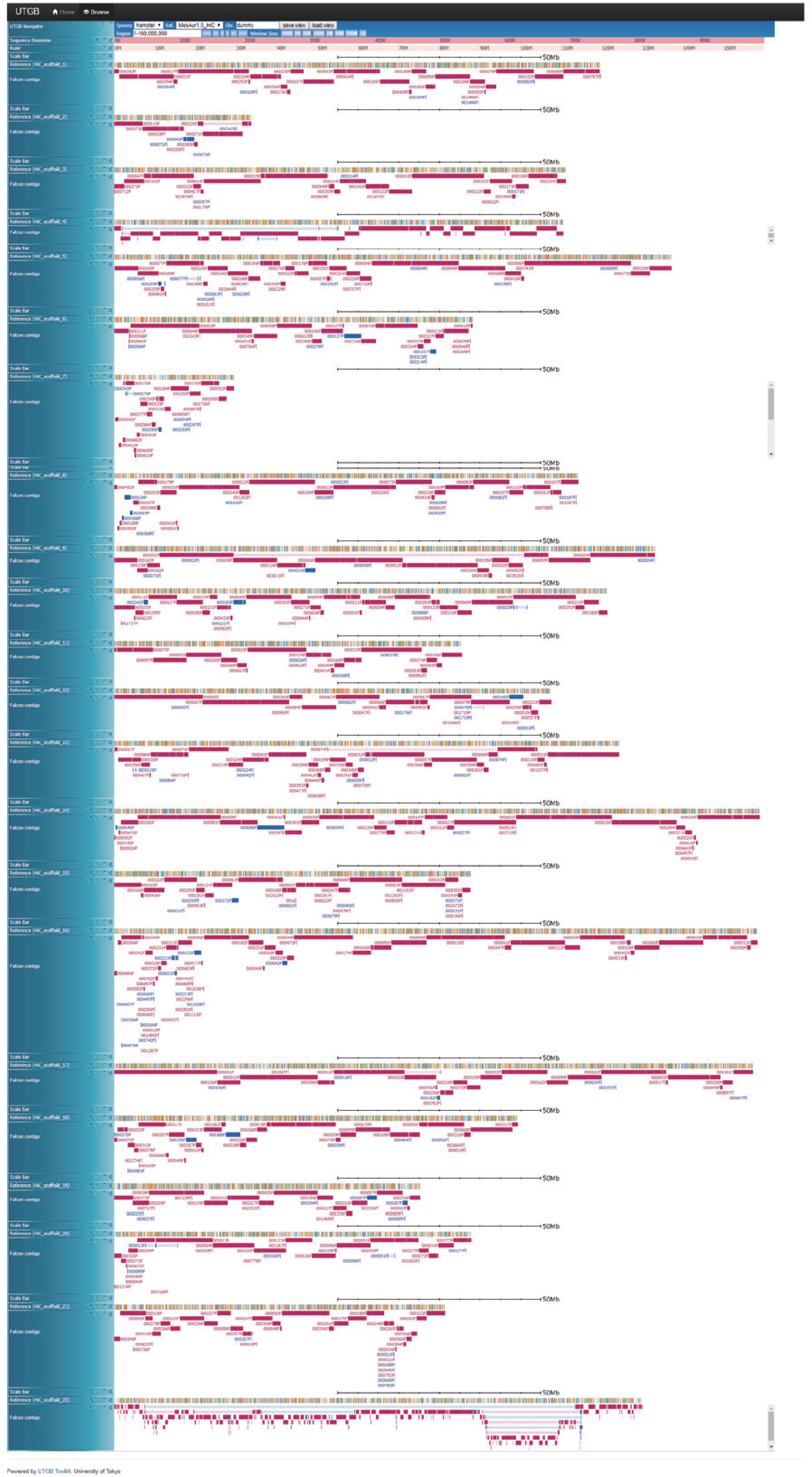
Contigs aligned to the reference assembly. The 22 panels show the alignments of our assembled contigs to the 22 Hi-C scaffolds of the DNA Zoo Hi-C assembly. Thick red and blue lines show alignments of contigs in the plus and minus strands, respectively. Labels beside each contig line indicate the contig identifier.

**Figure S8.**
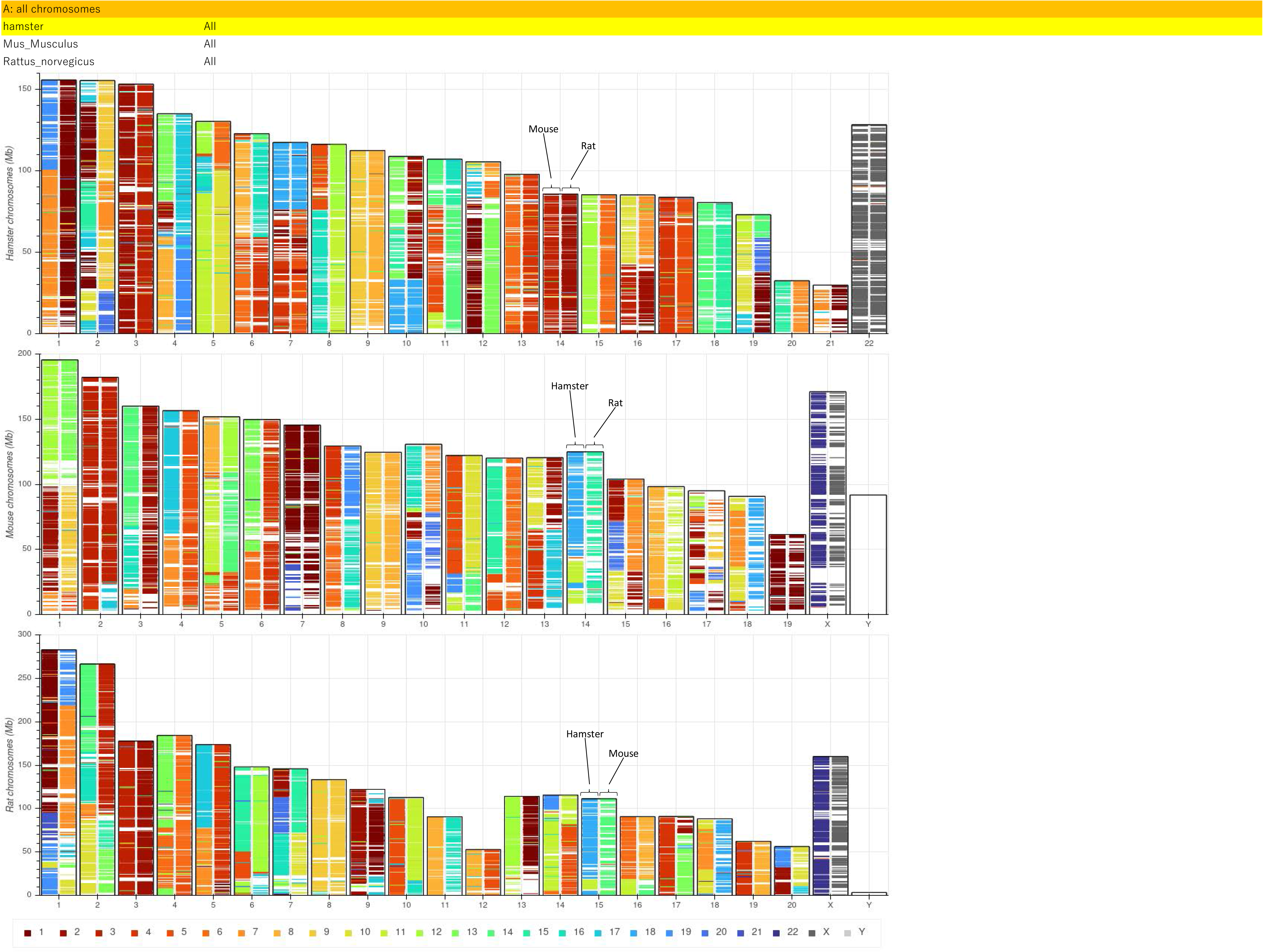
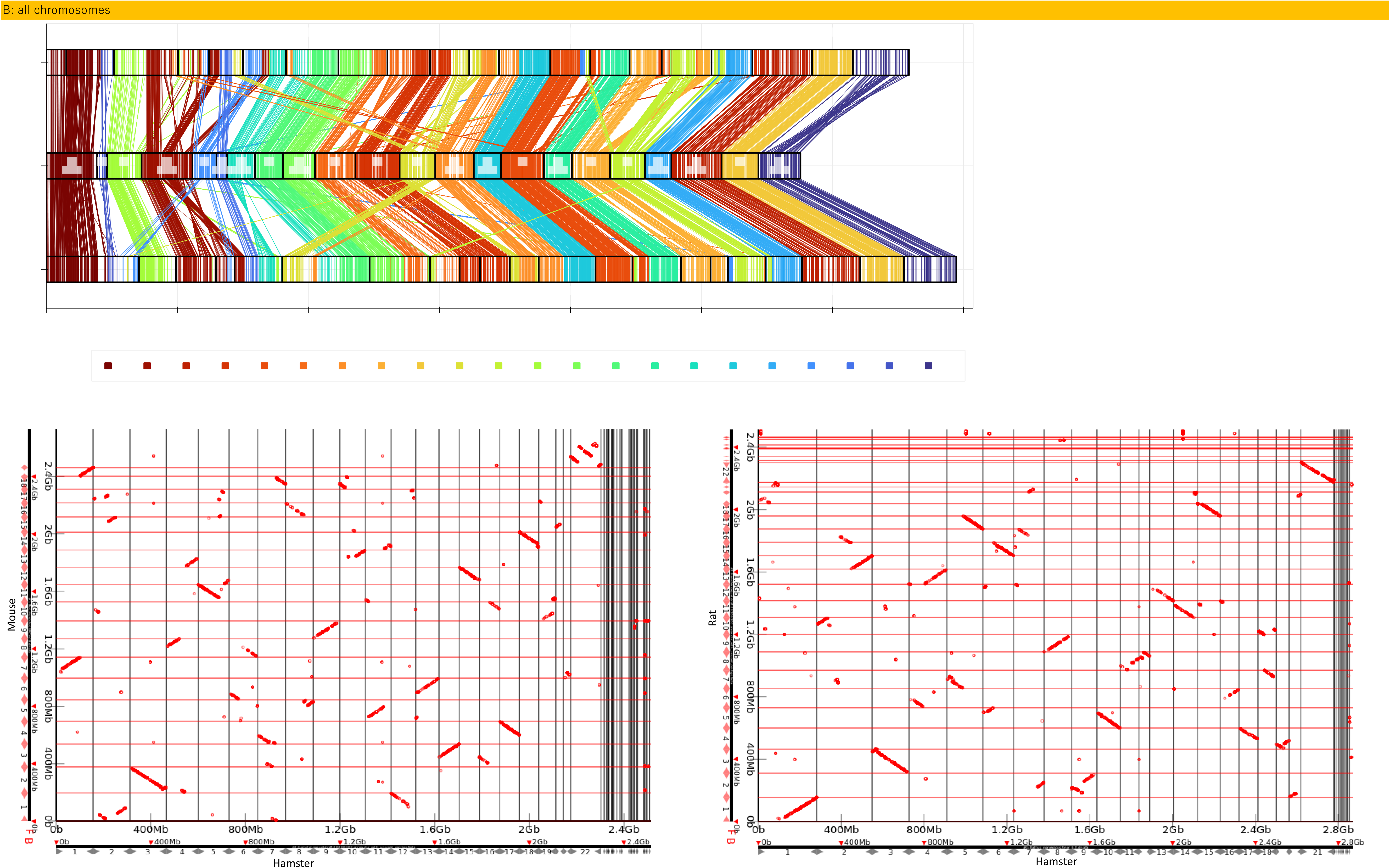
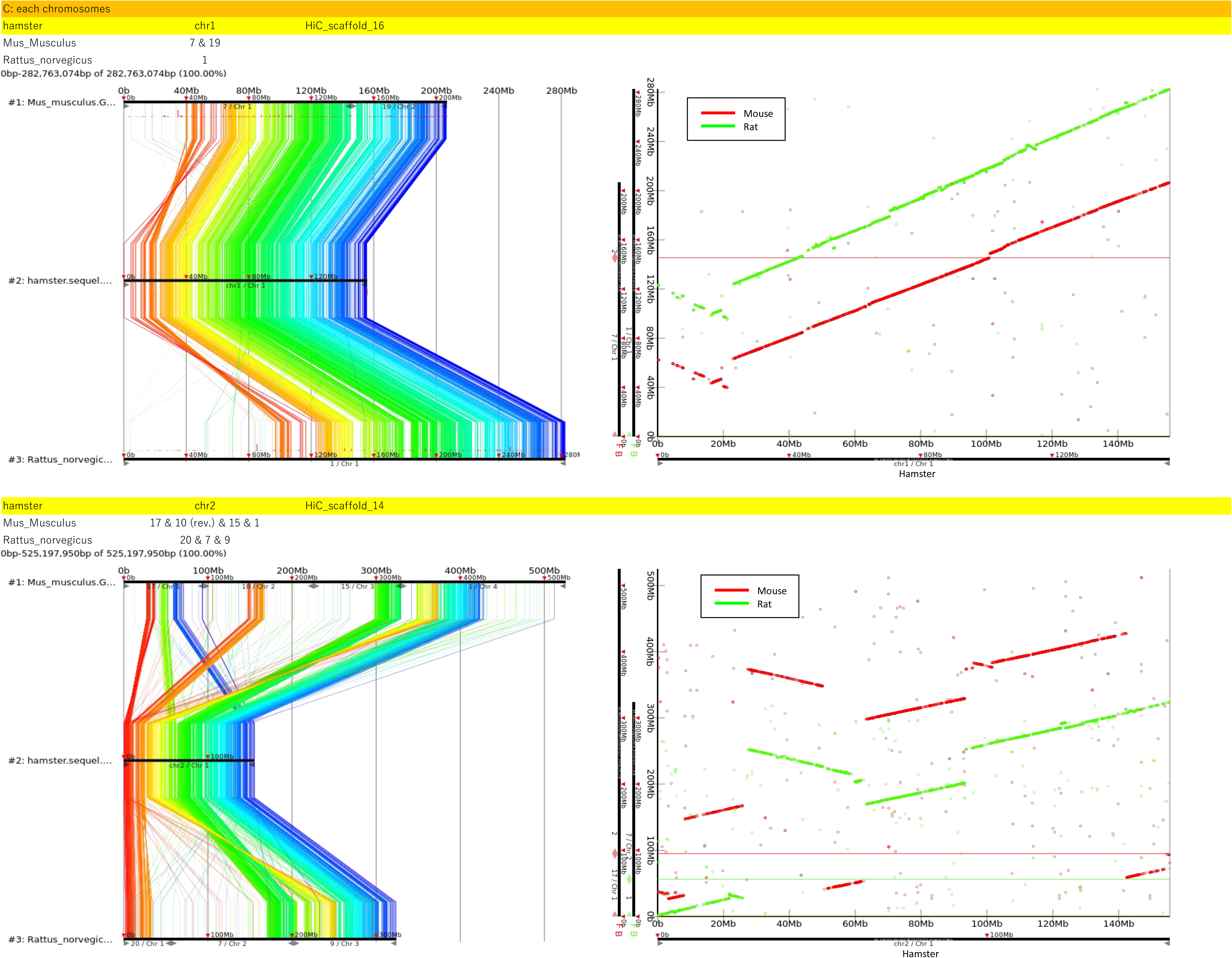
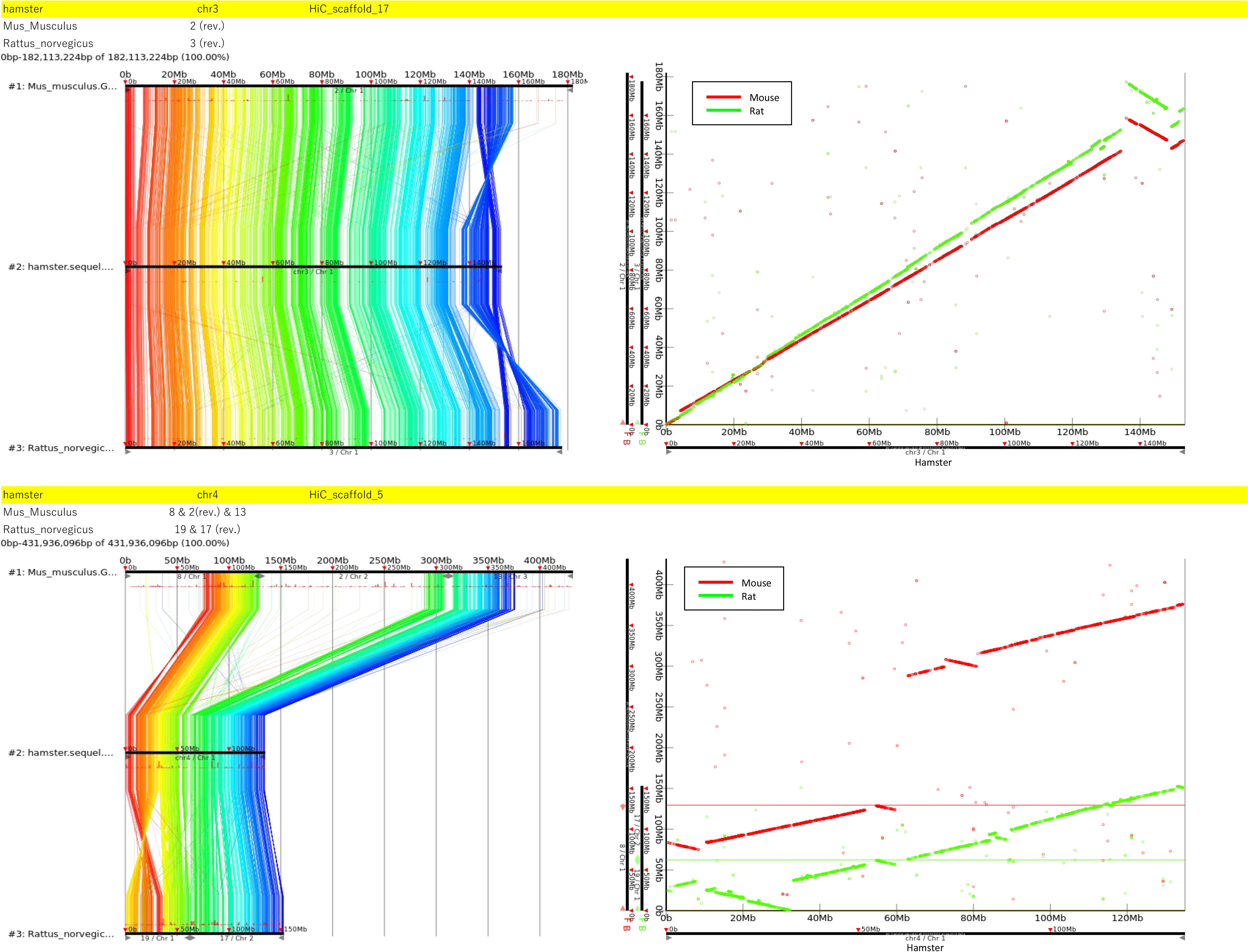
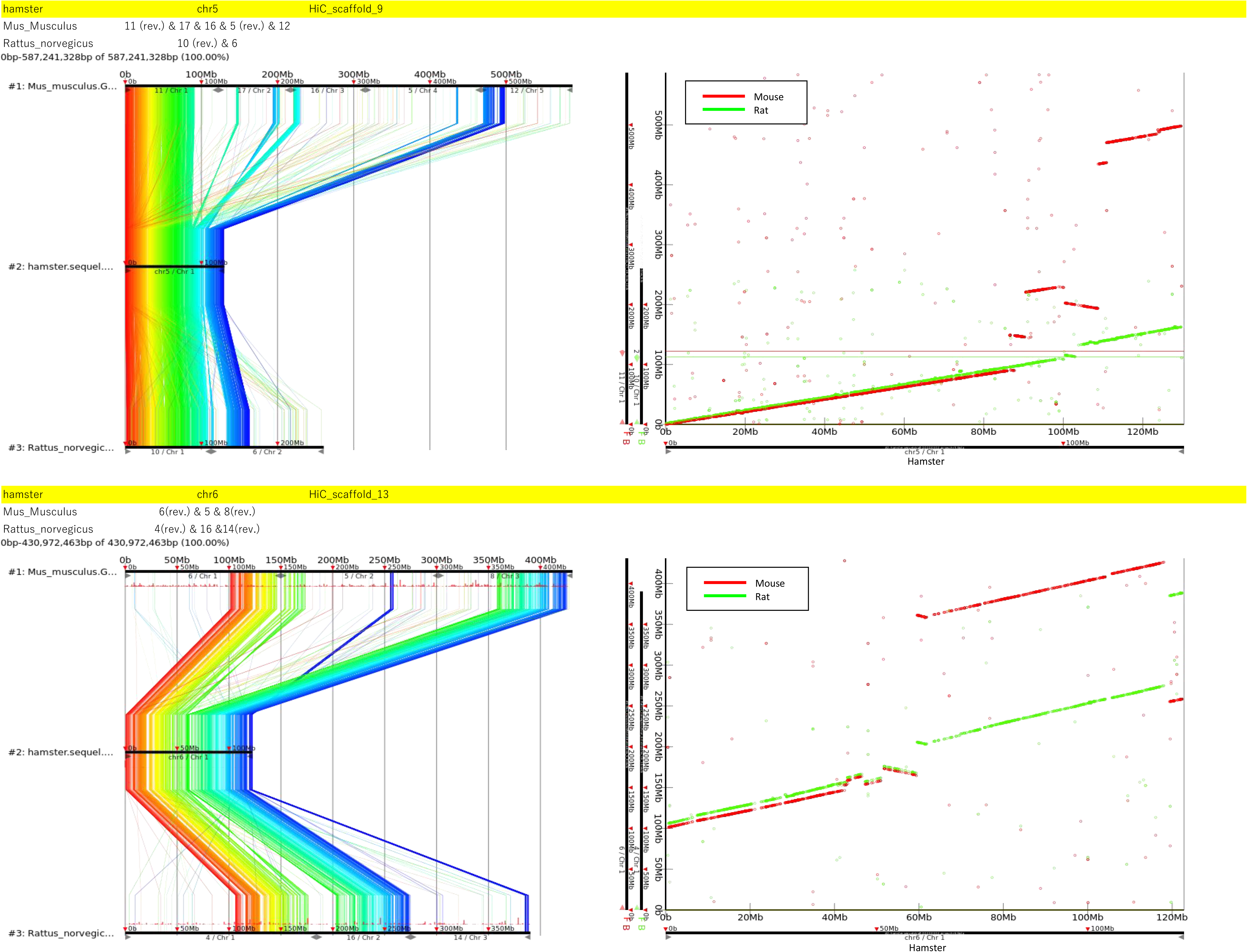
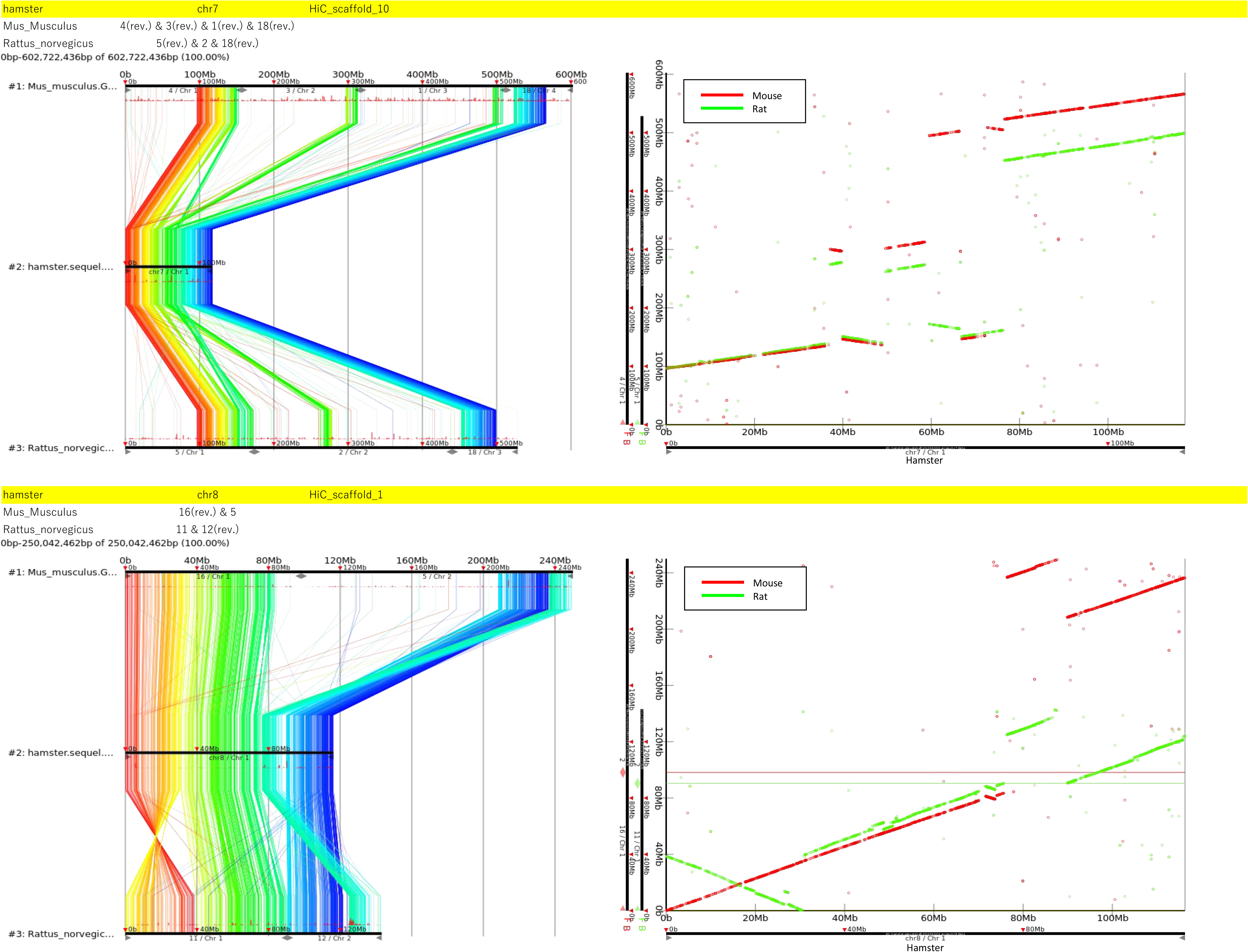
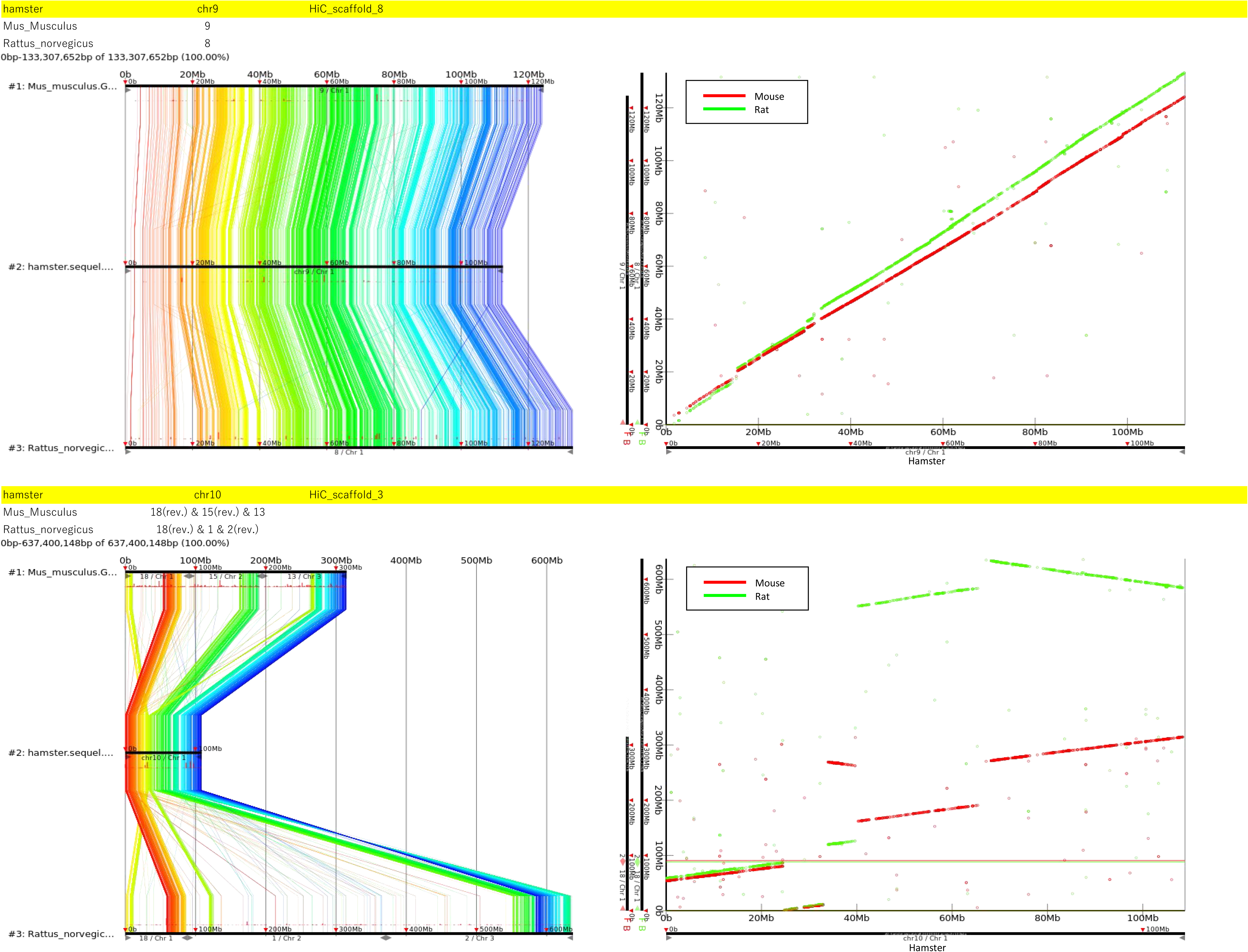
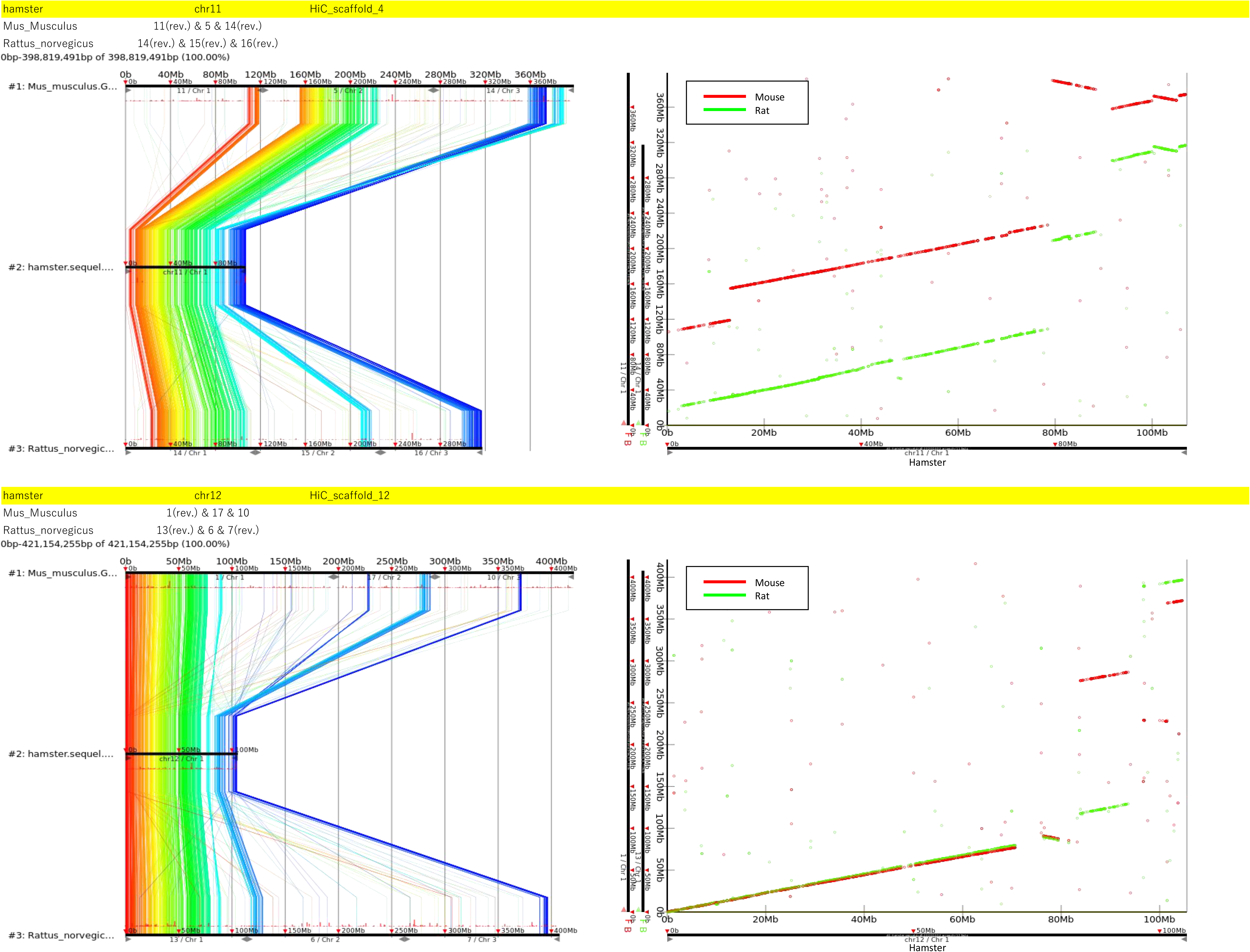
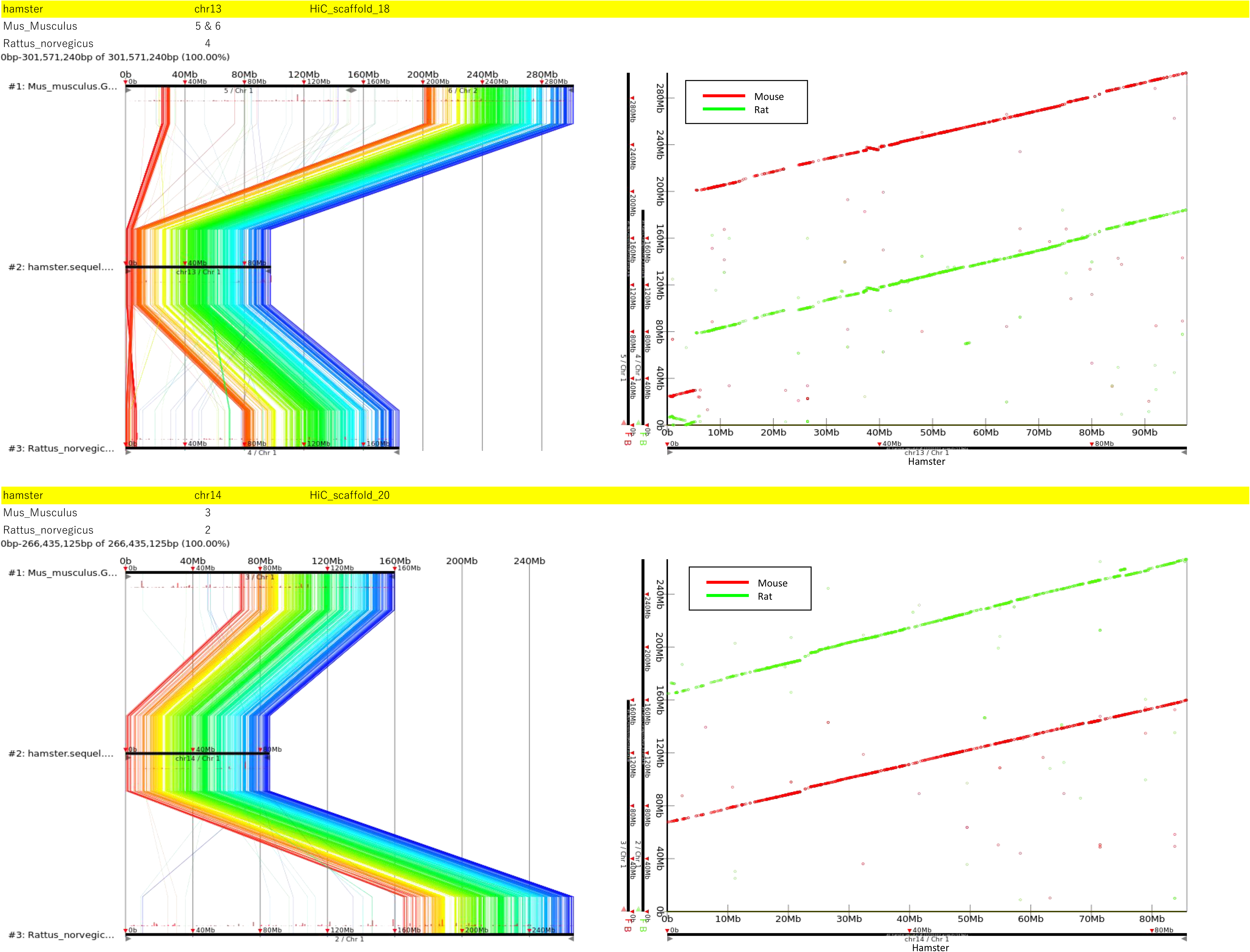
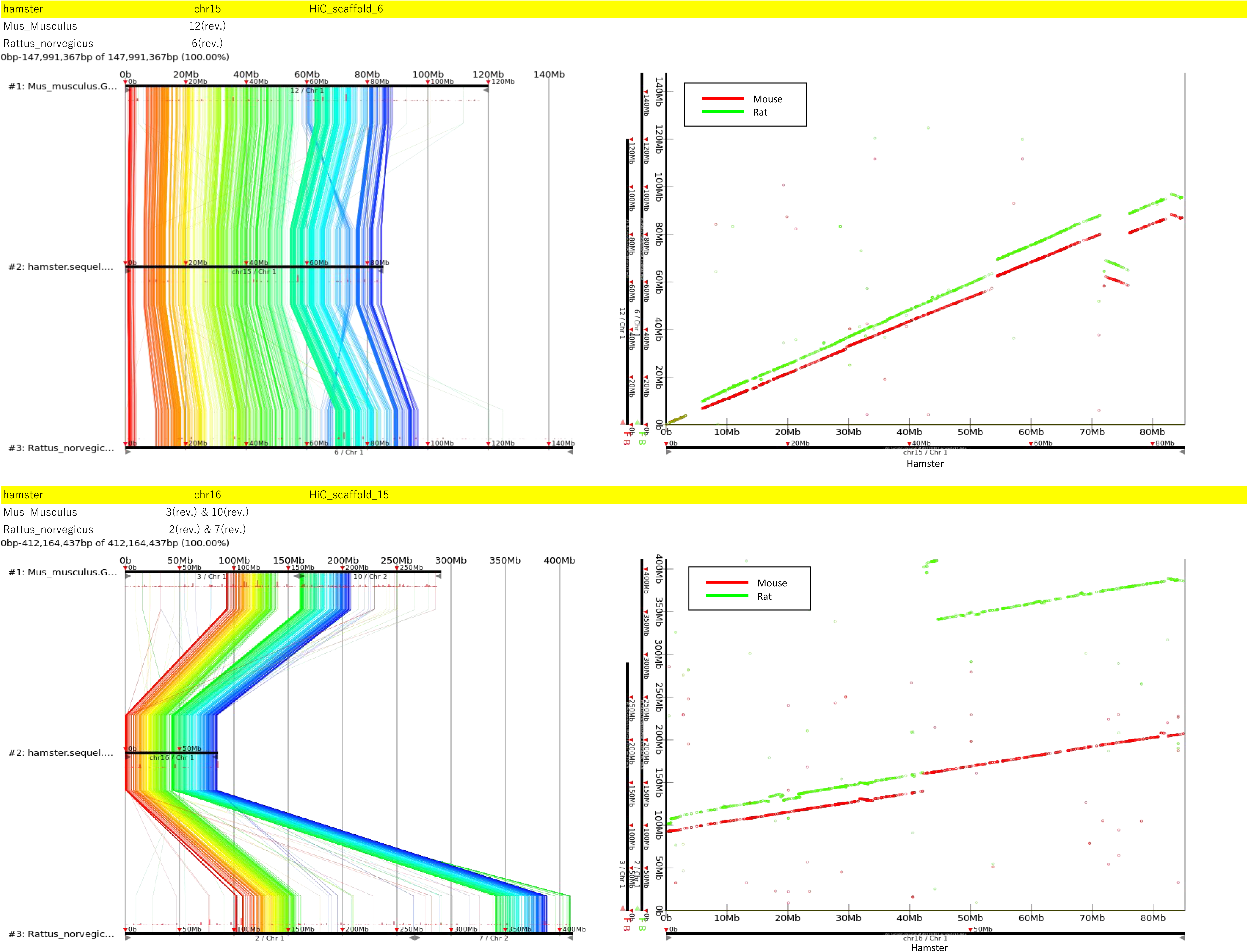
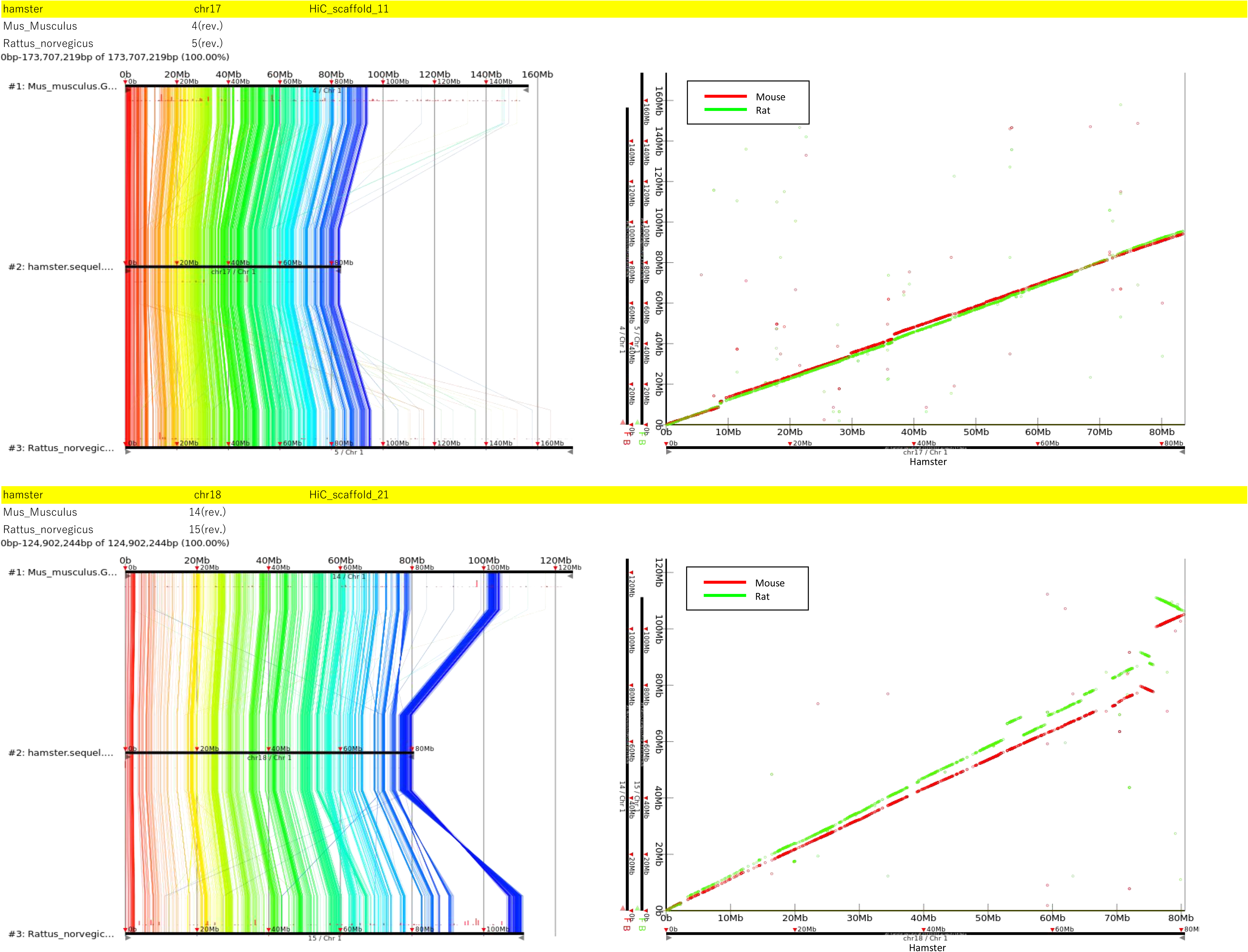
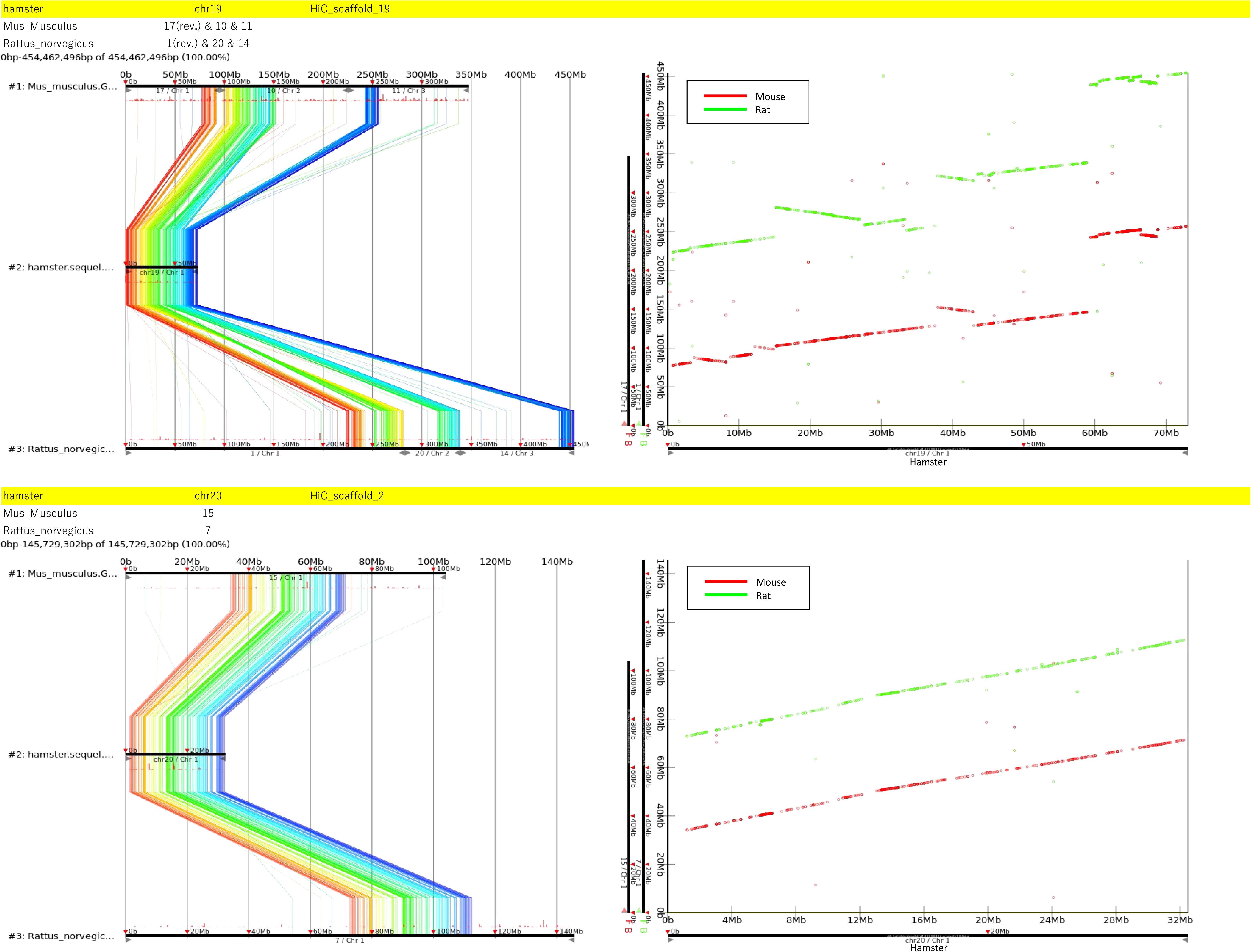
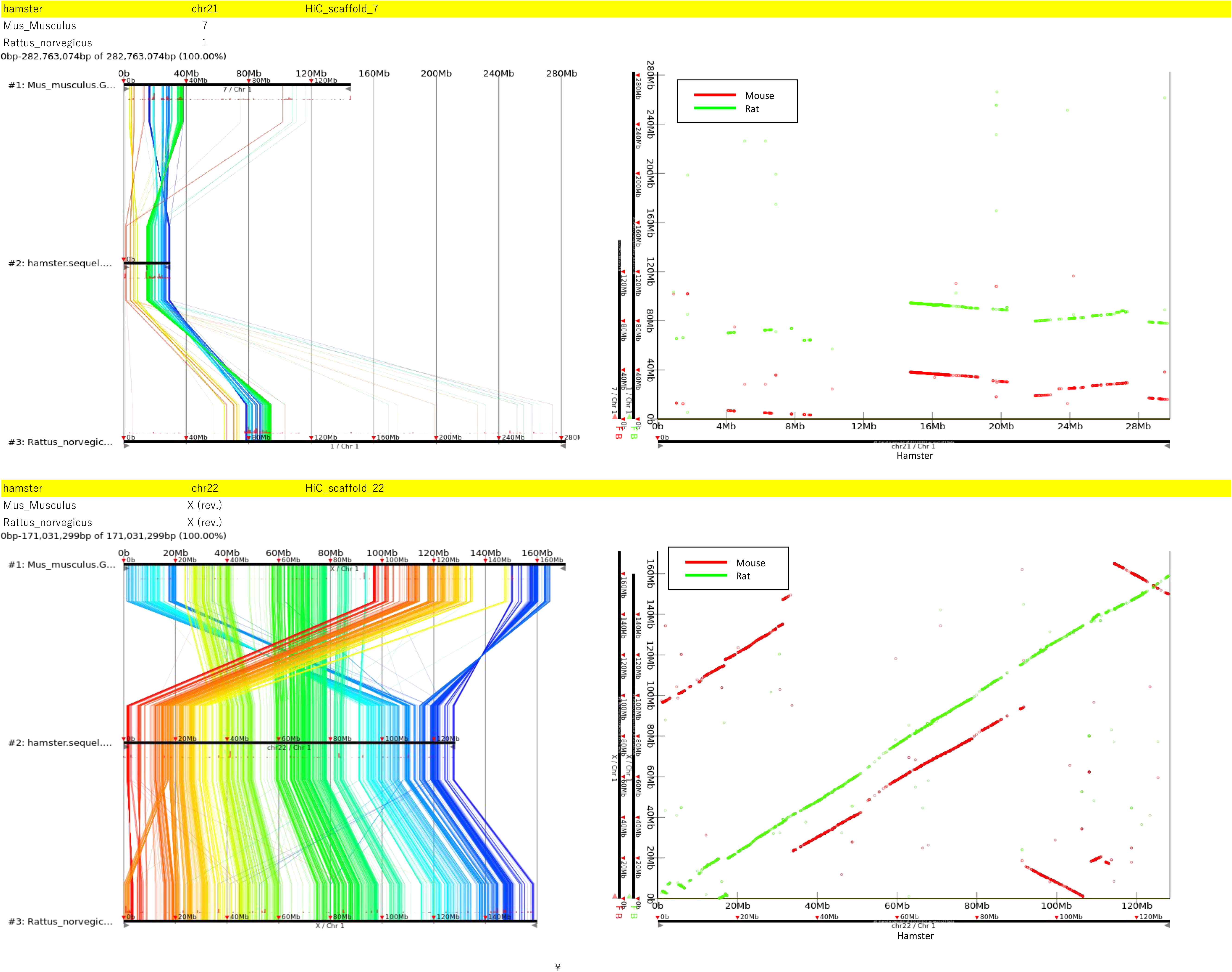
Map of conserved synteny between the hamster, mouse, and rat genomes. (A) Each chromosome in the hamster (top), mouse (middle), and rat (bottom) genomes are associated with a two-column box. The left and right columns show chromosomes in the genomes of mouse and rat (top row), hamster and rat (middle row), and hamster and rat (bottom row), respectively. The same color-coding of chromosomes shown in the color pallet at the bottom was used for the three species. (B) The upper portion is identical to Figure 1B. To facilitate the identification of synteny blocks, dot plots of the hamster (x-axis) and mouse (y-axis) genomes (bottom left), and the hamster (x-axis) and rat (y-axis) genomes (bottom right) are shown. (C) Left panel, color-coded lines showing reciprocally best-matching pairs of positions in the hamster (middle) and mouse (top) or rat chromosomes (bottom) for each of the 22 hamster chromosomes. Right panel: synteny blocks between the hamster genome on the x-axis and the mouse (red) and rat (green) genomes on the y-axis.

**Table S1. Related to Table 1.** Statistics of raw and “polished” long genome sequencing reads from the golden hamster sample using the PacBio Sequel system.

|  | Sequel raw reads |  | polished Sequel reads |  |
| --- | --- | --- | --- | --- |
| Number of reads | 12,586,368 |  | 8,957,271 |  |
| Total Bases | 234,079,417,624 |  | 118,229,107,194 |  |
| Coverage (2.5Gb) | 93.6 |  | 47.3 |  |
| Max length | 225,226 |  | 65,535 |  |
| Min length | 50 |  | 500 |  |
| Average length | 18,598 |  | 13,199 |  |
| N10 | 46,646 | 422,154 | 37,153 | 269,328 |
| N20 | 39,659 | 970,508 | 31,004 | 620,374 |
| N30 | 35,359 | 1,597,485 | 26,988 | 1,029,810 |
| N40 | 32,142 | 2,292,726 | 23,080 | 1,502,938 |
| N50 | 29,513 | 3,053,500 | 19,413 | 2,061,746 |
| N60 | 26,841 | 3,883,122 | 16,115 | 2,730,037 |
| N70 | 22,718 | 4,823,737 | 12,979 | 3,546,302 |
| N80 | 17,327 | 5,996,473 | 9,902 | 4,586,019 |
| N90 | 11,041 | 7,662,185 | 6,659 | 6,028,787 |
| N99 | 2,834 | 10,890,109 | 2,609 | 8,339,269 |

**Table S2. Related to Table 1.**
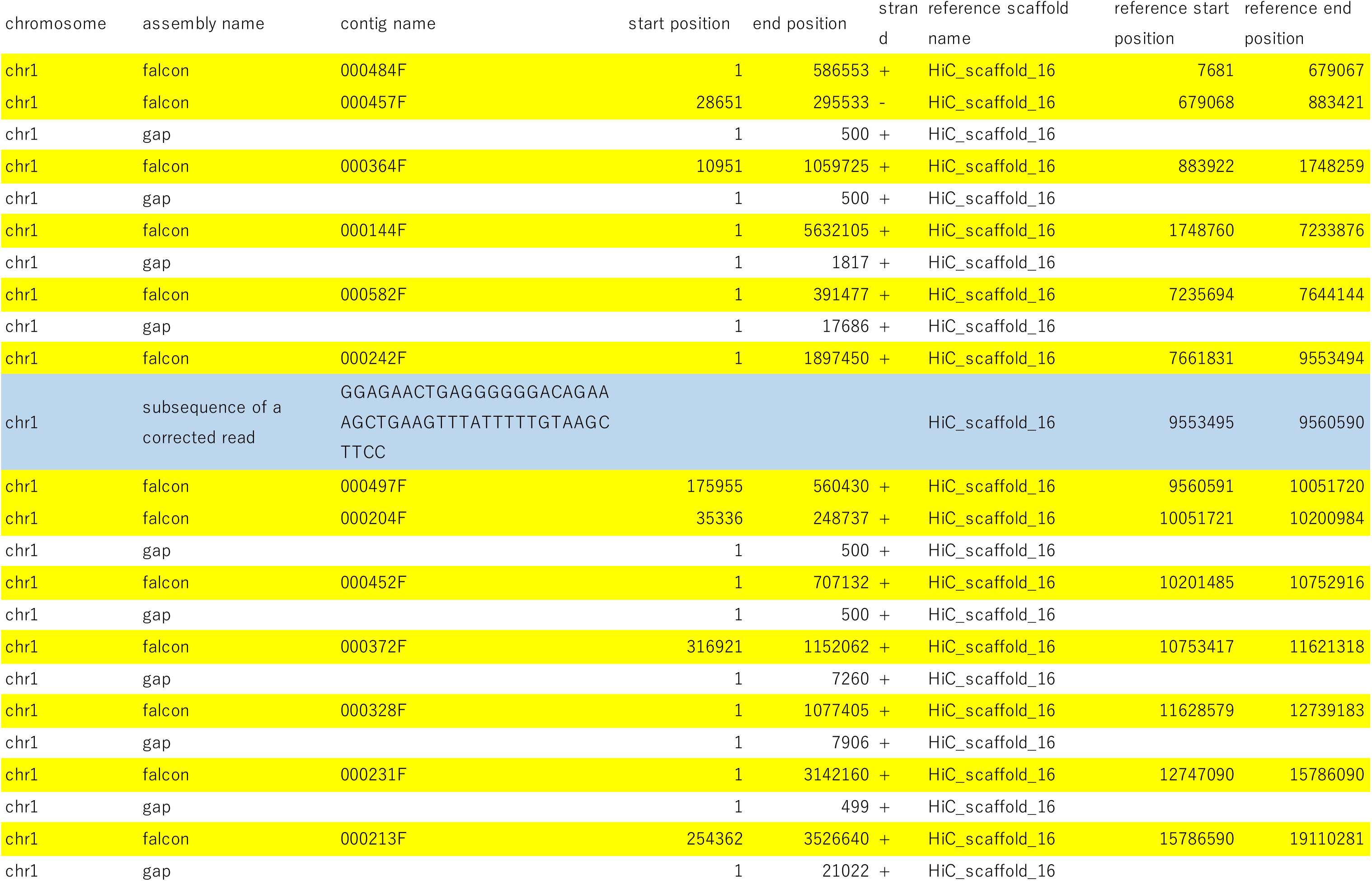
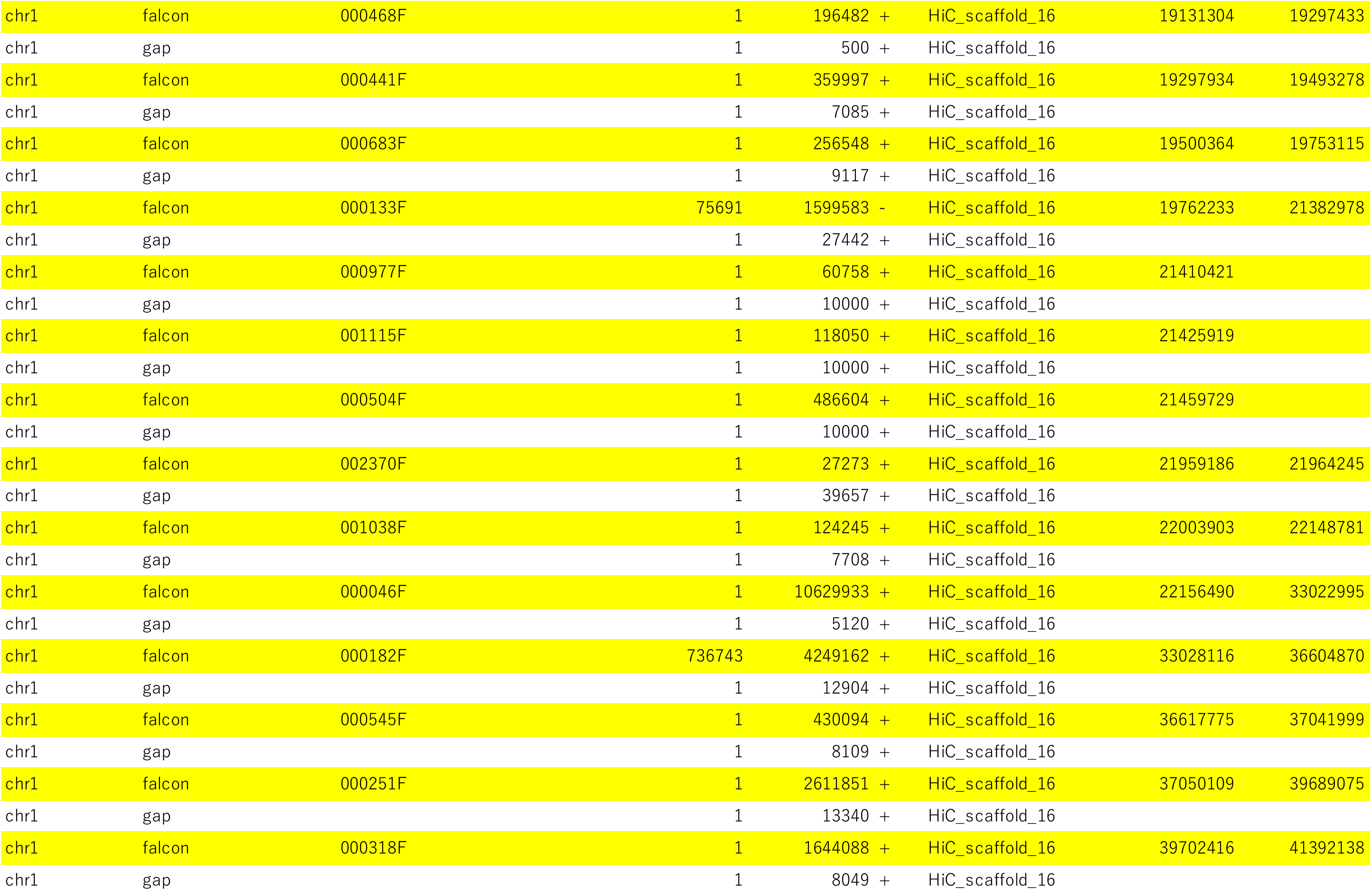
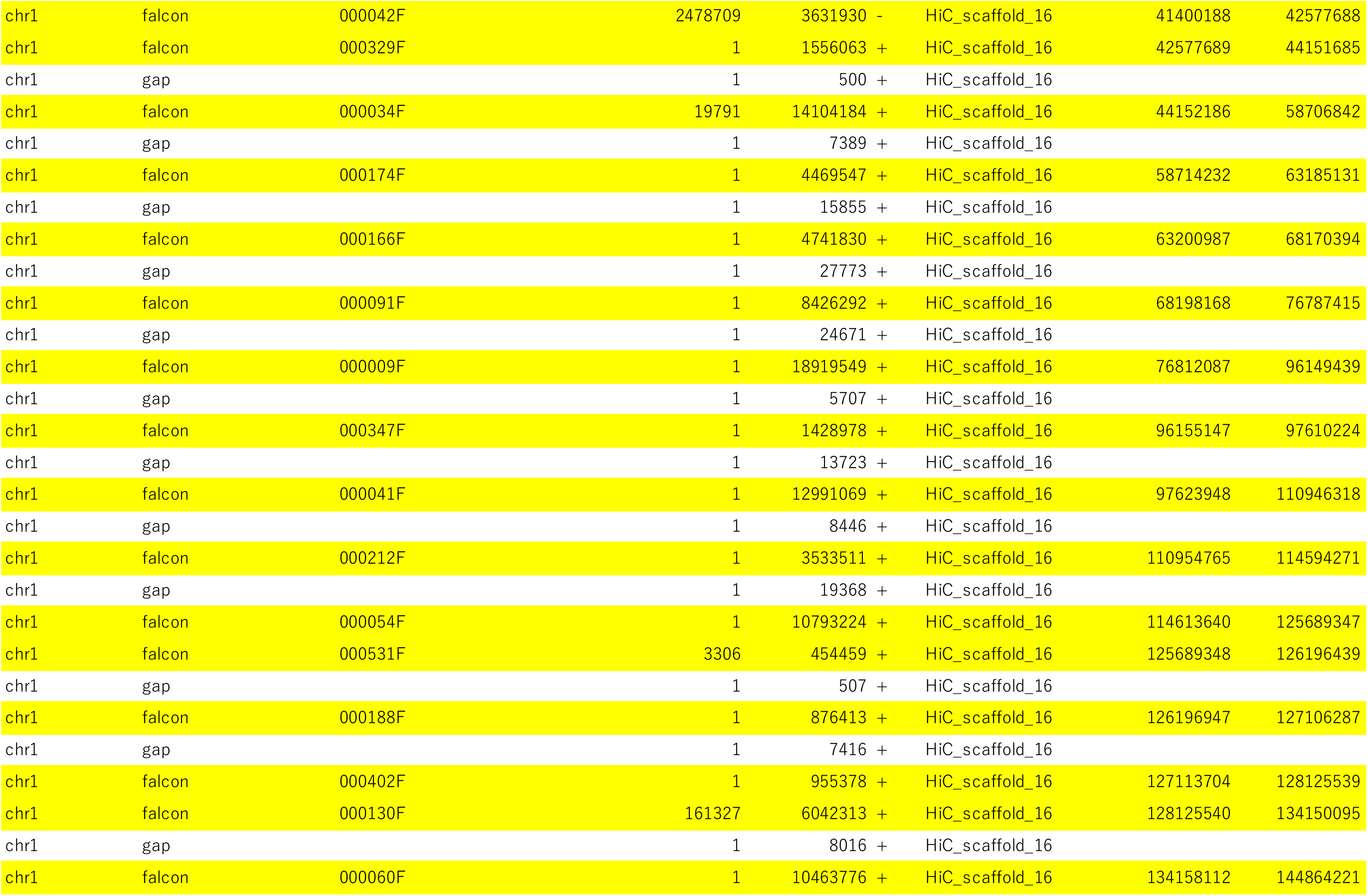
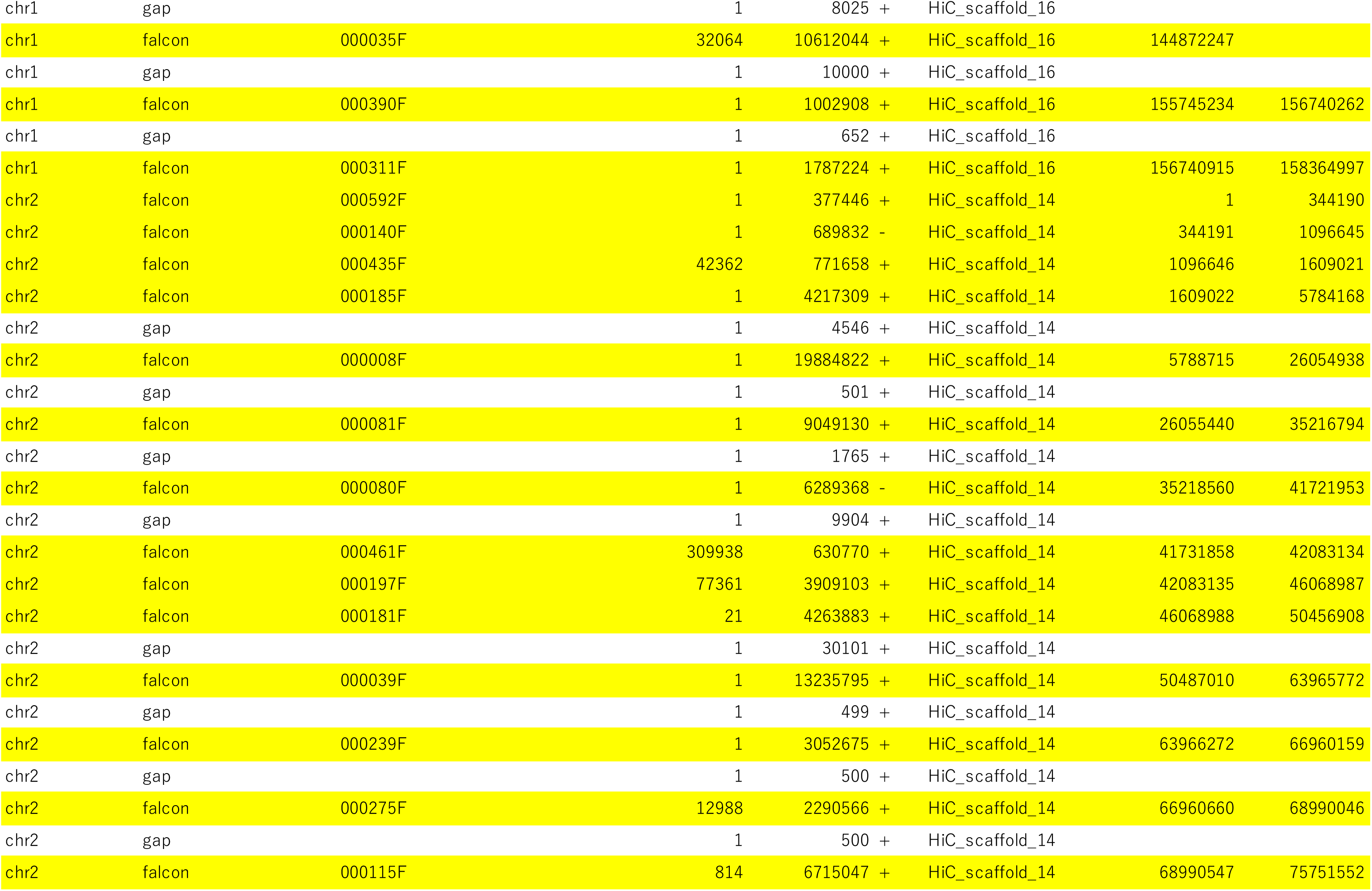
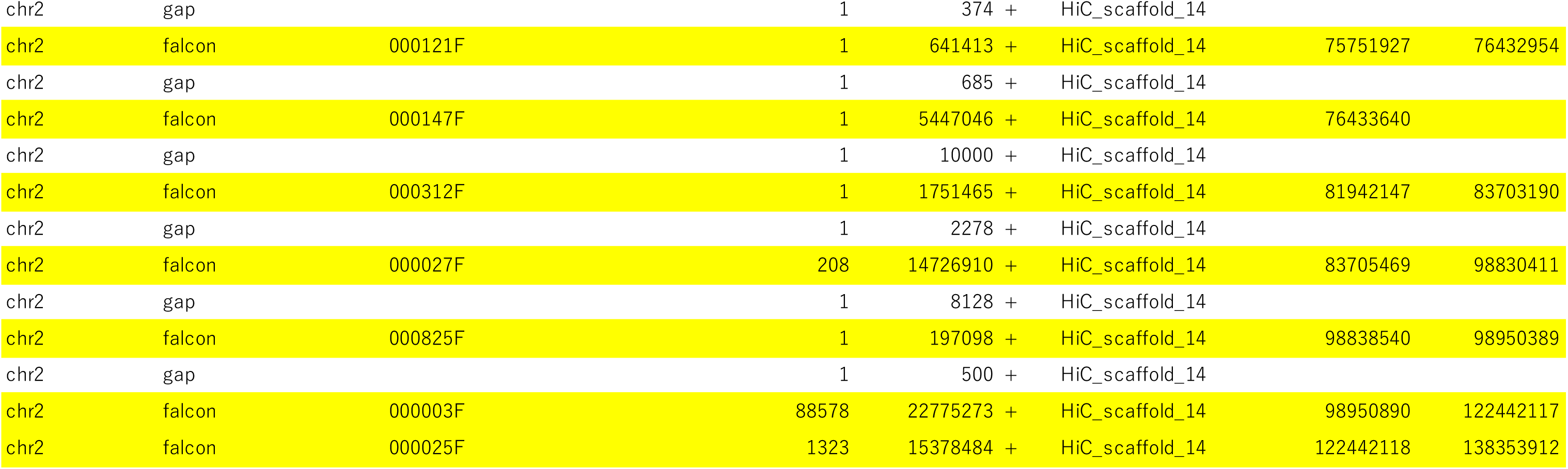
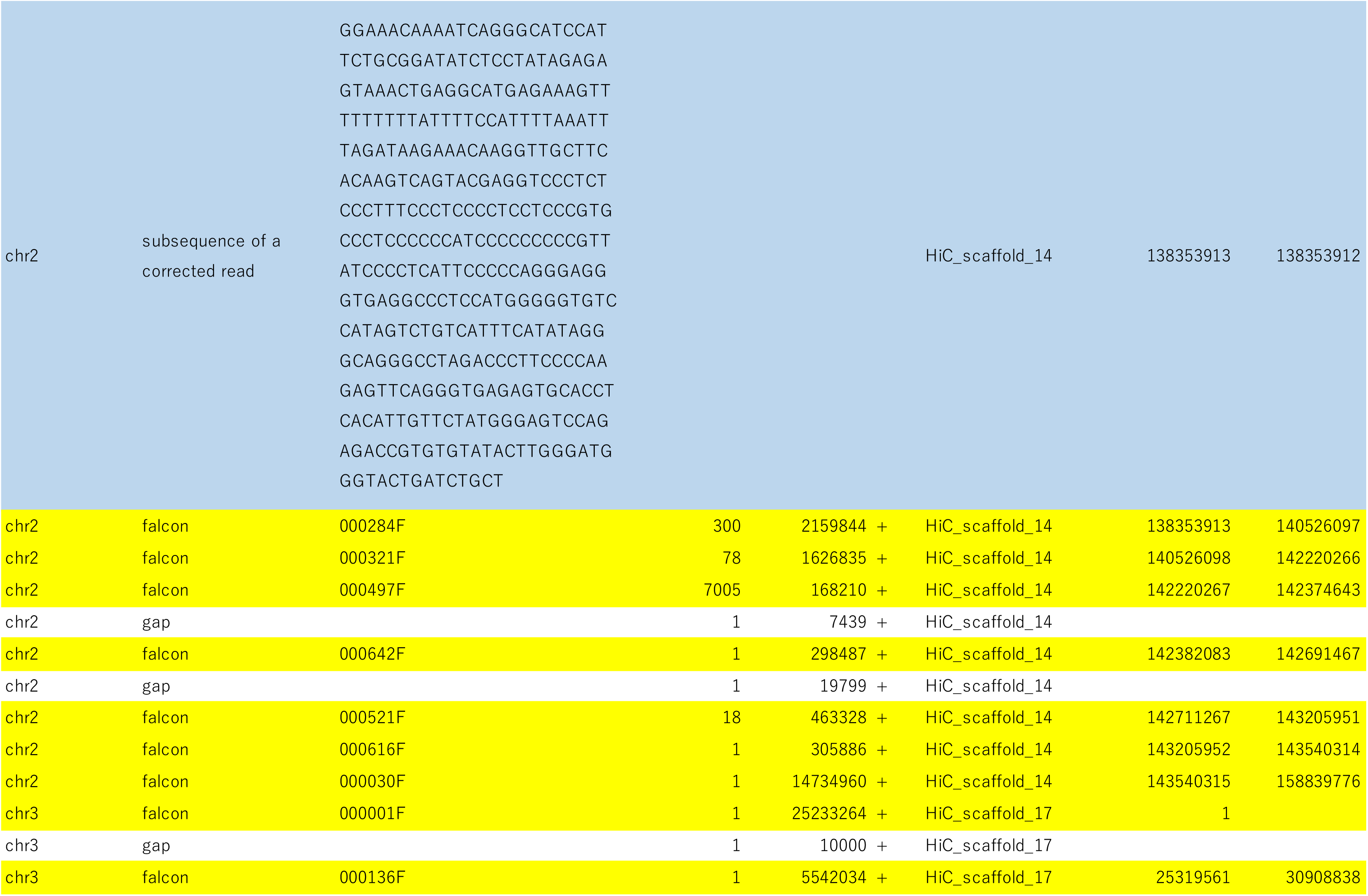
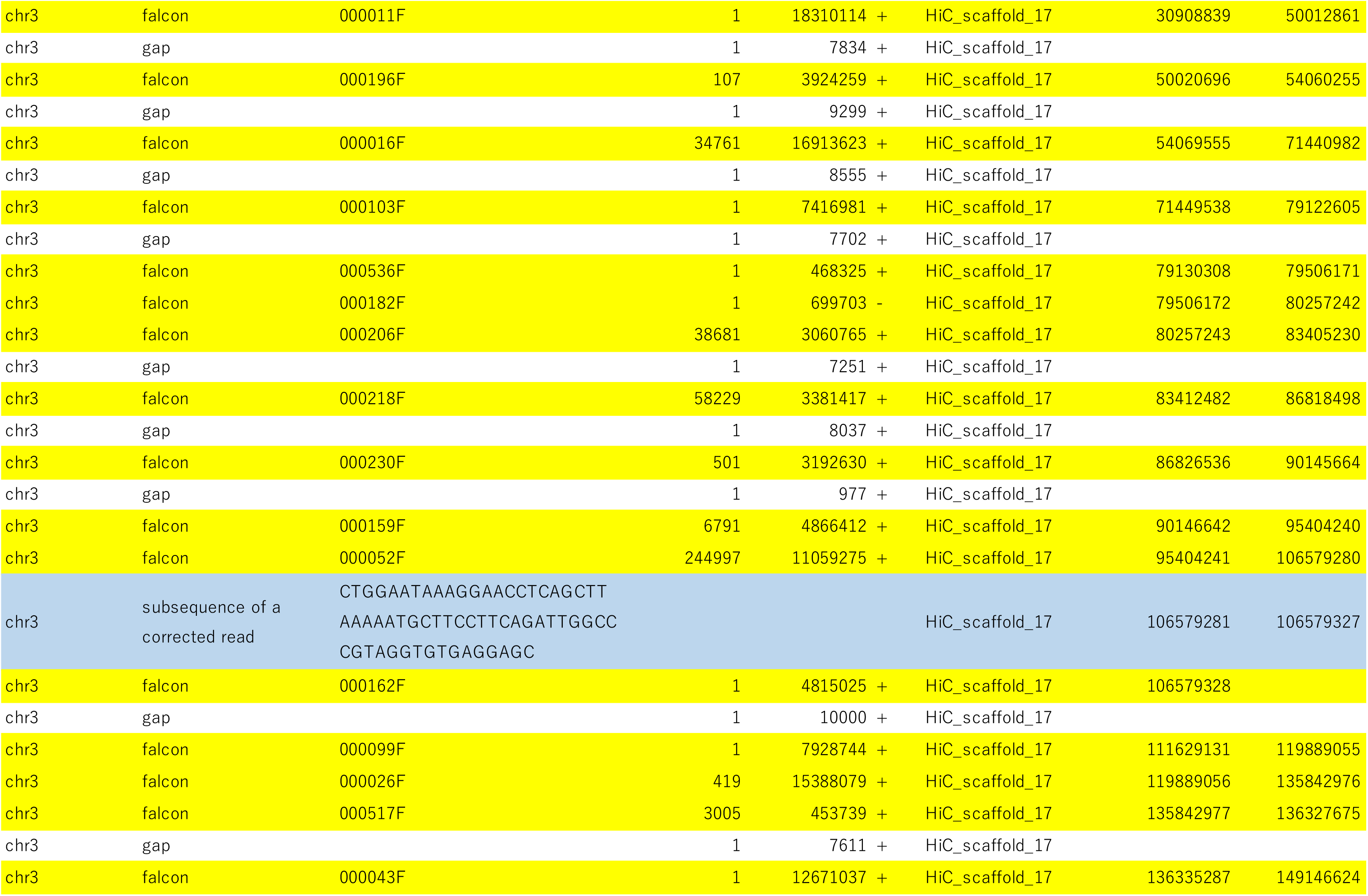
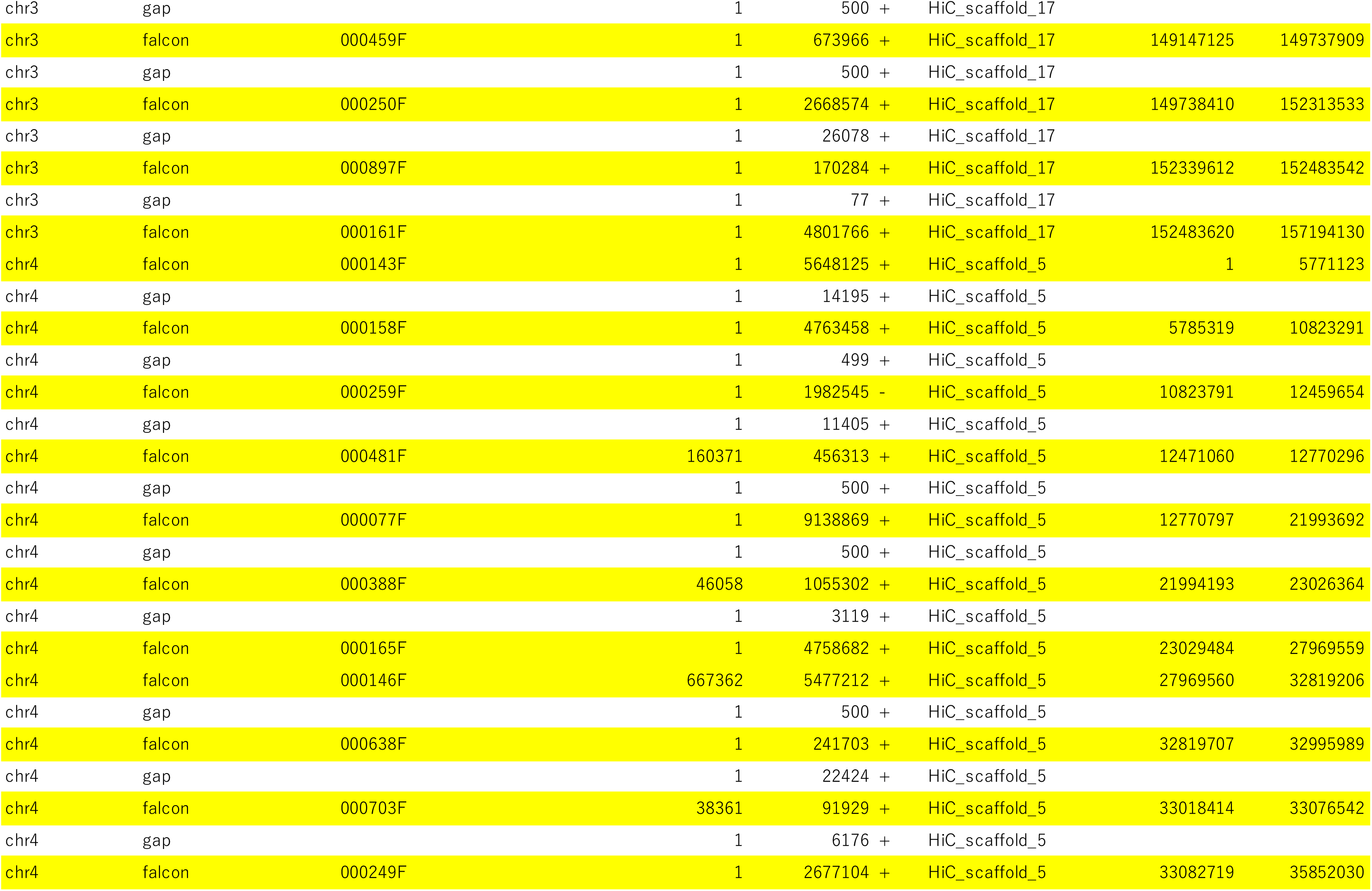
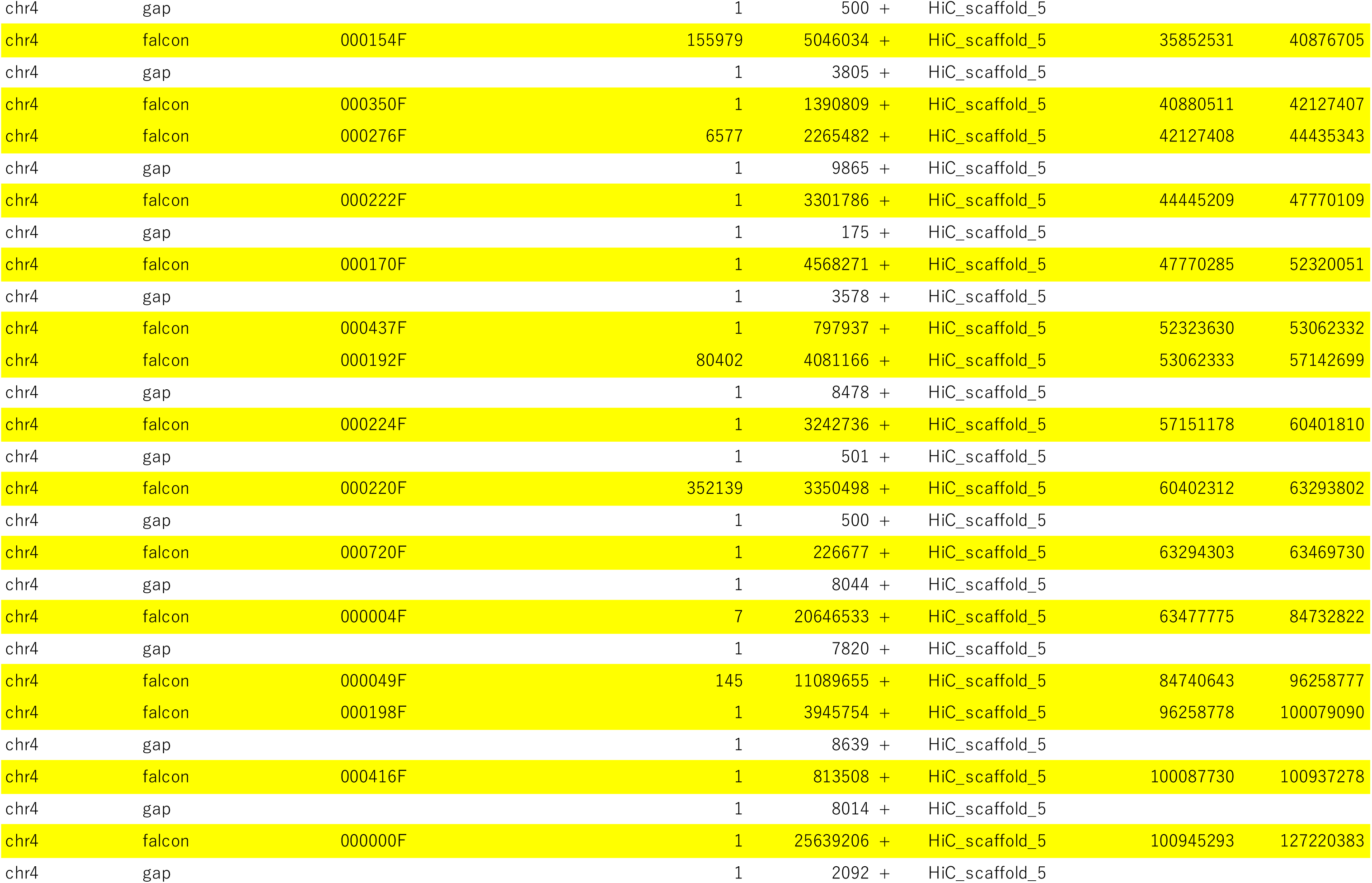
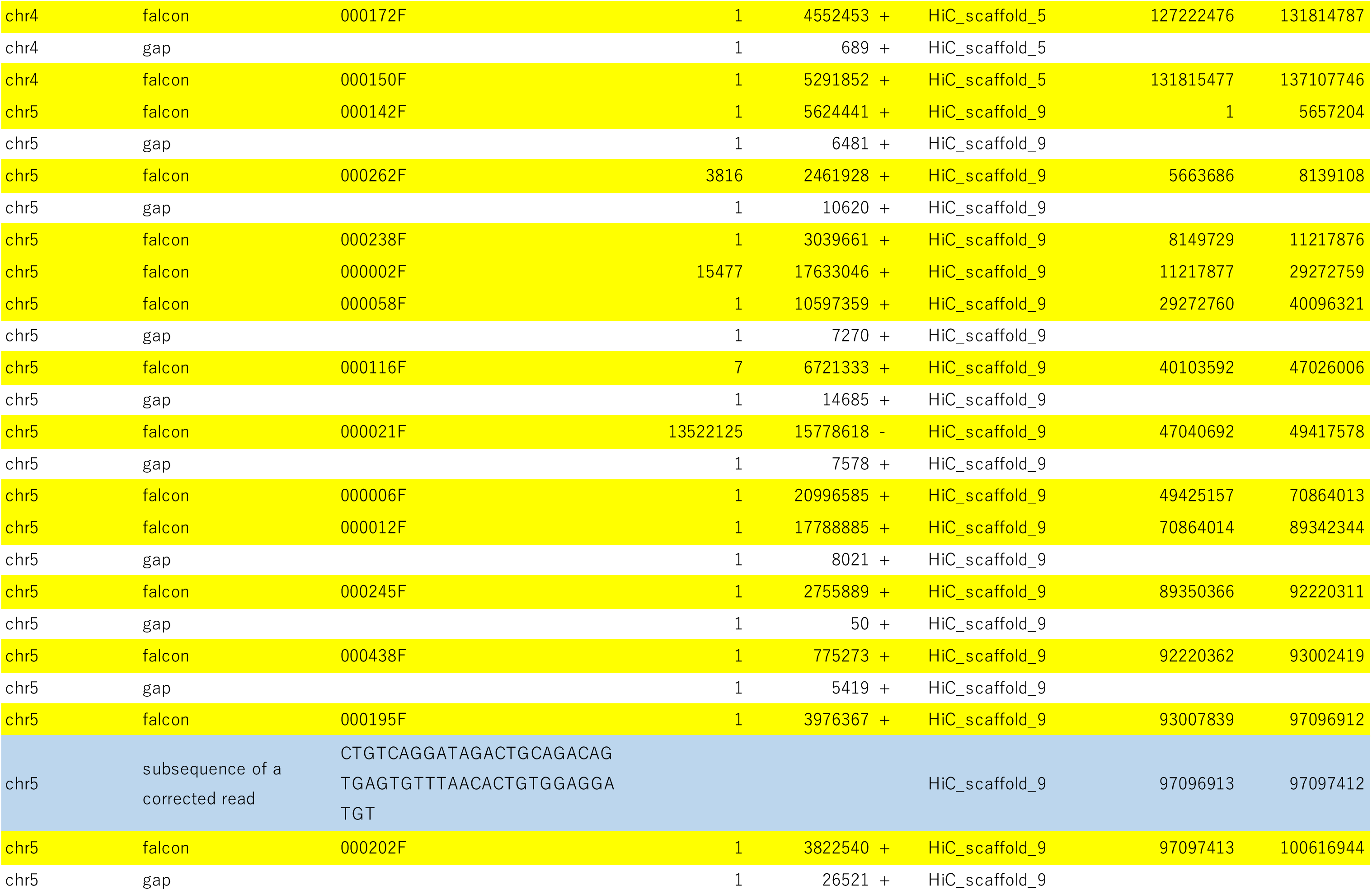
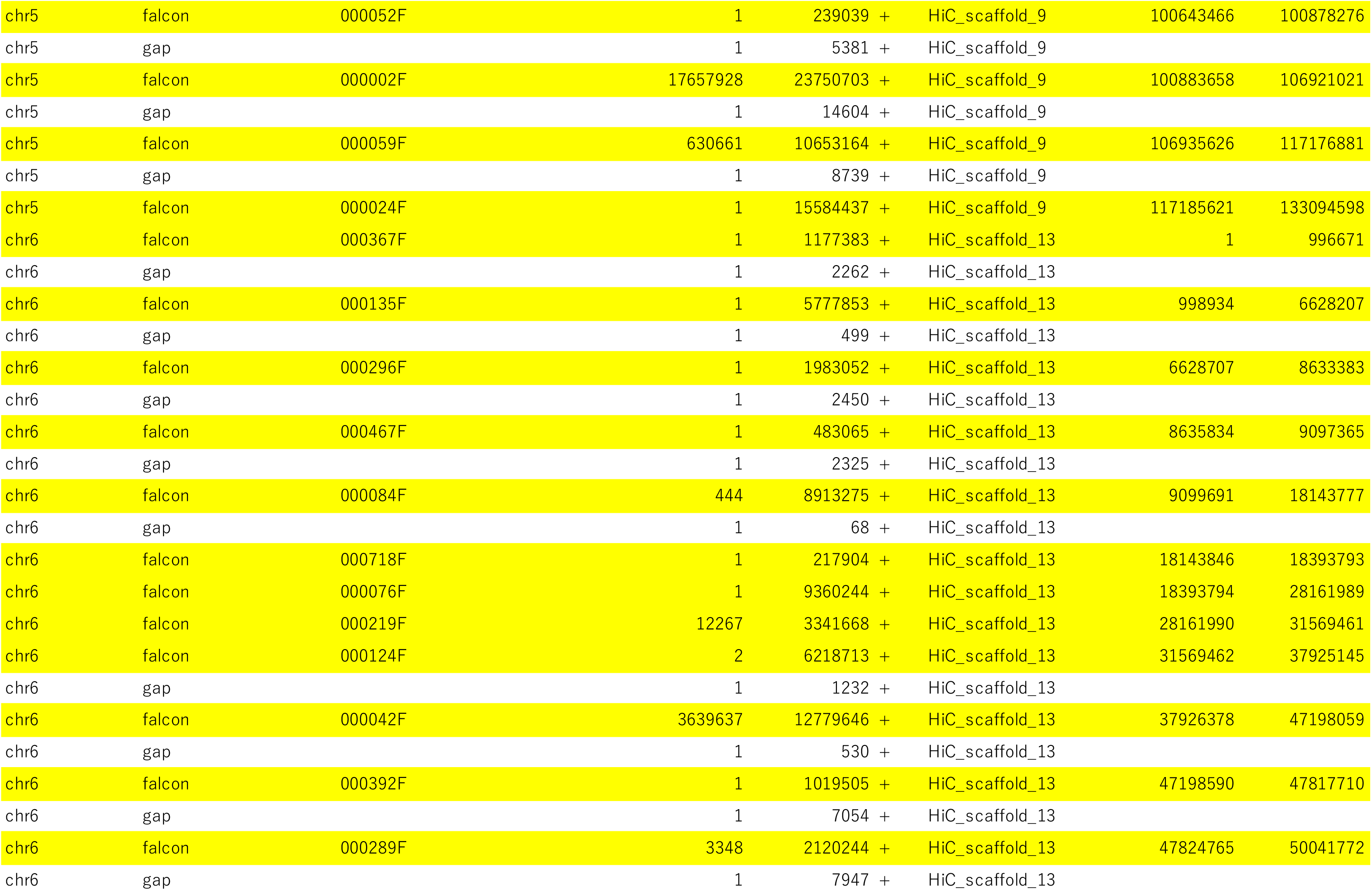
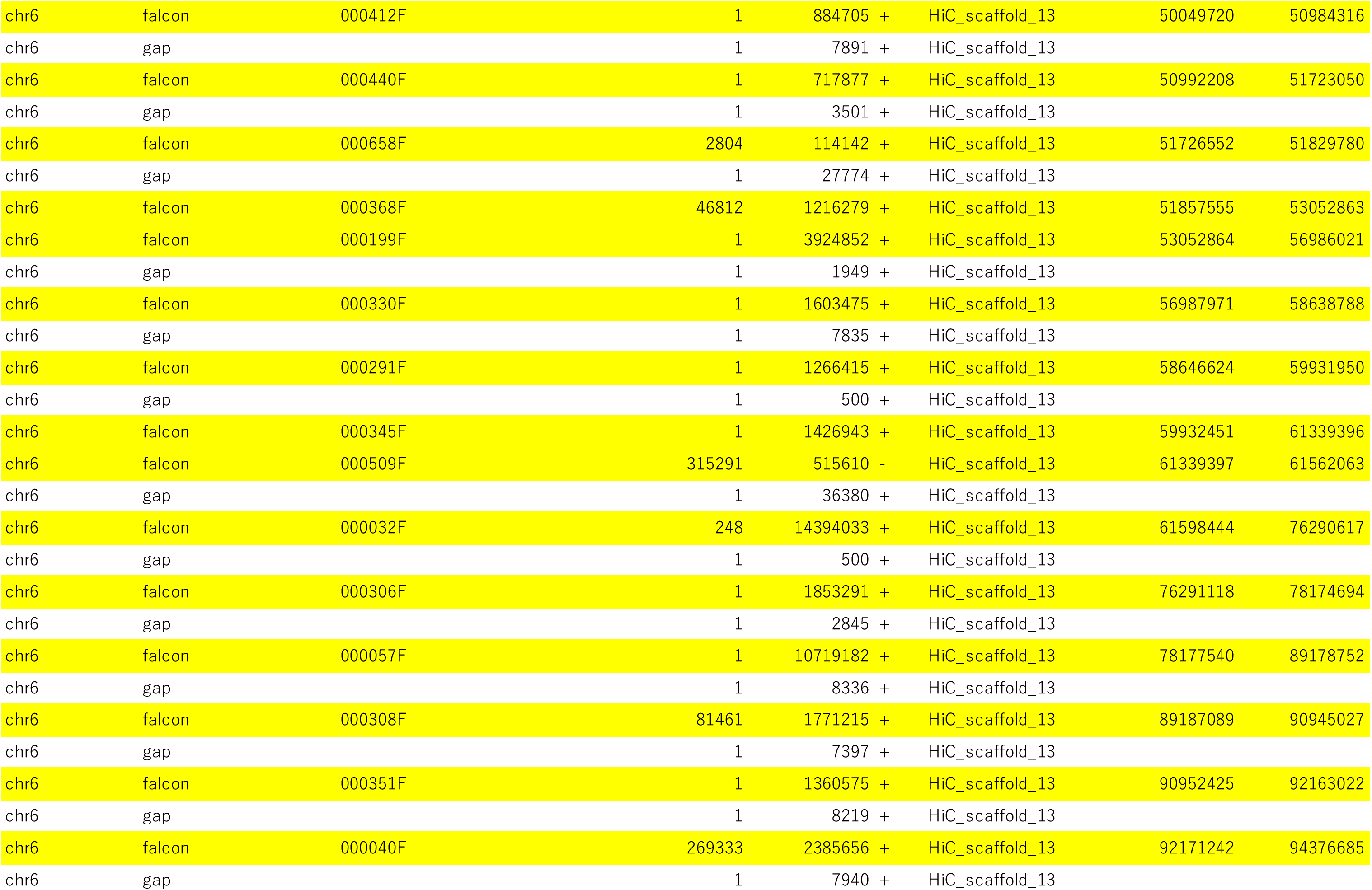
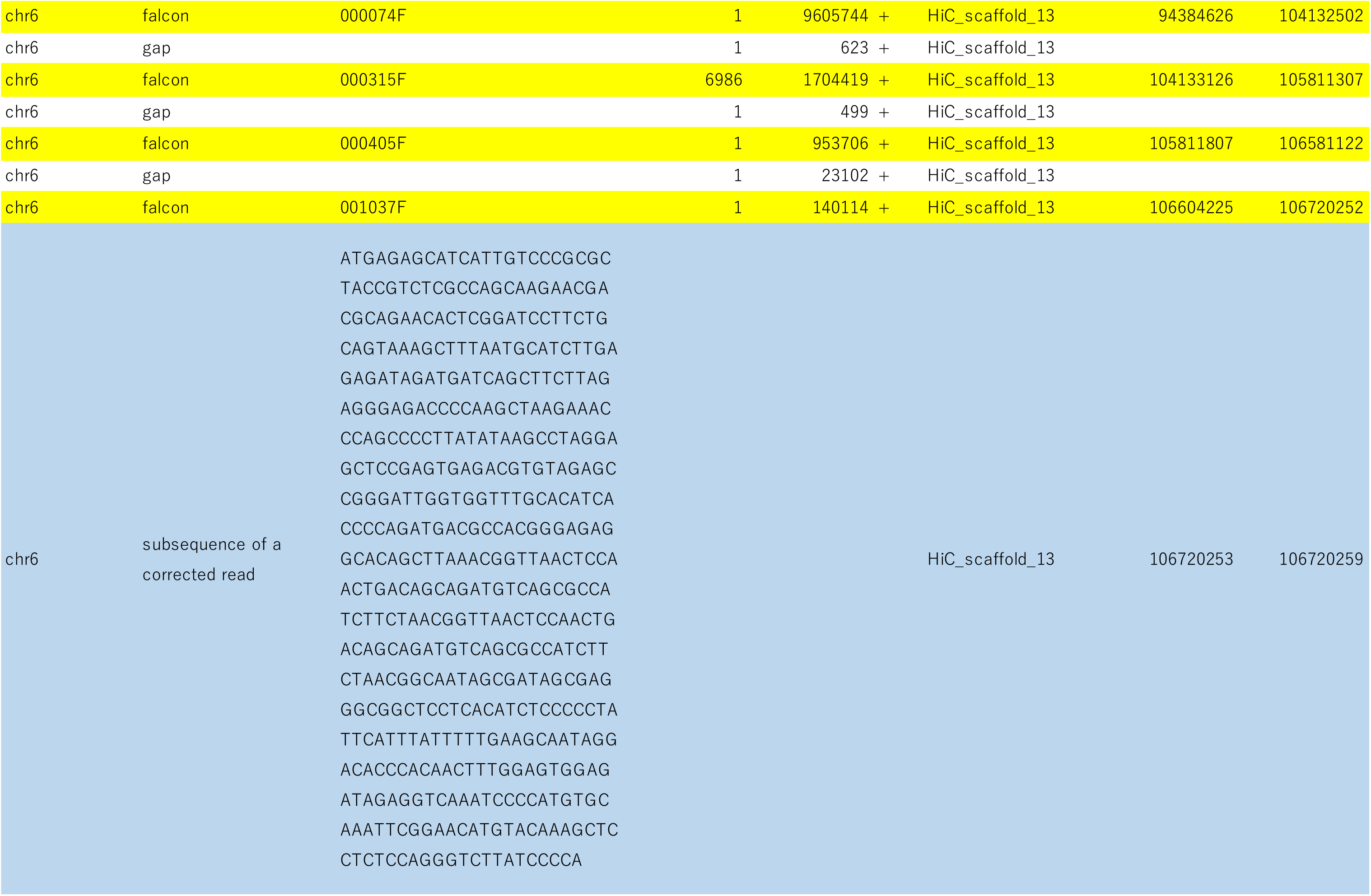
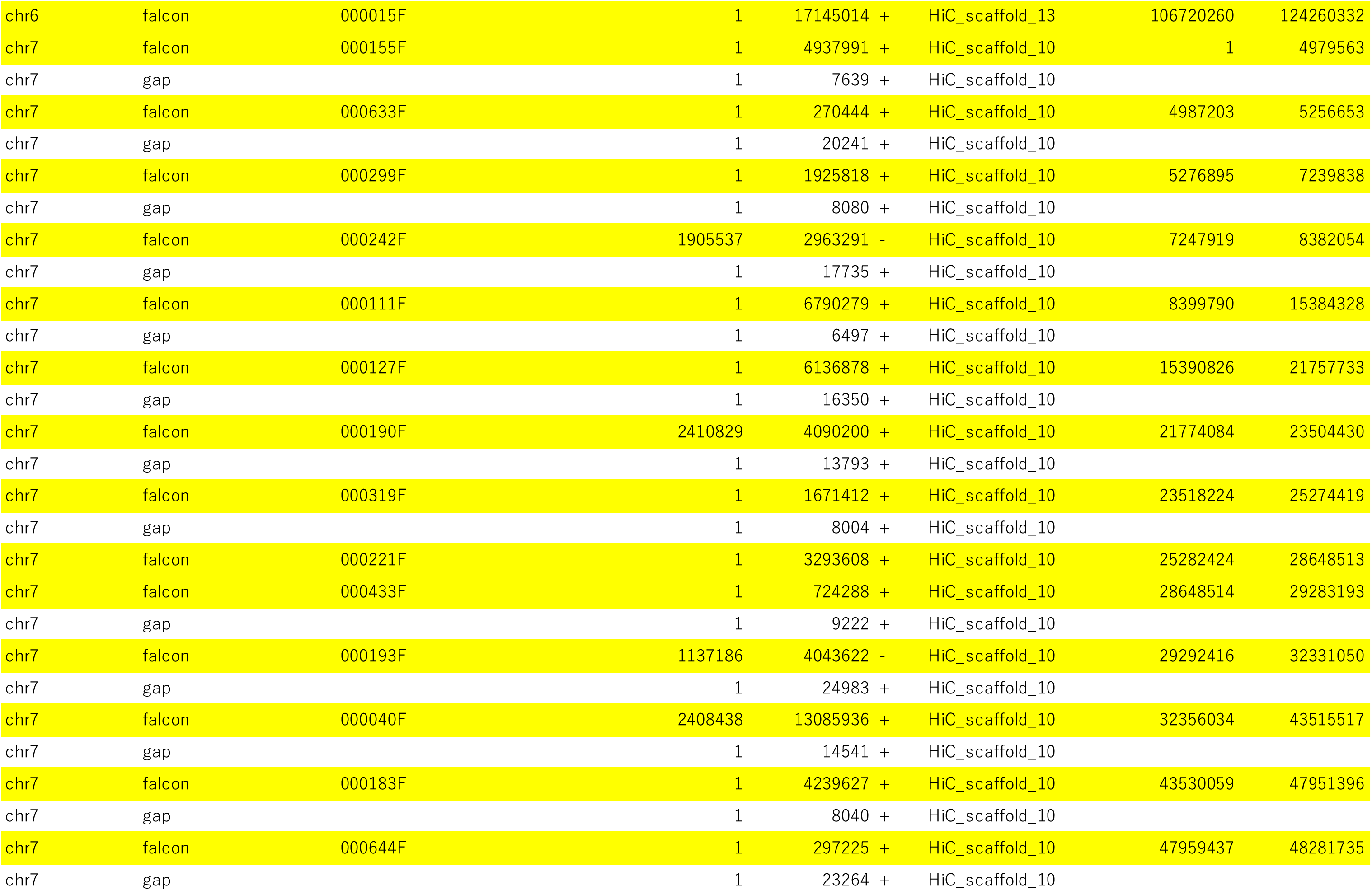
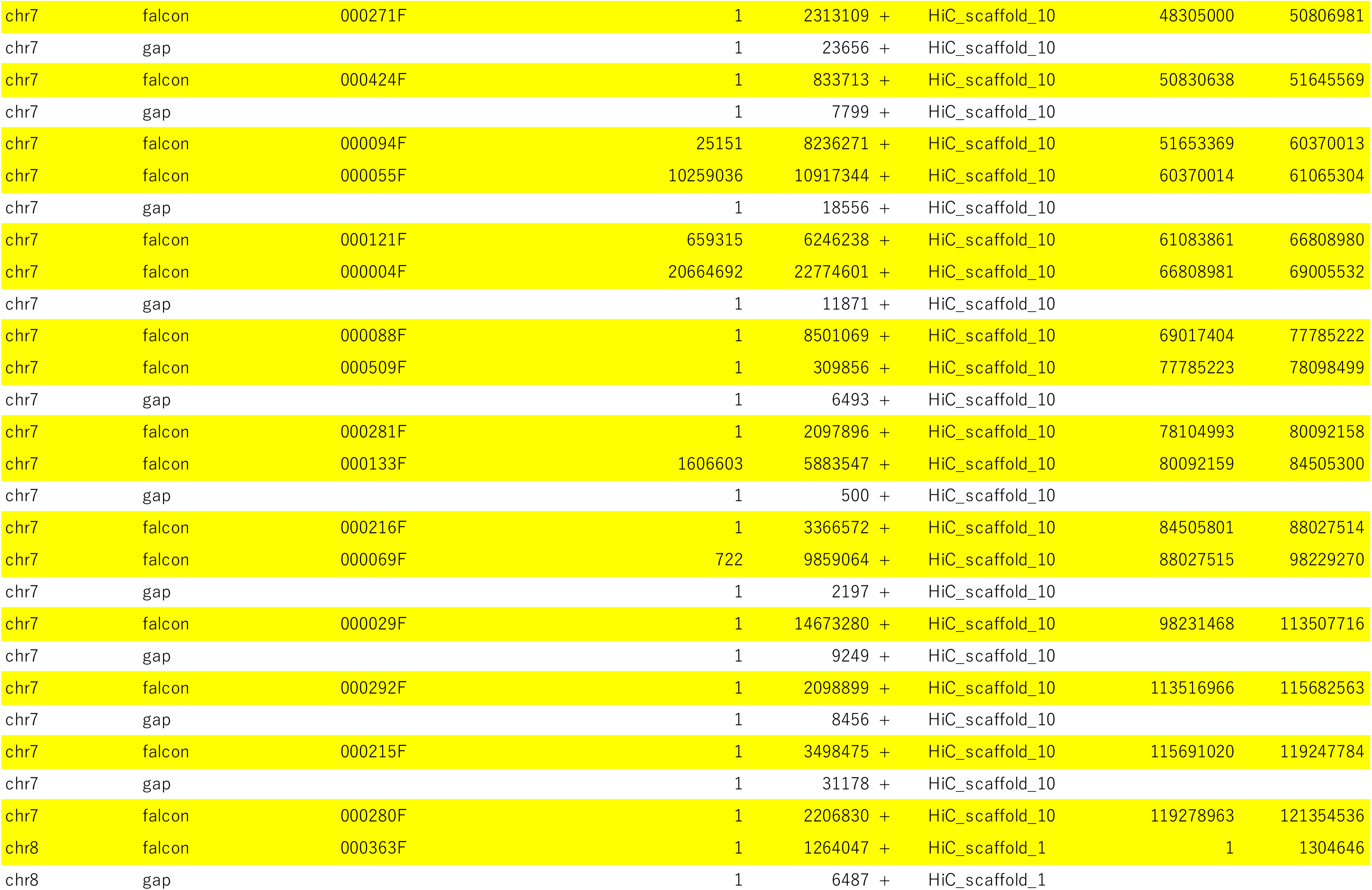
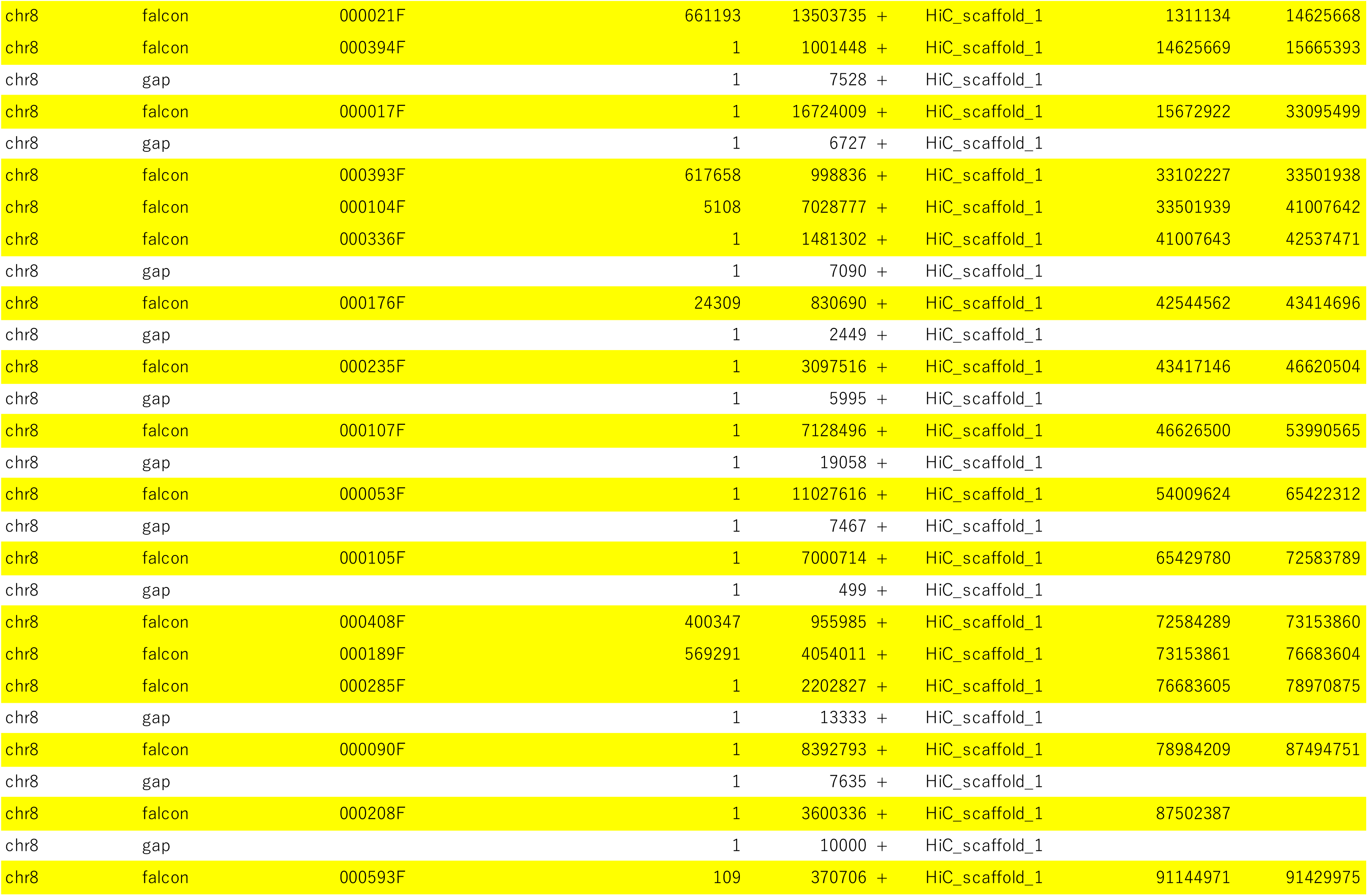
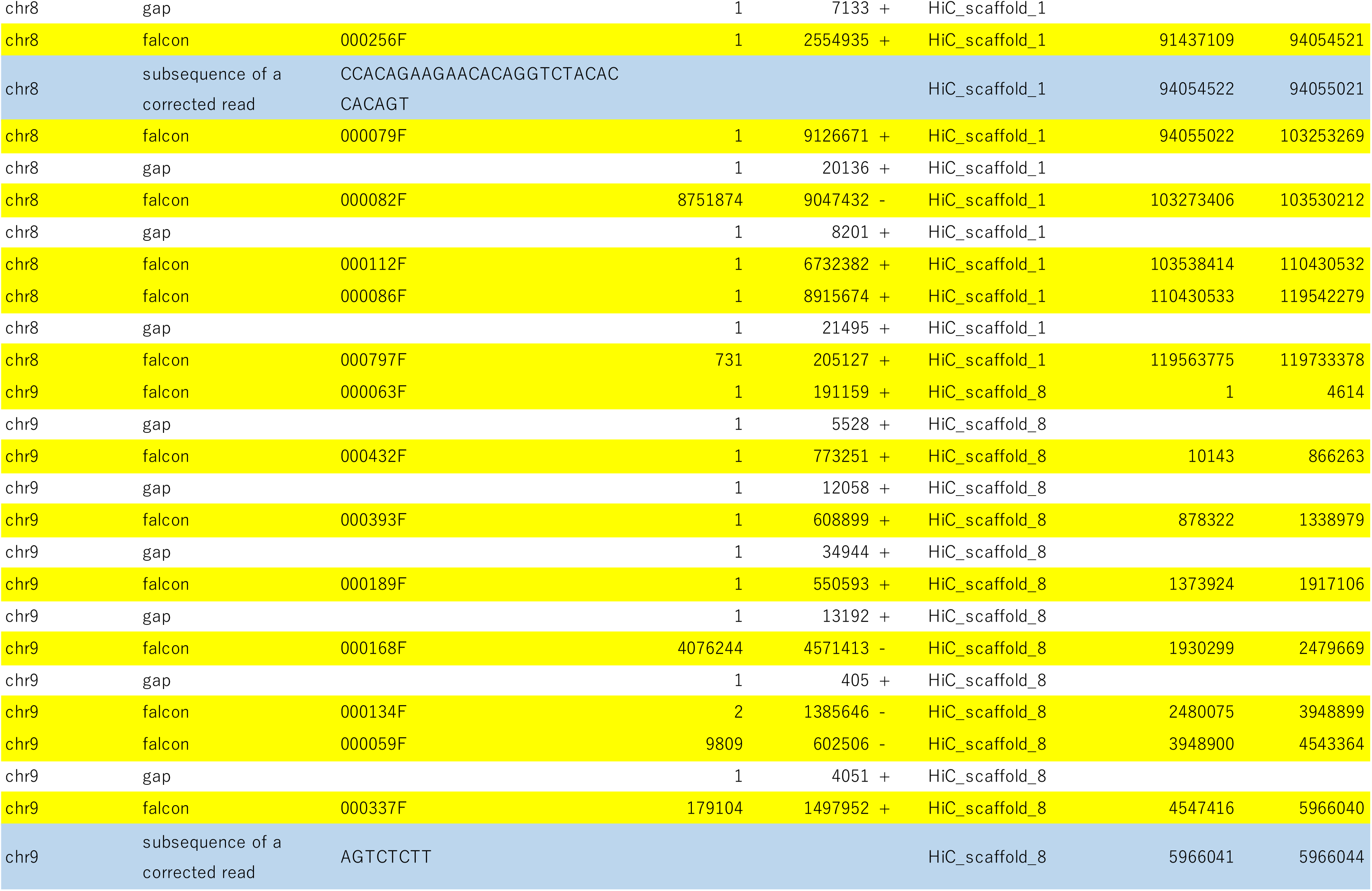
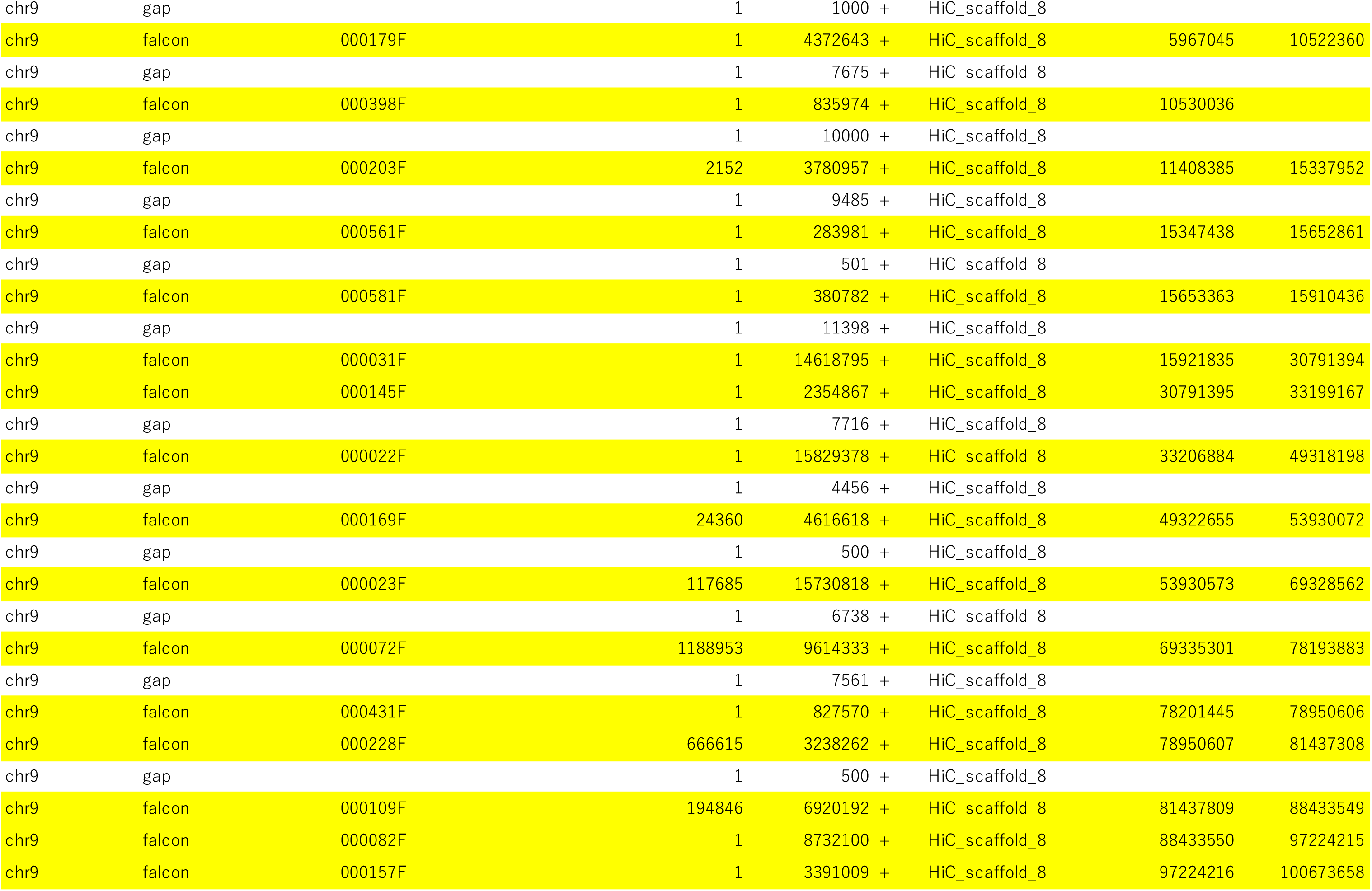
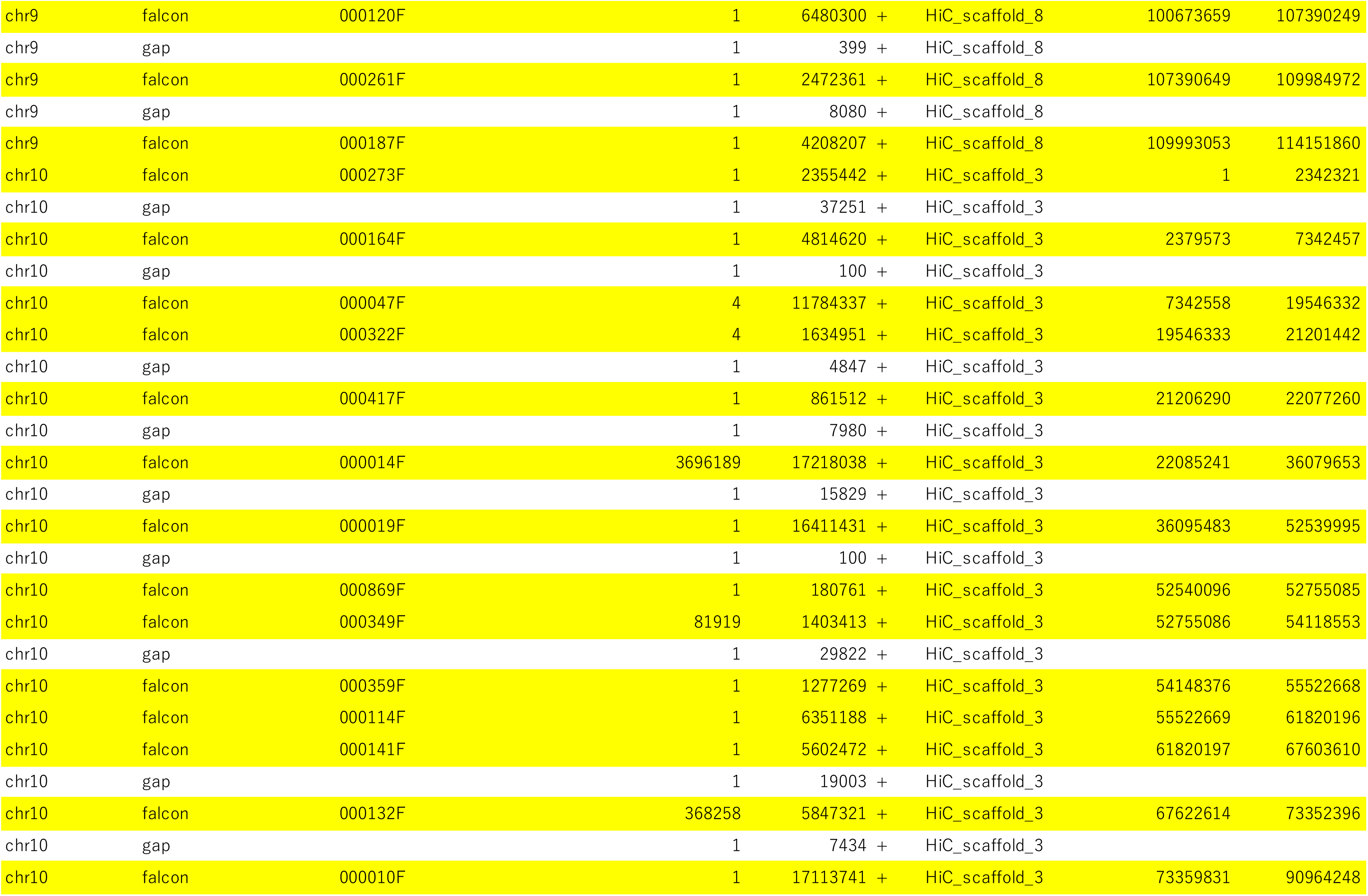
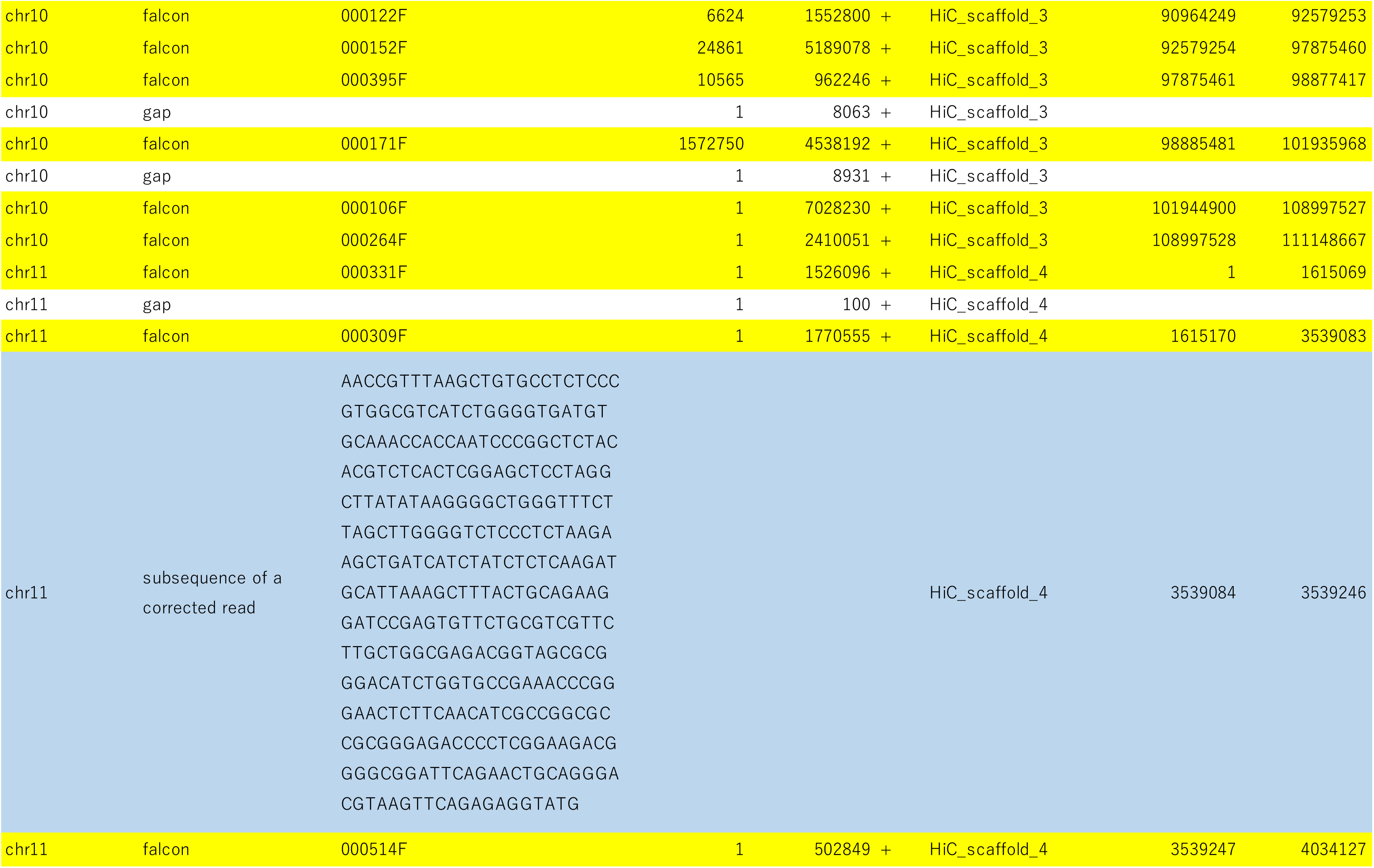
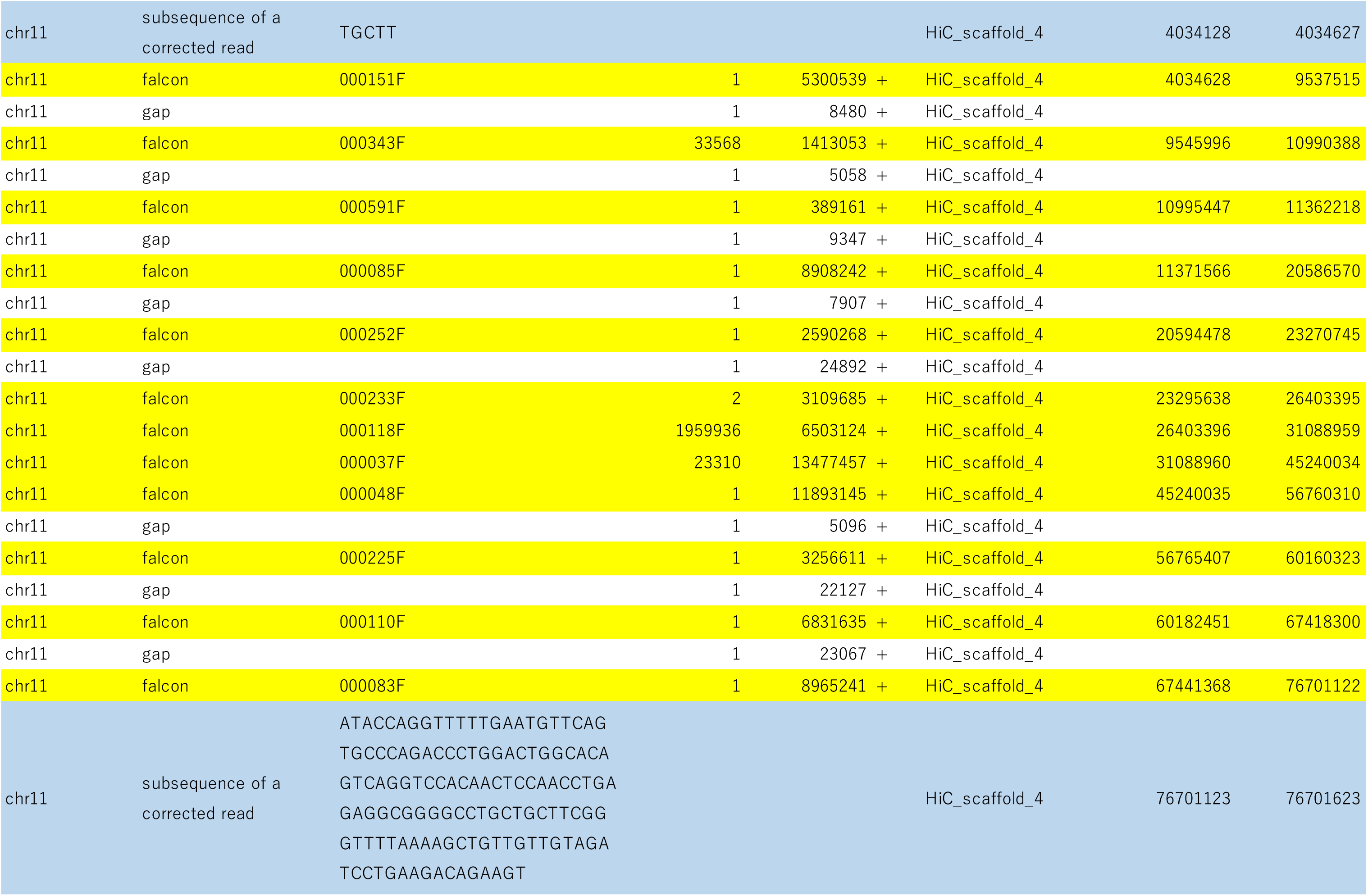
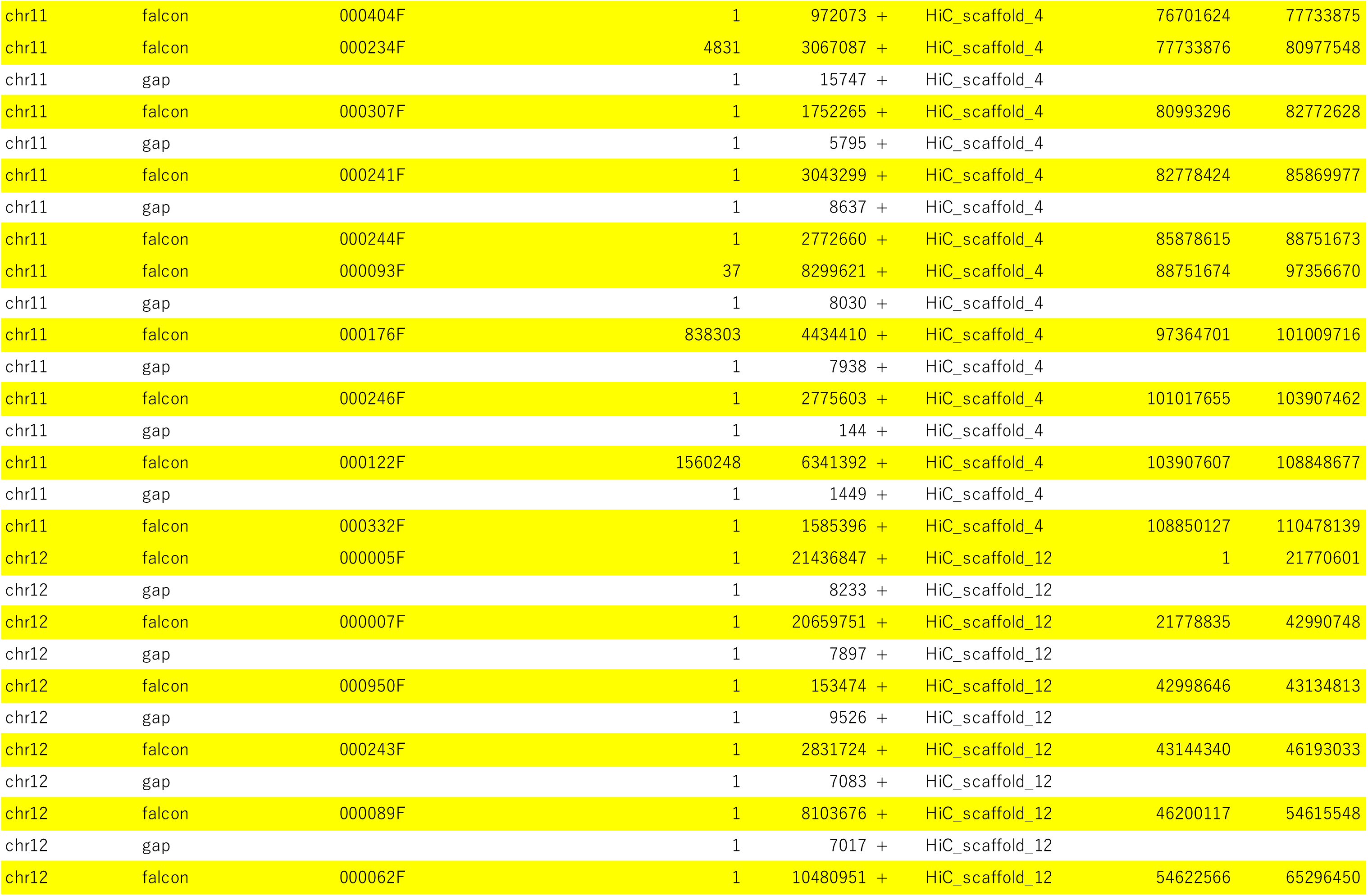
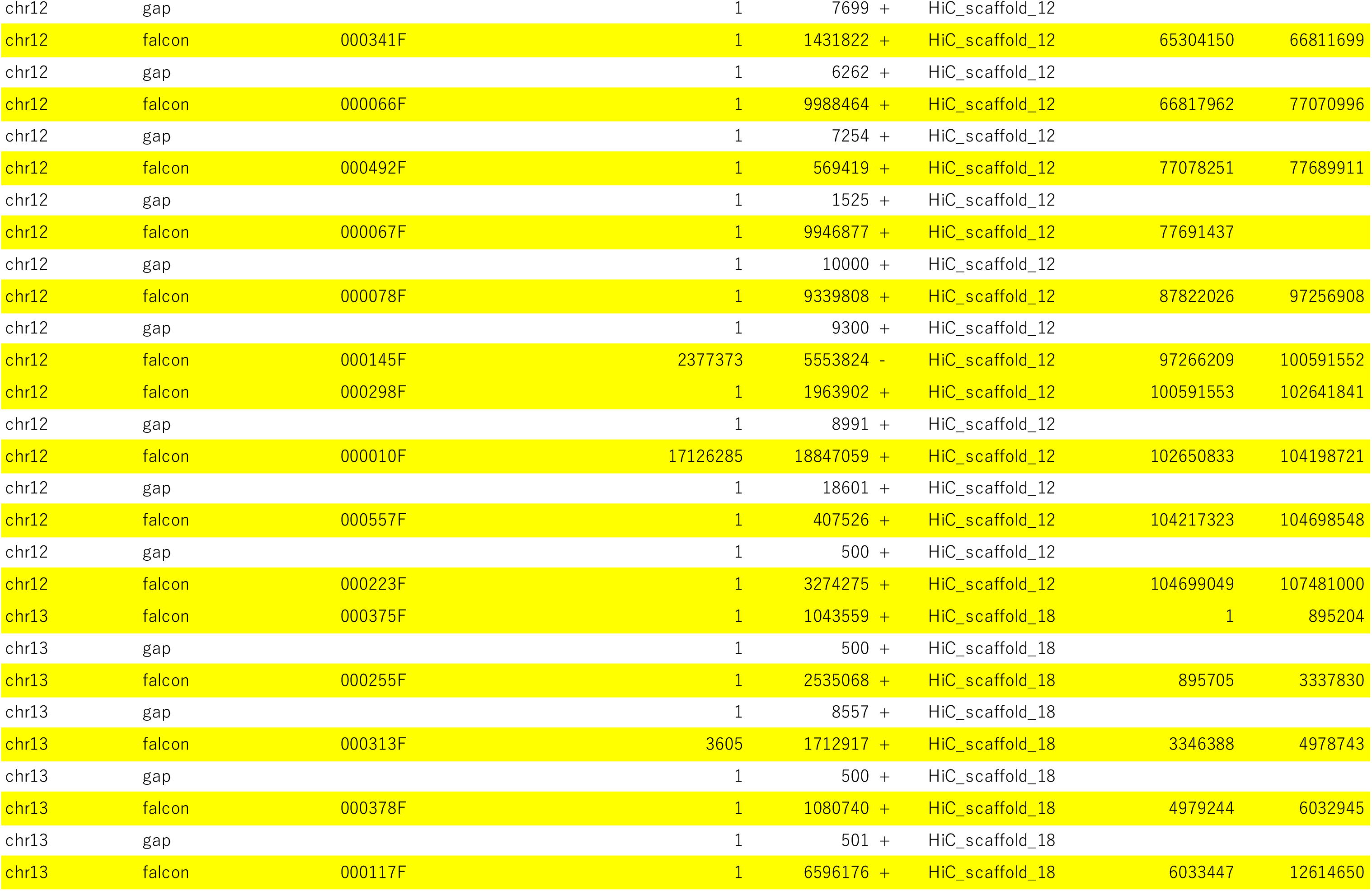
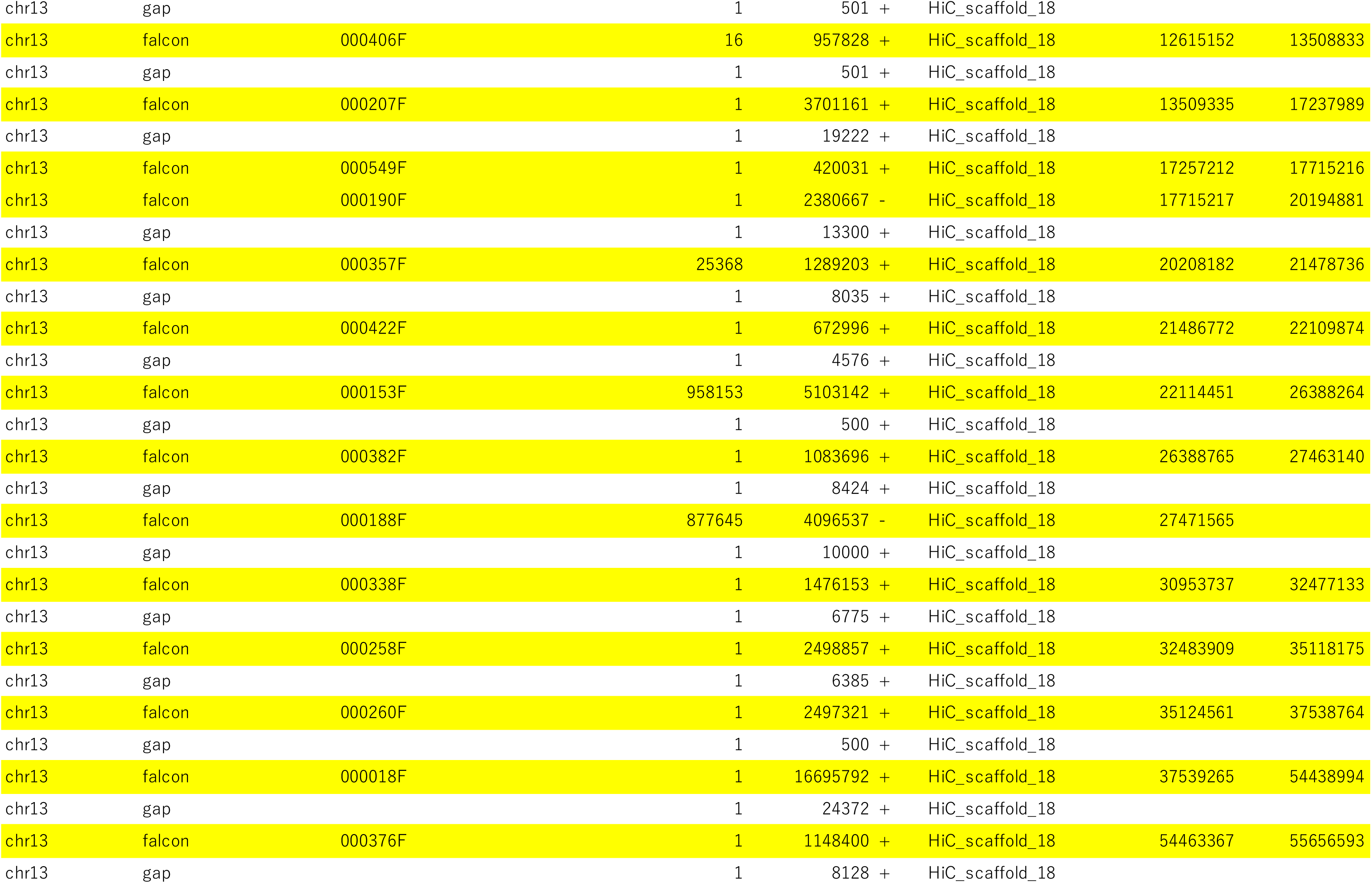
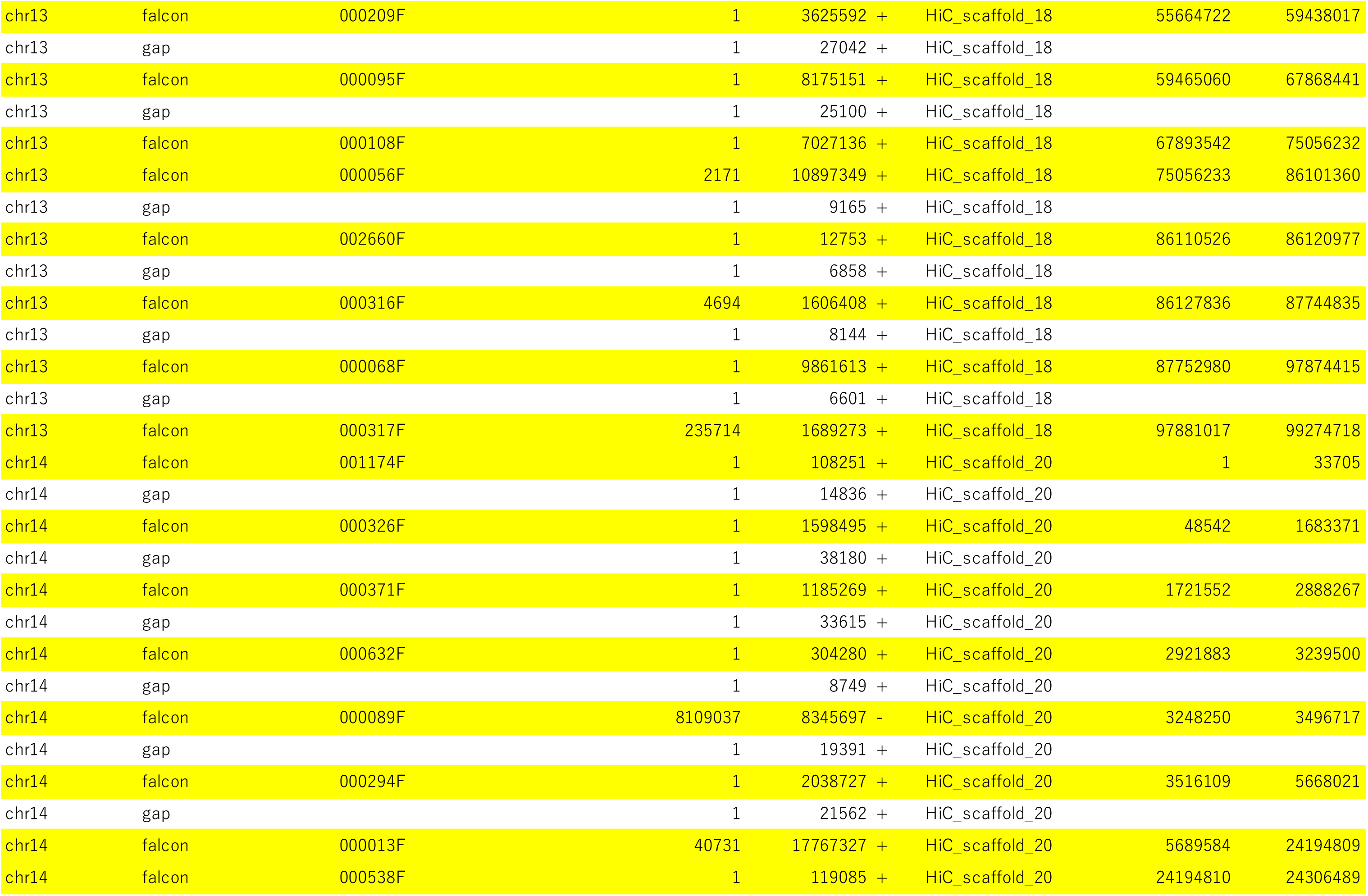
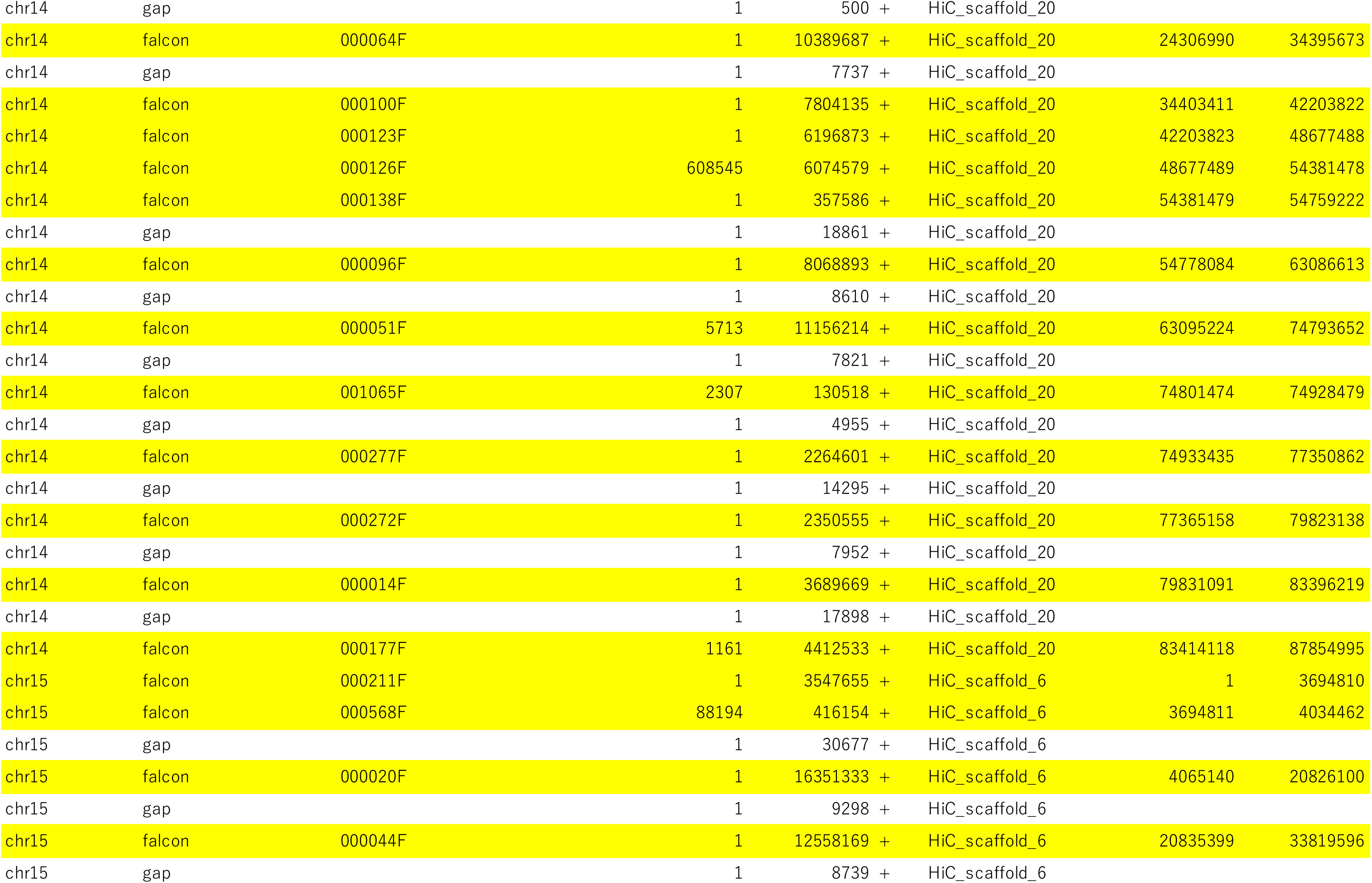
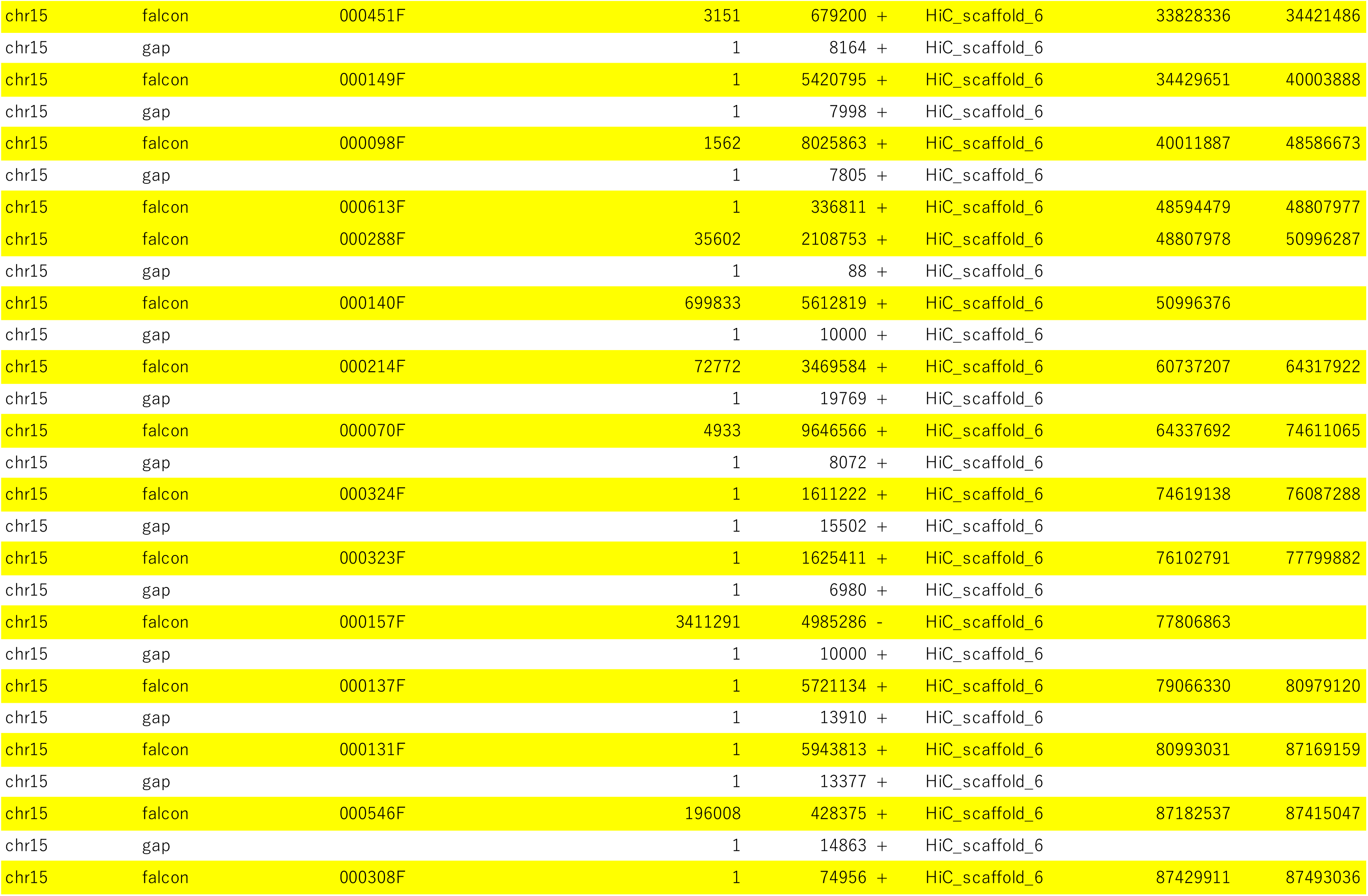
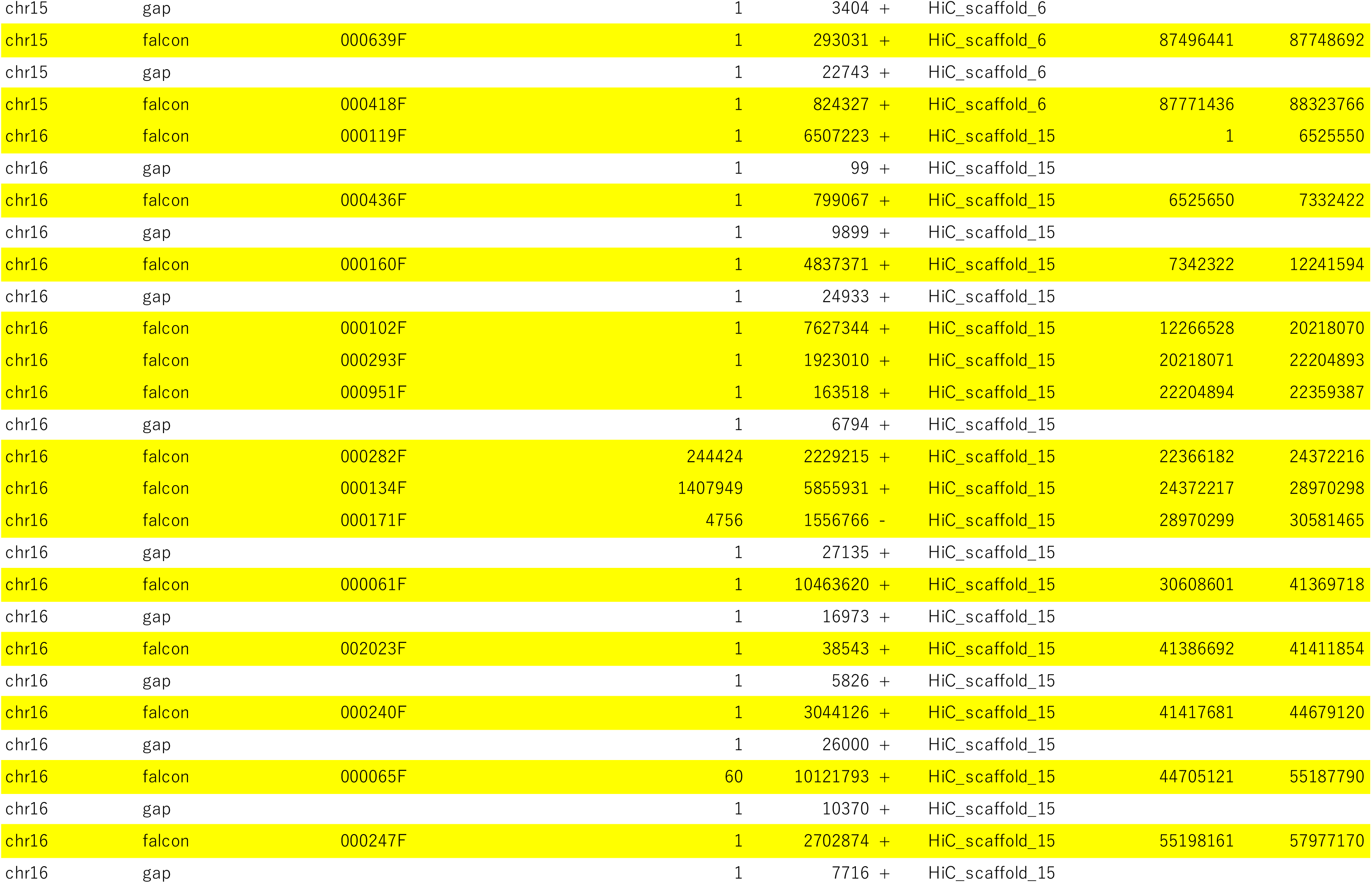
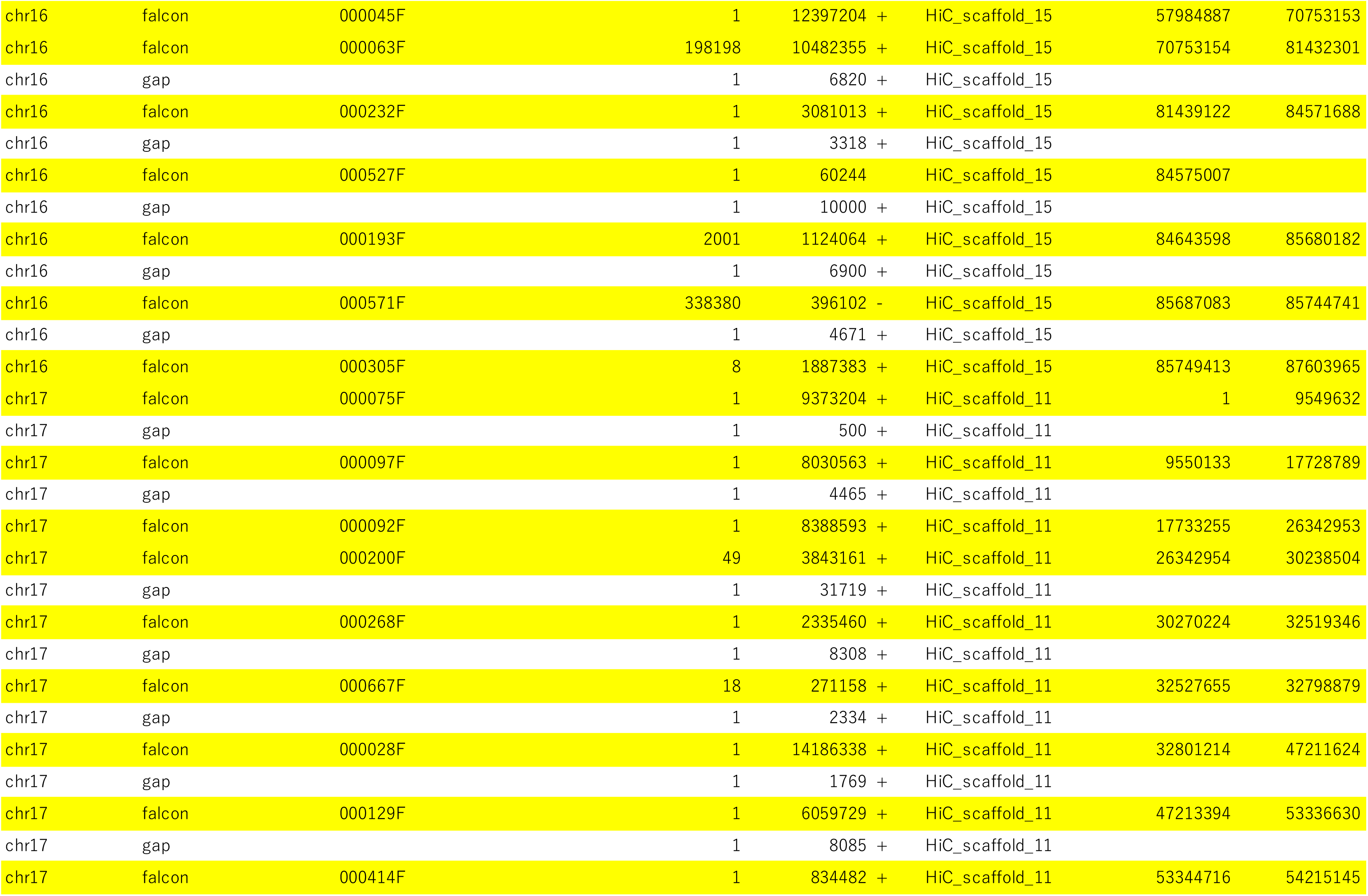
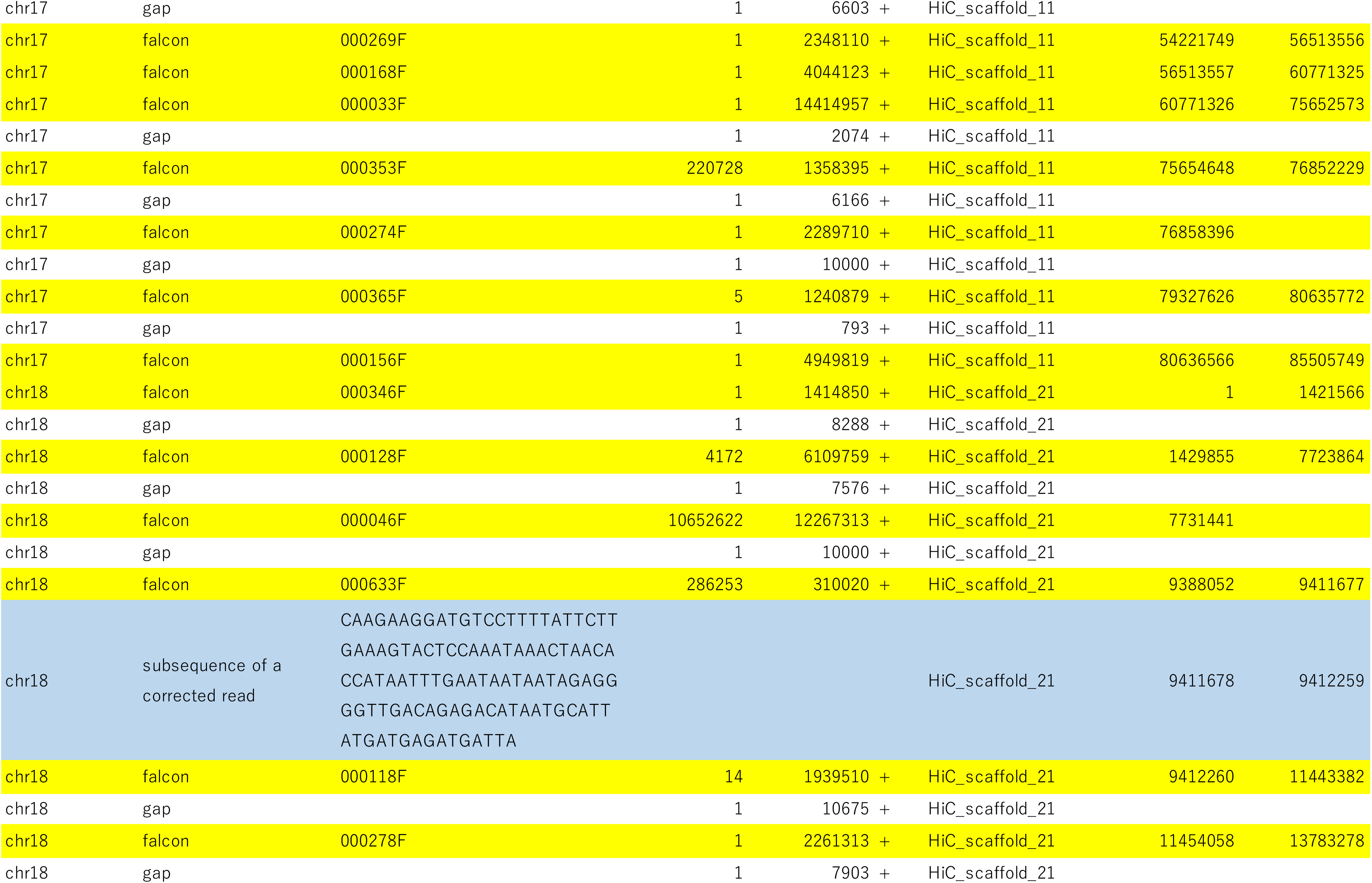
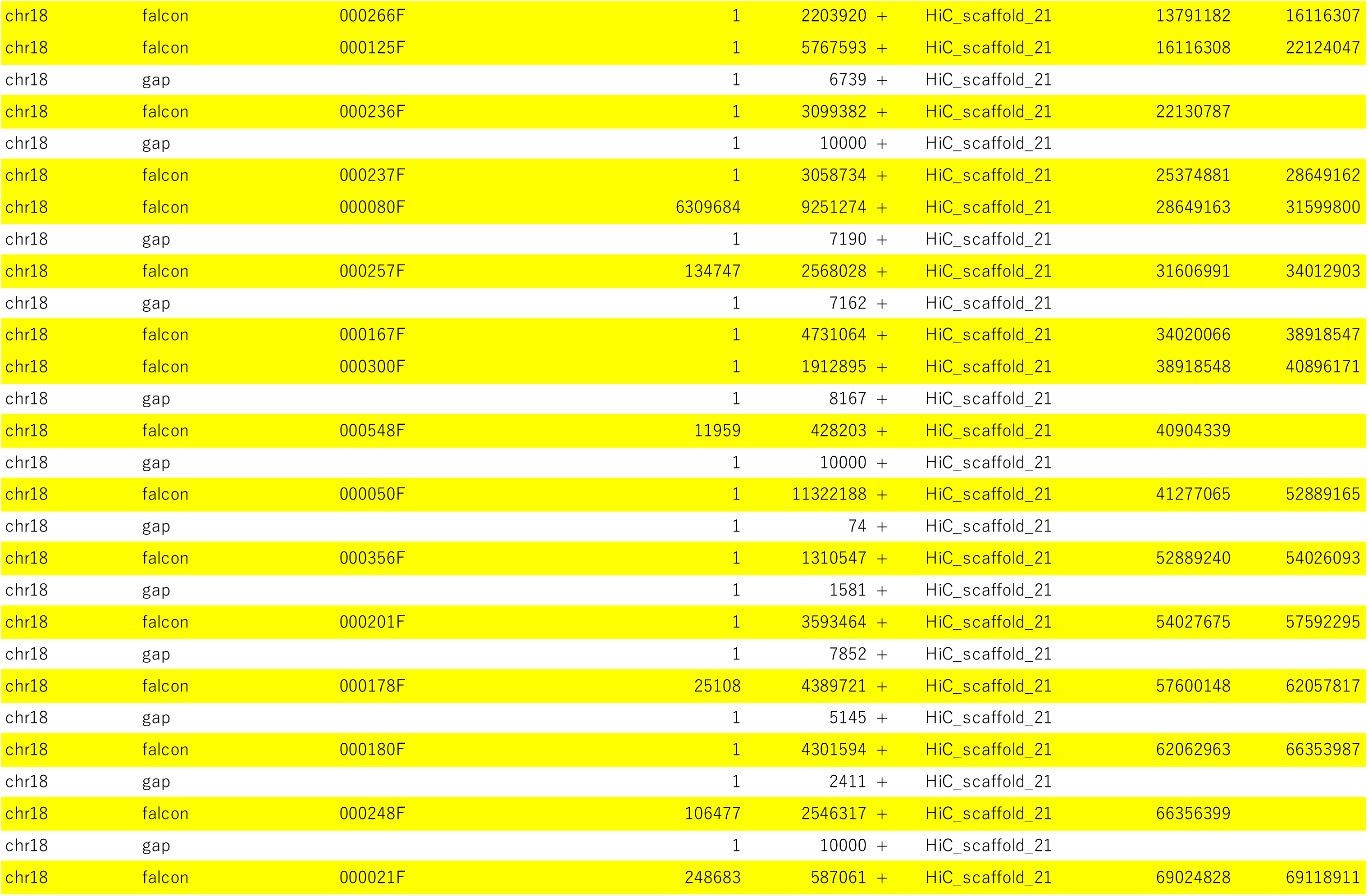
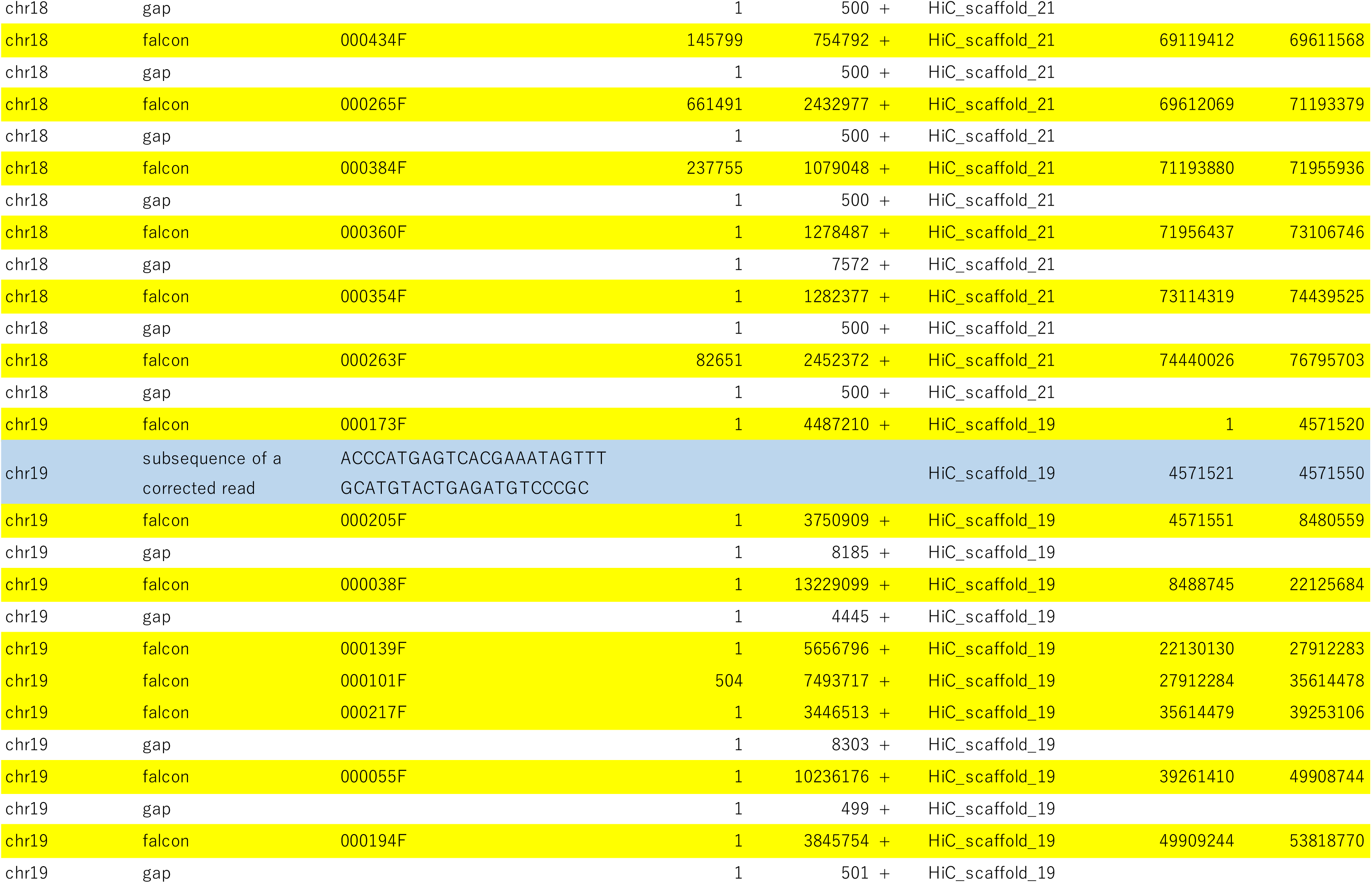
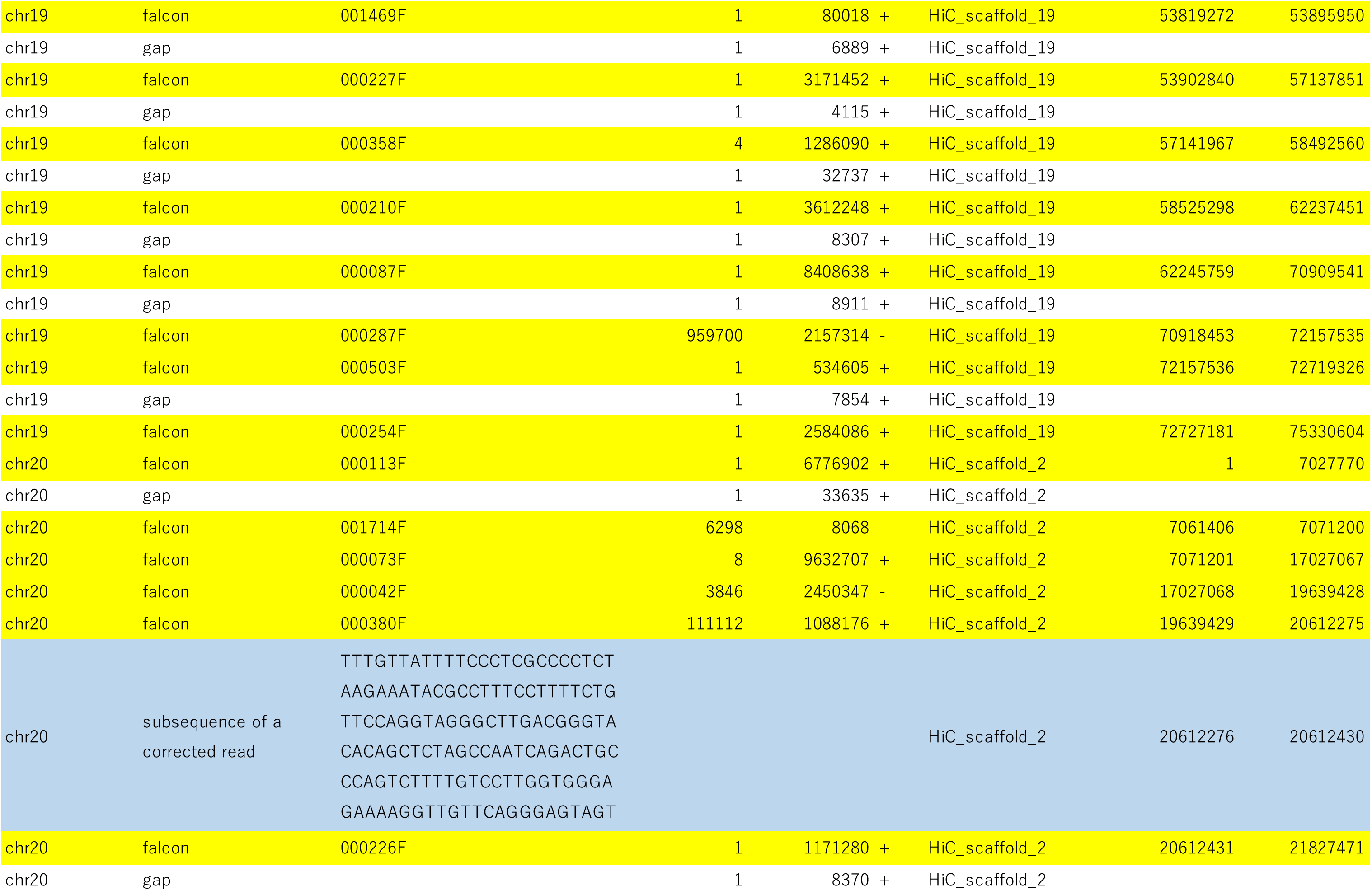
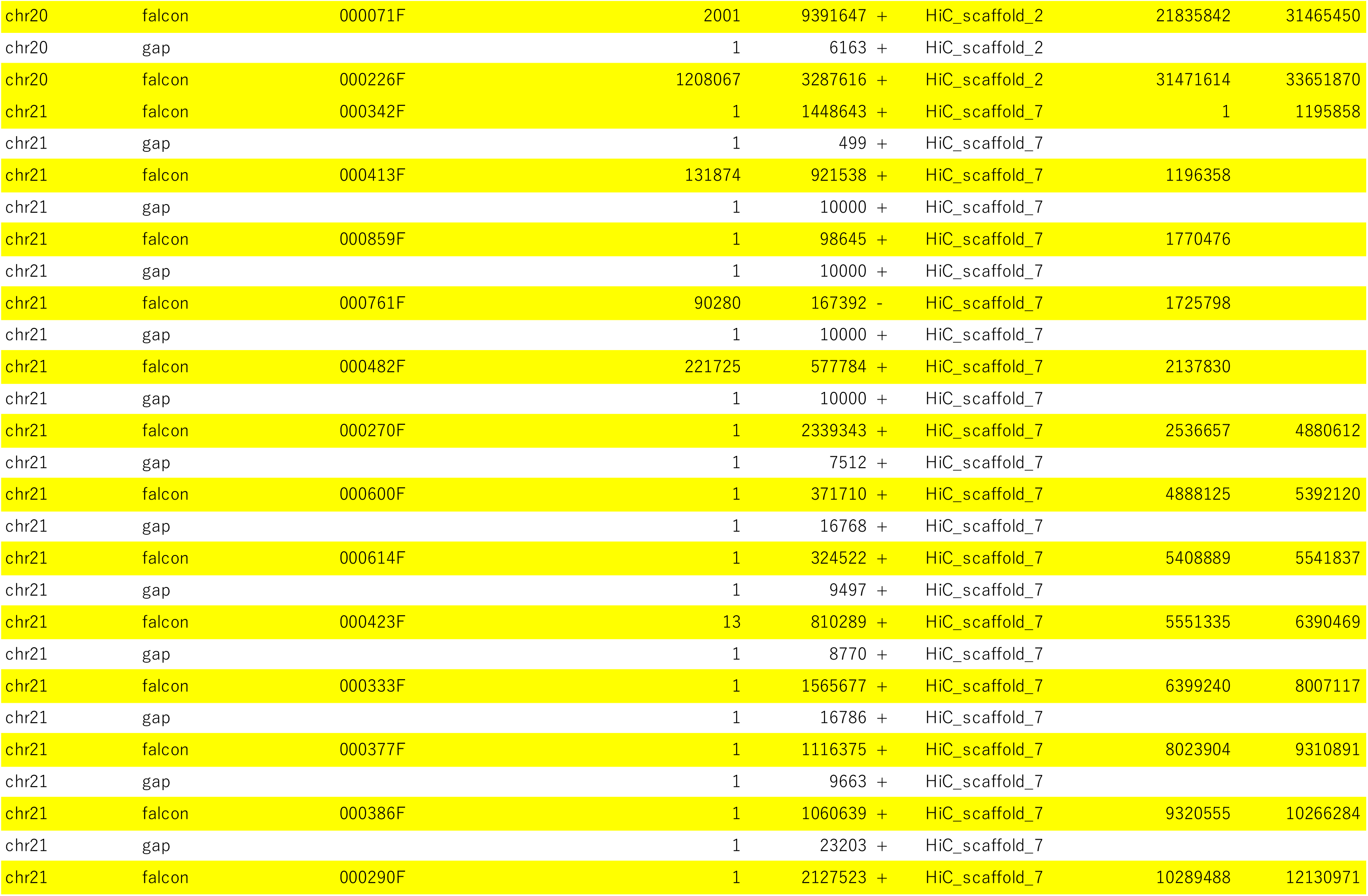
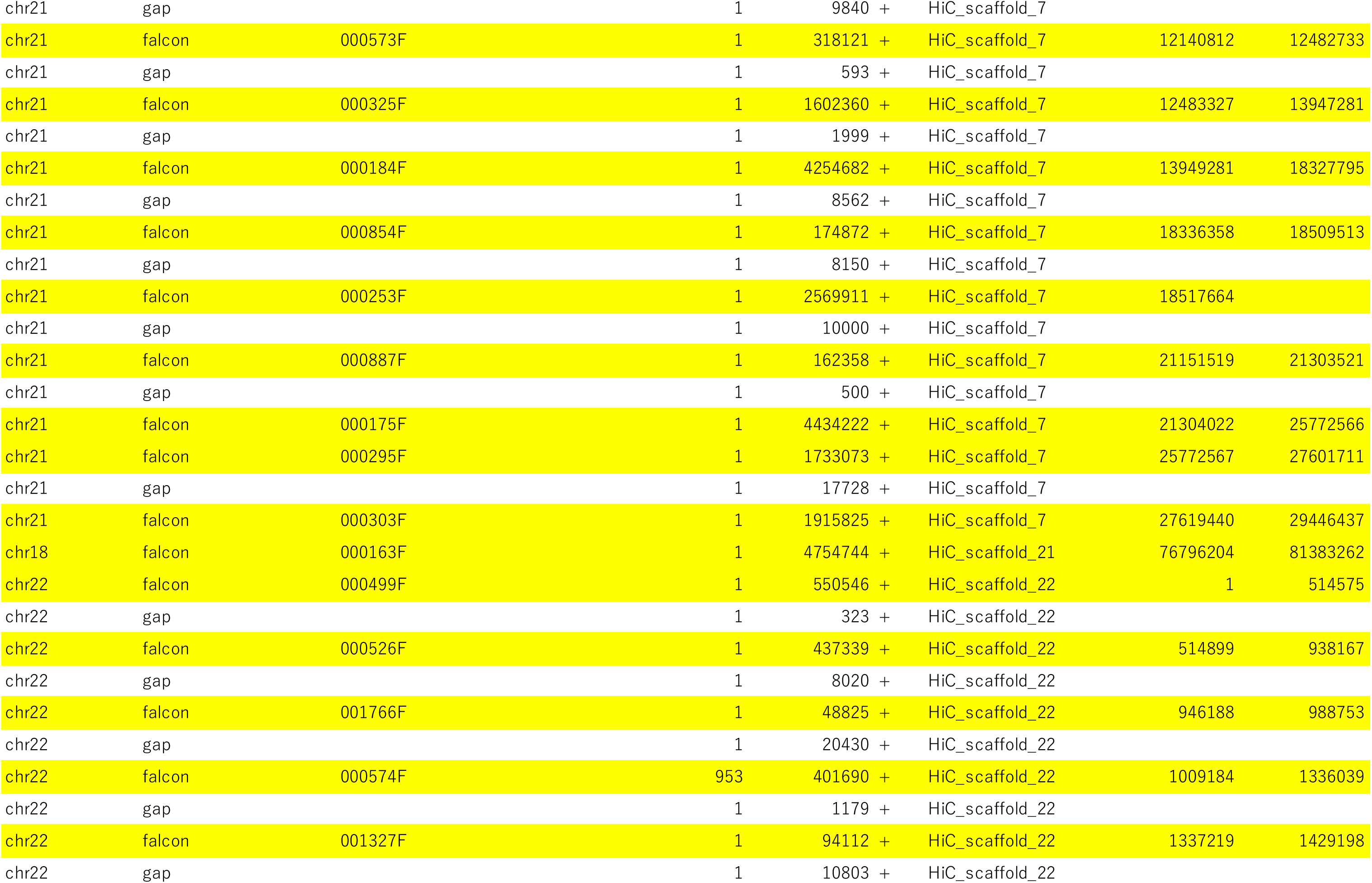
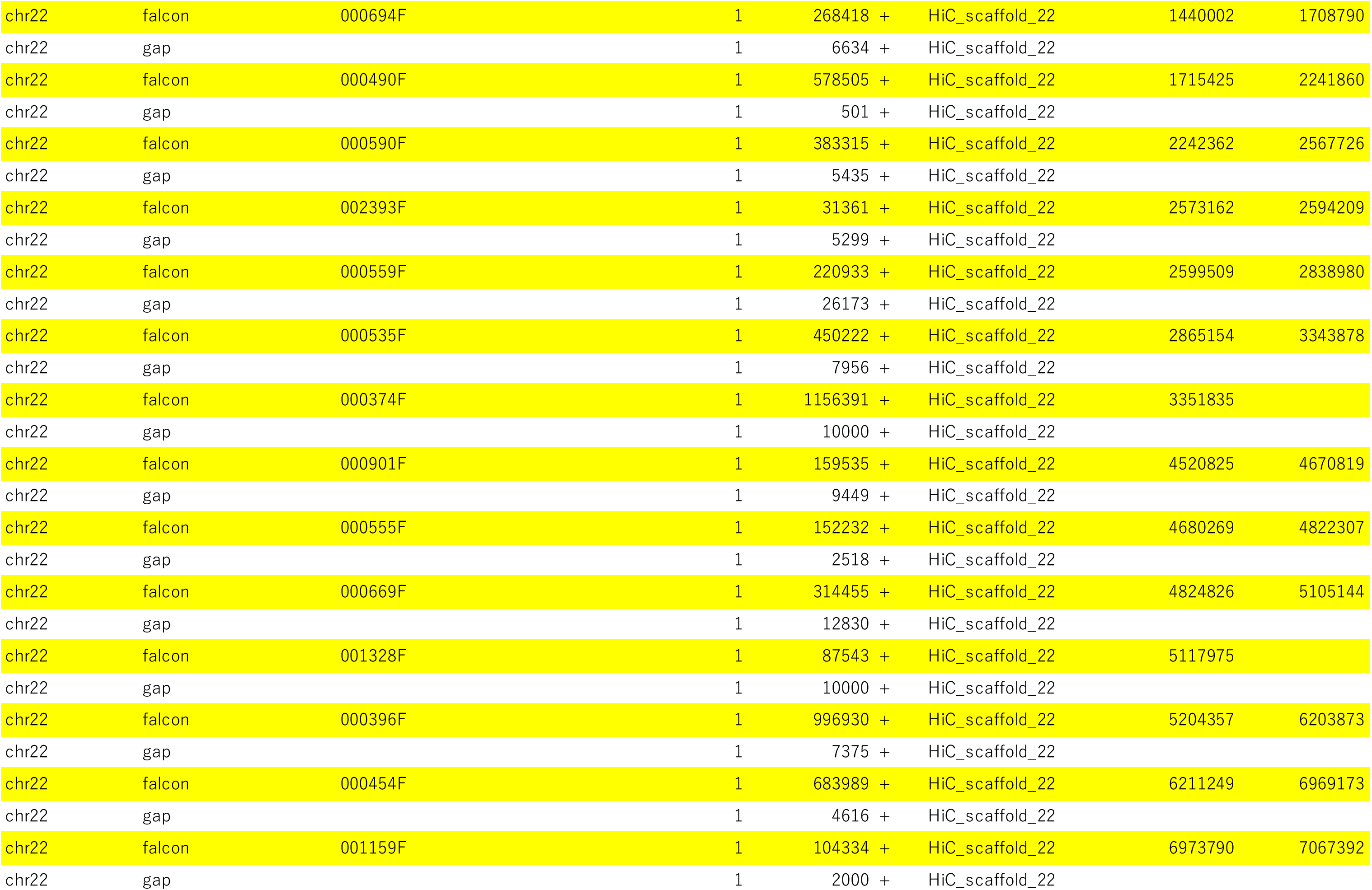
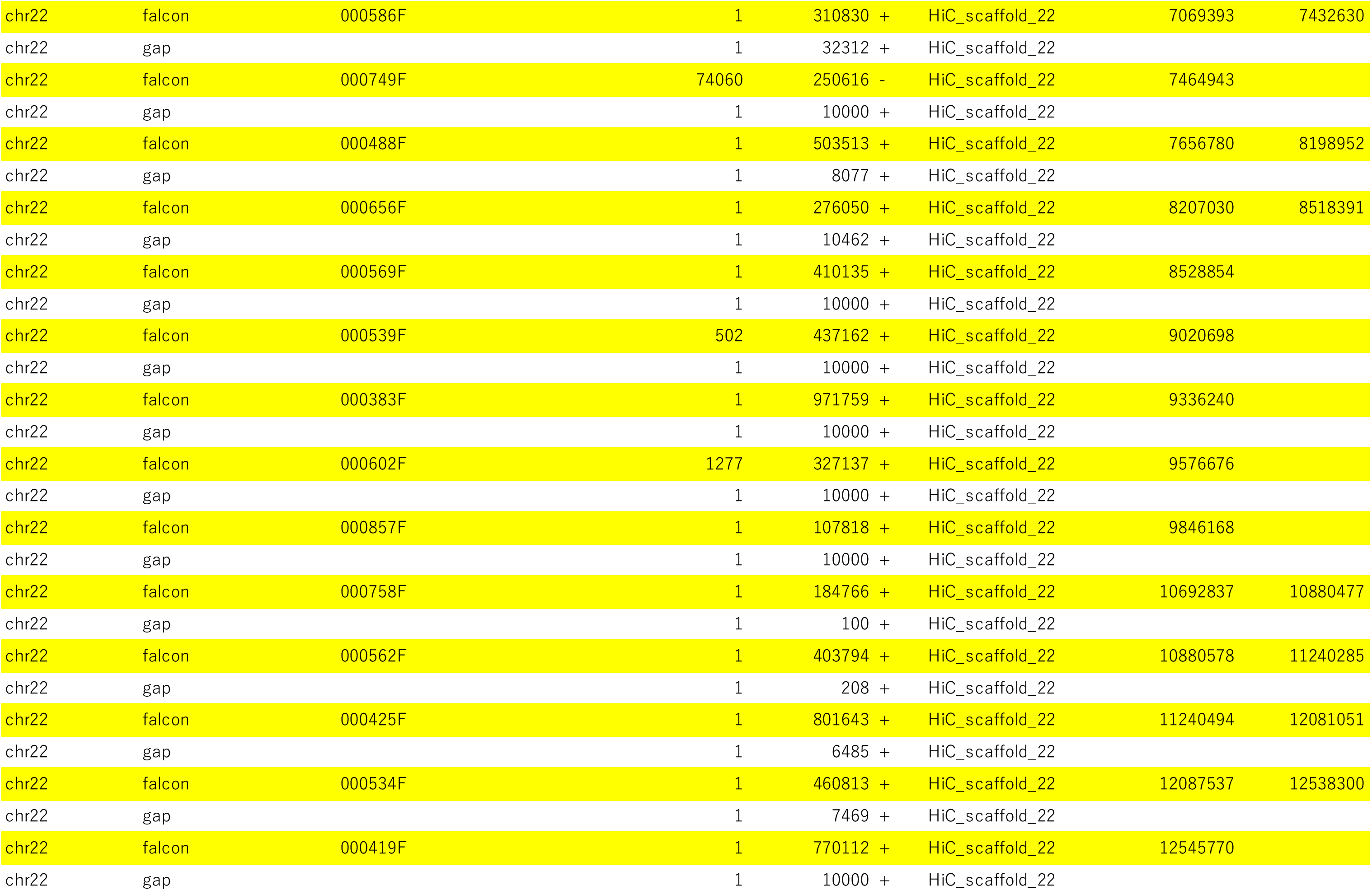
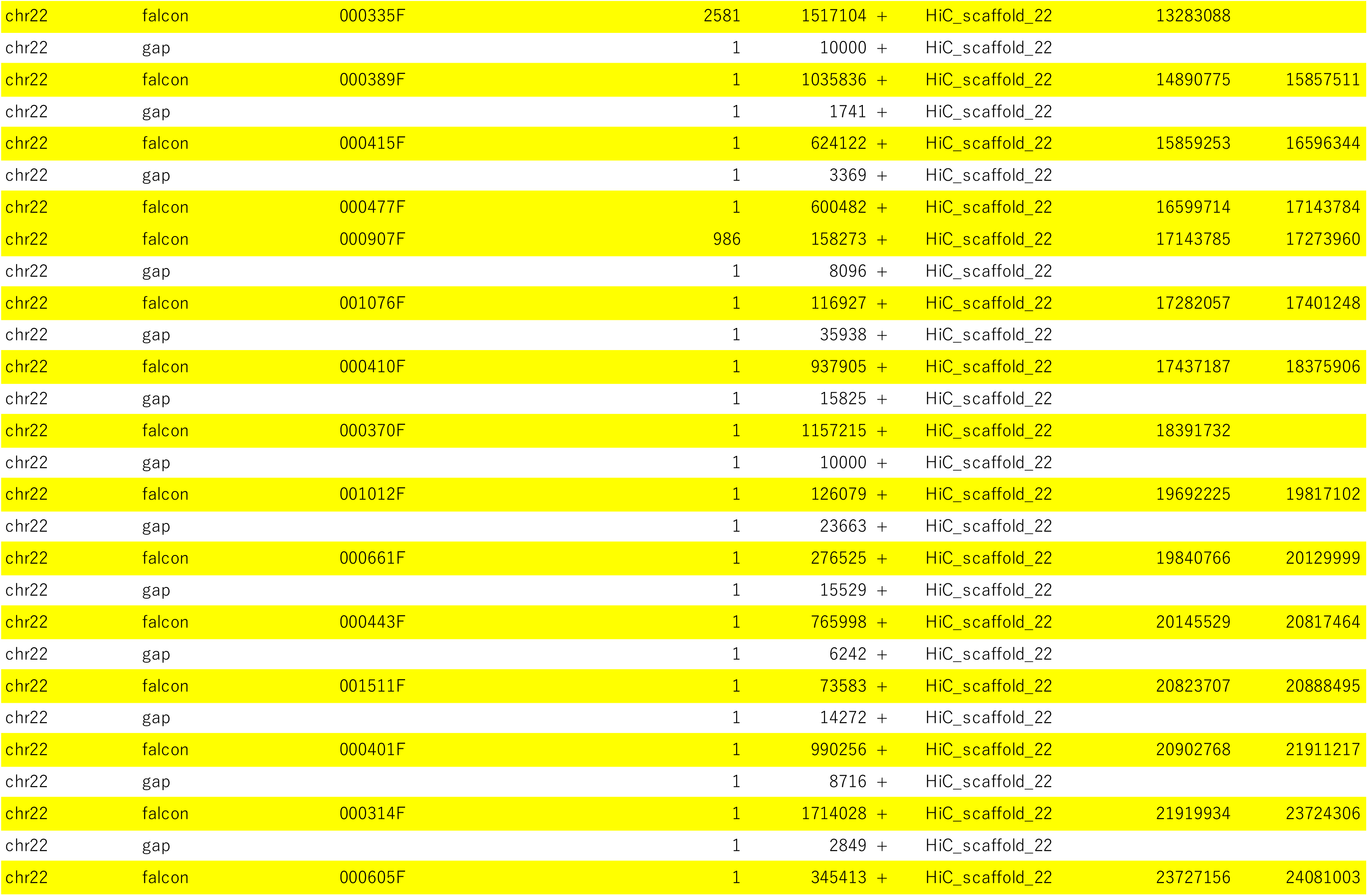
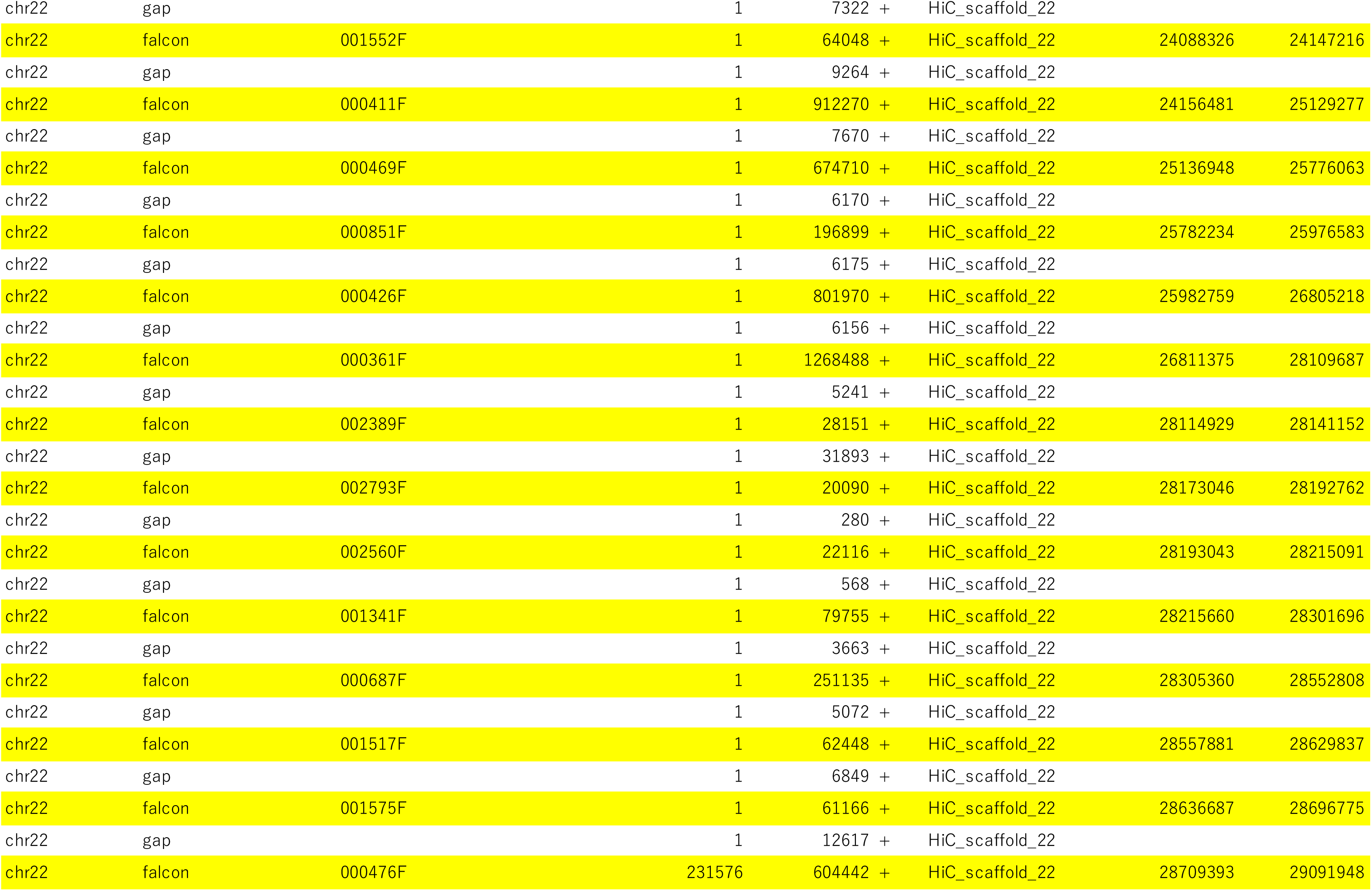
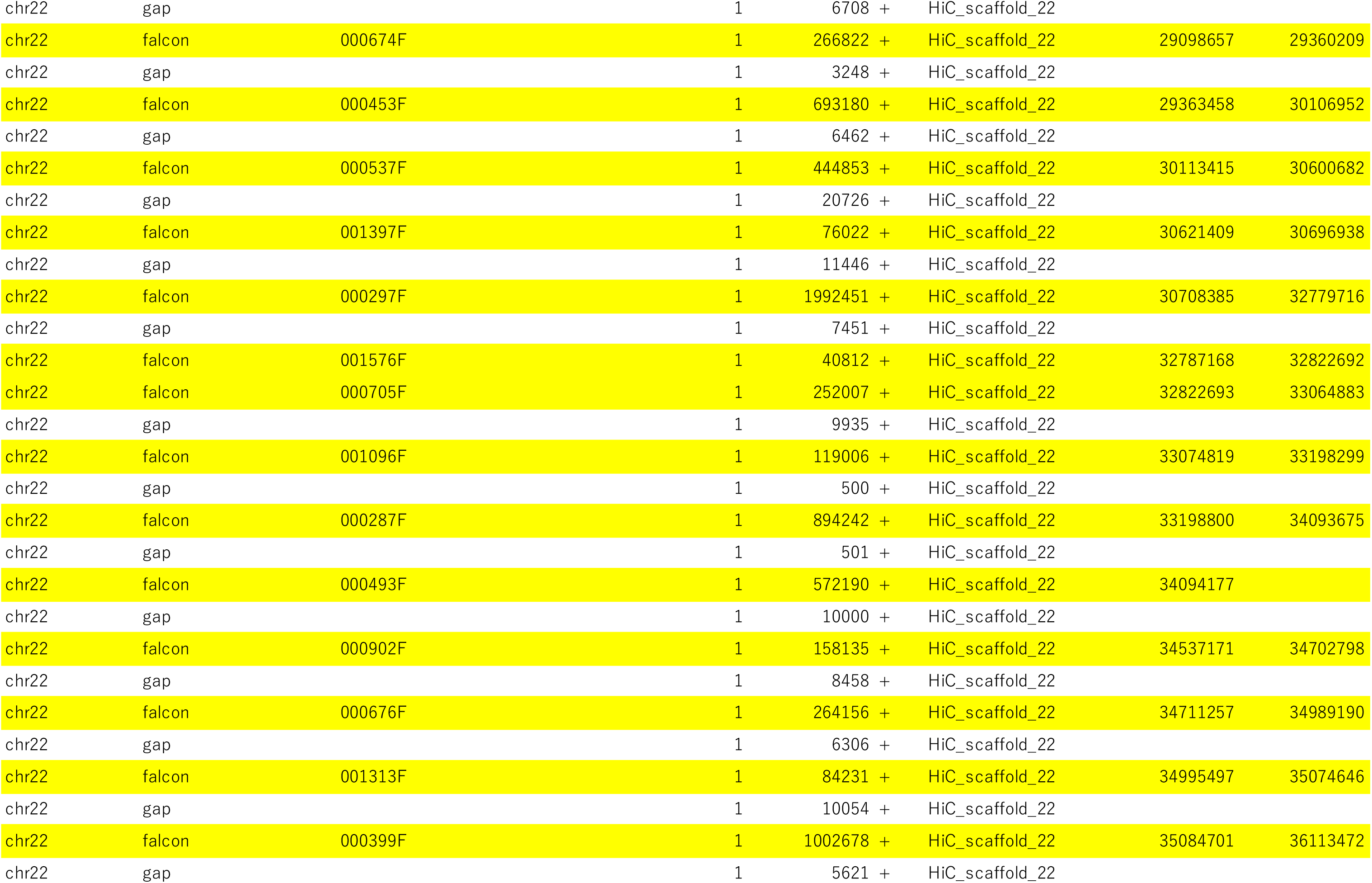
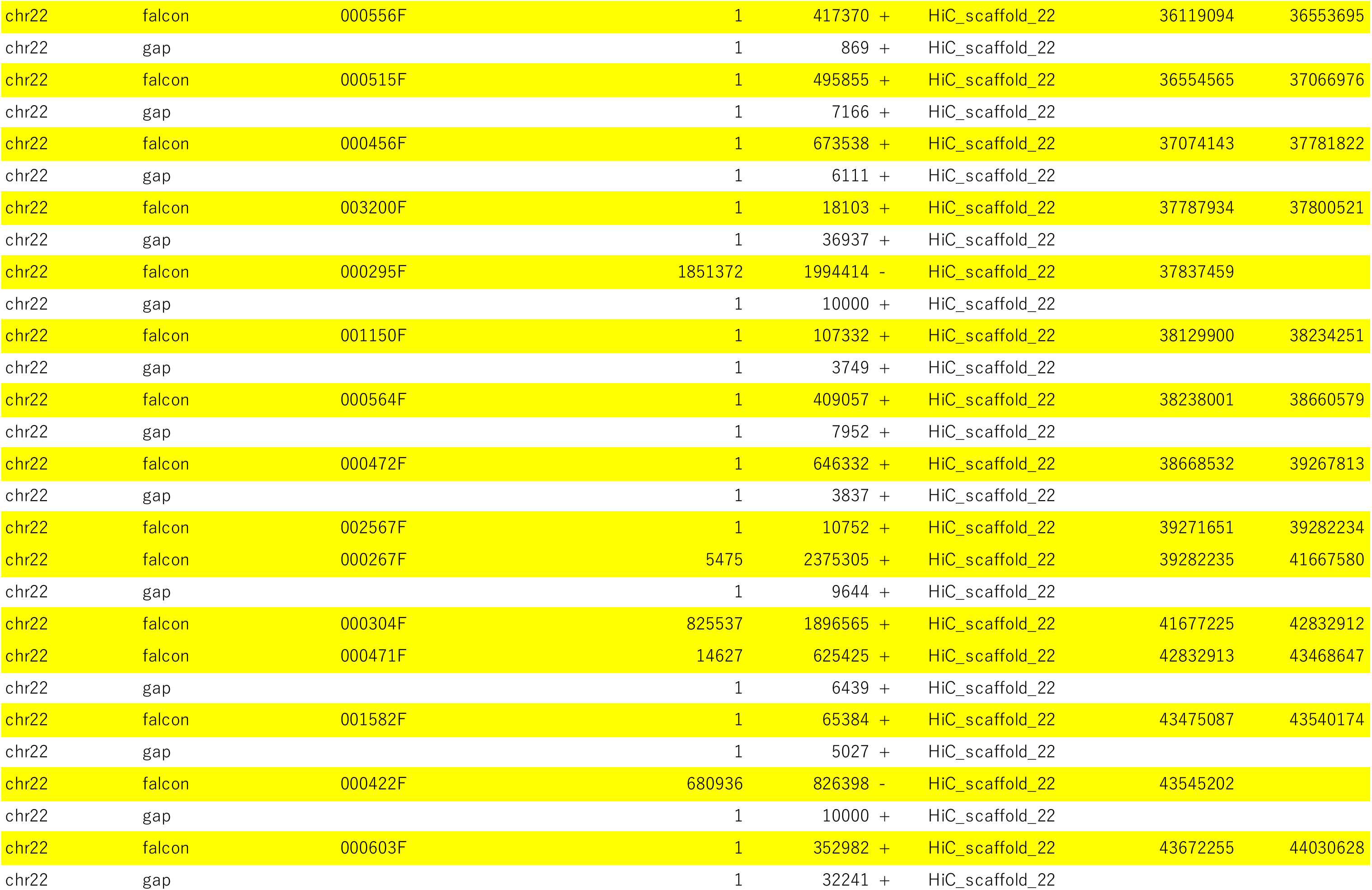
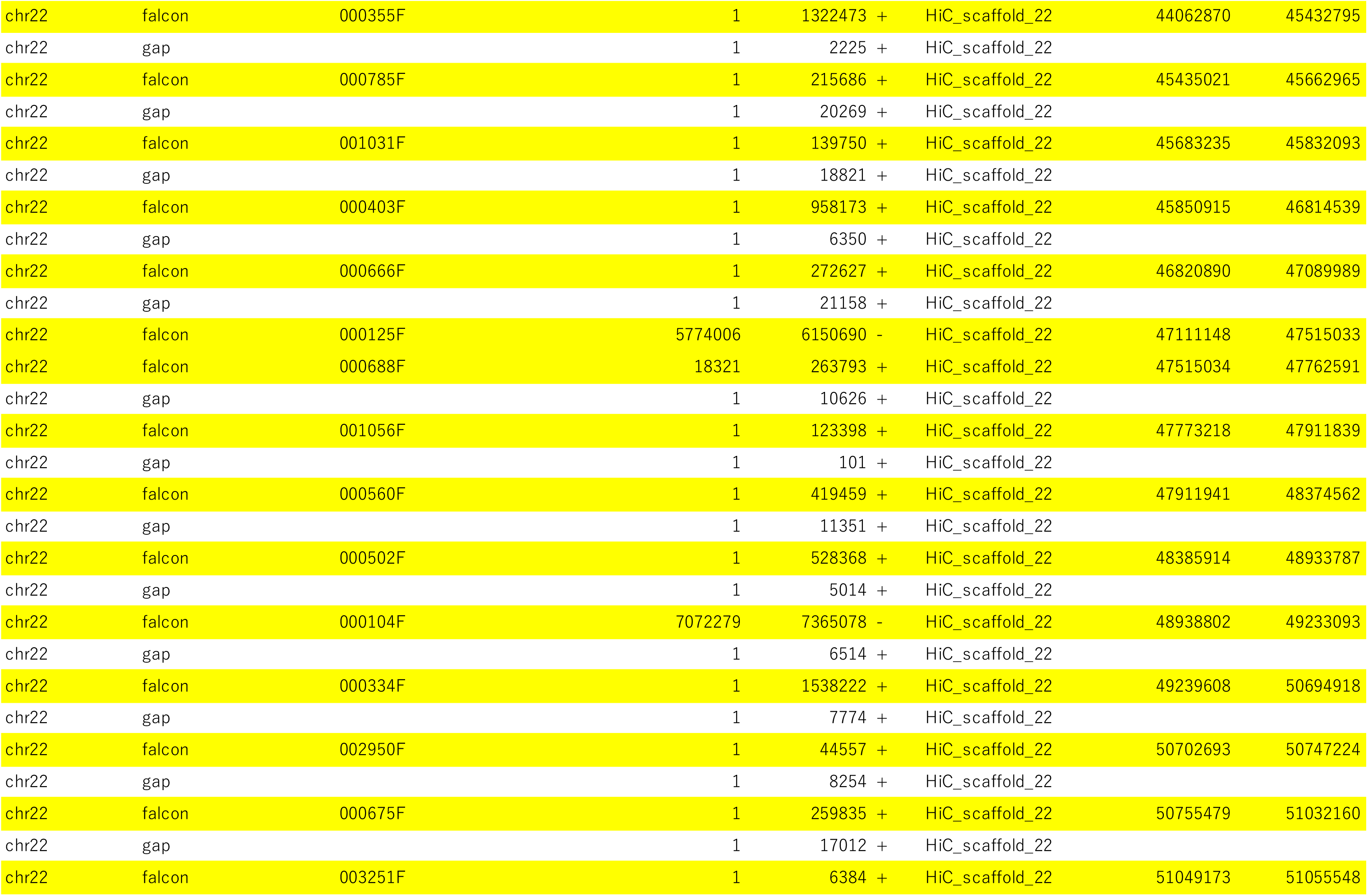
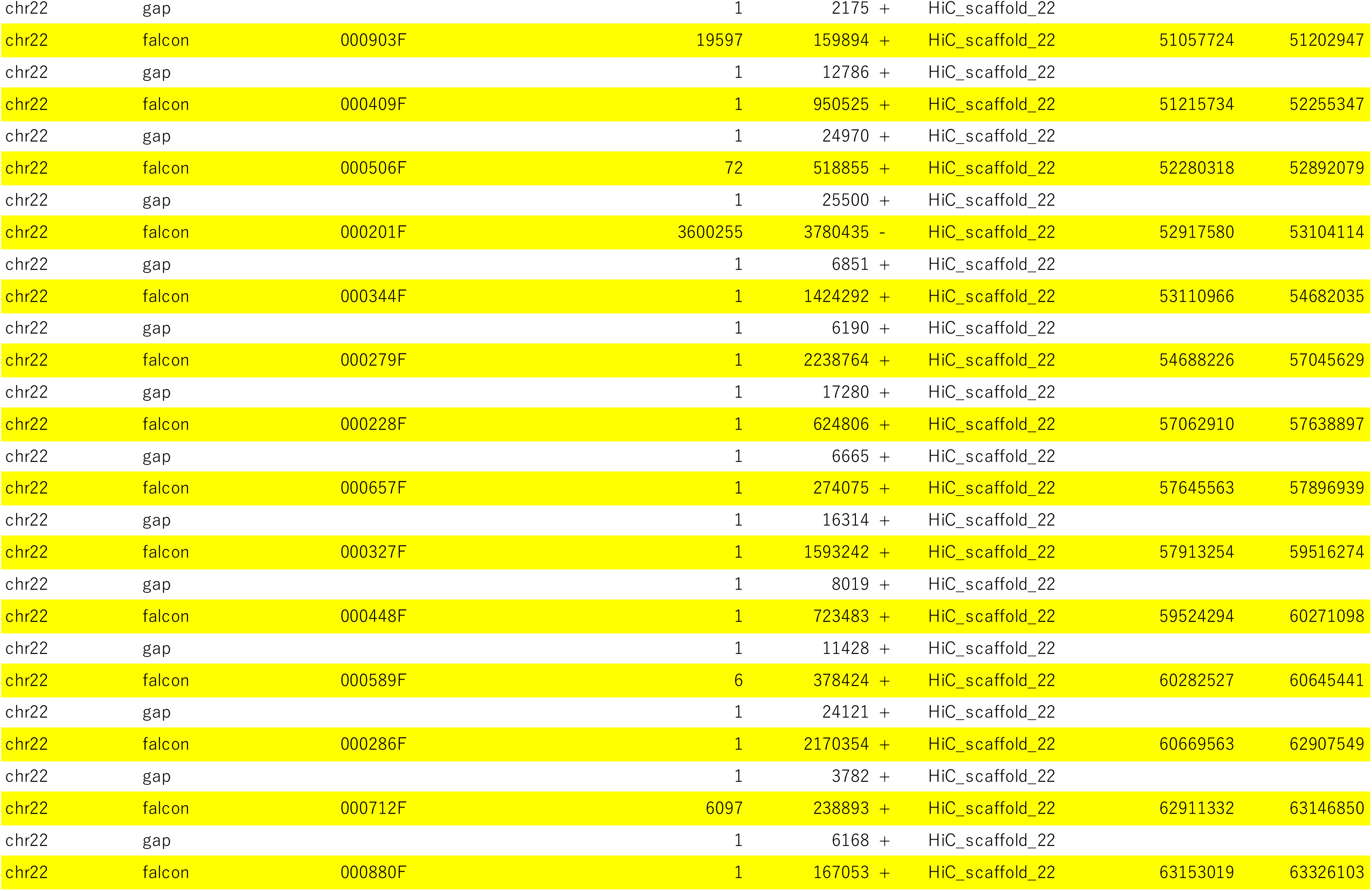
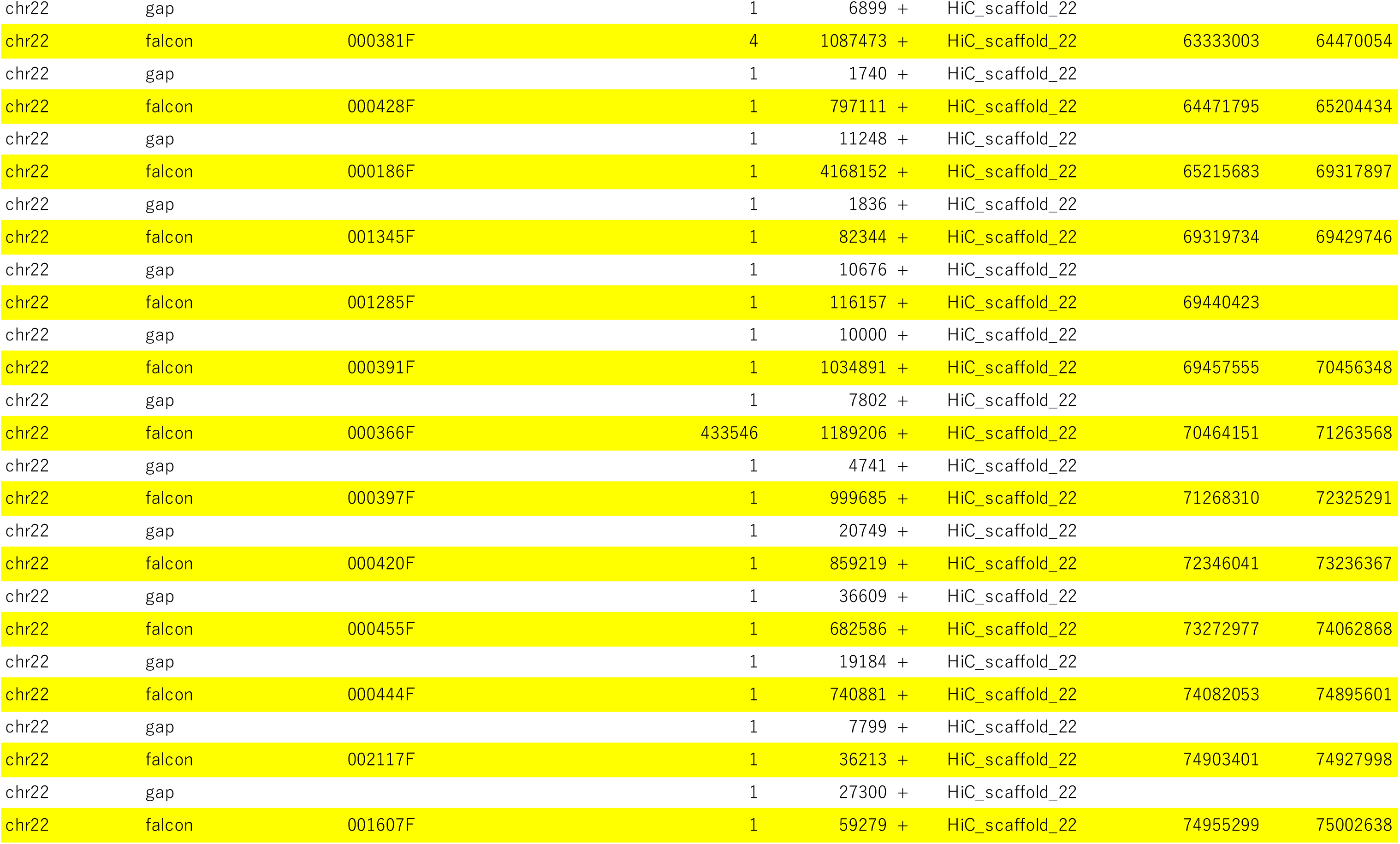
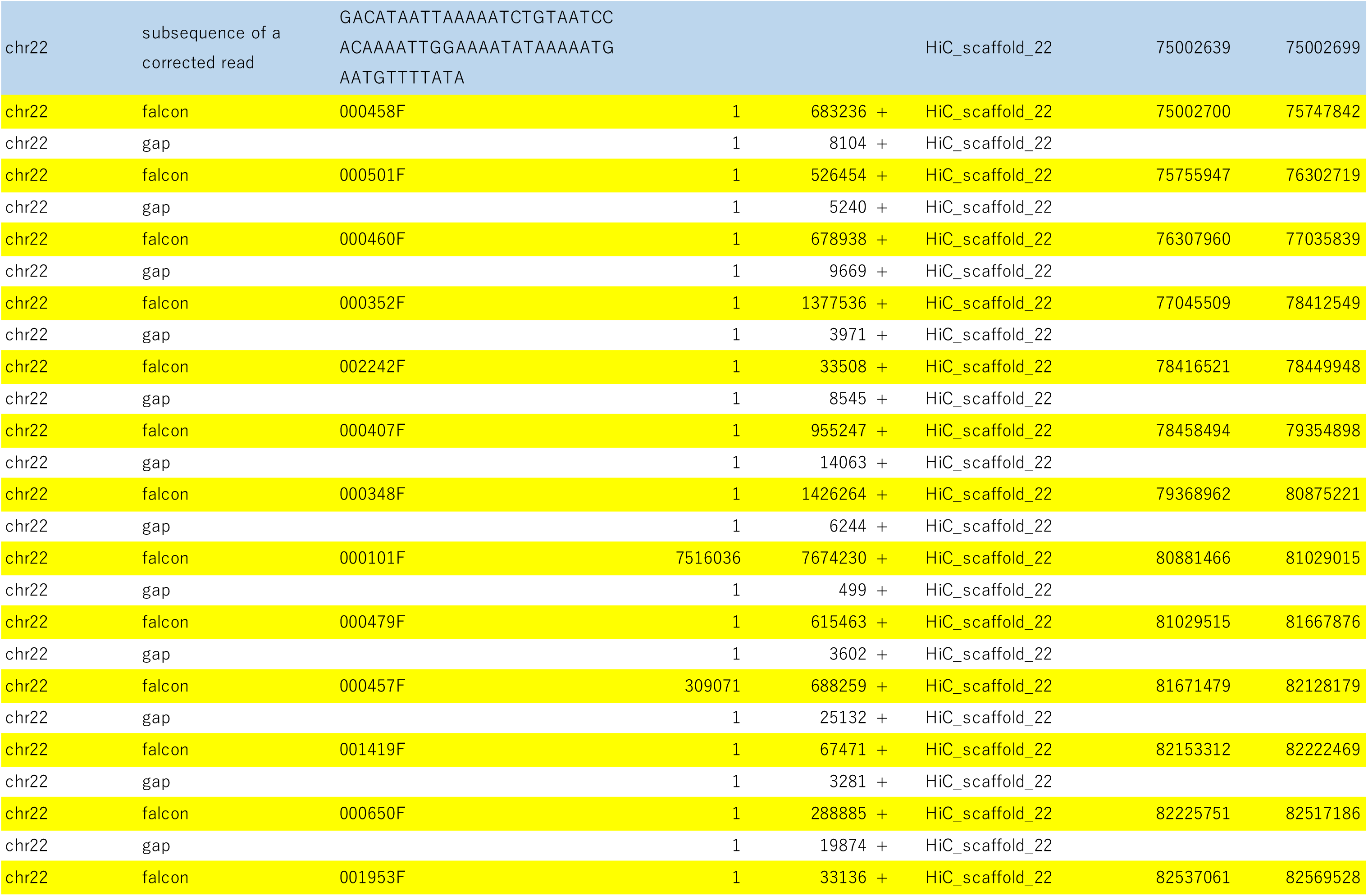
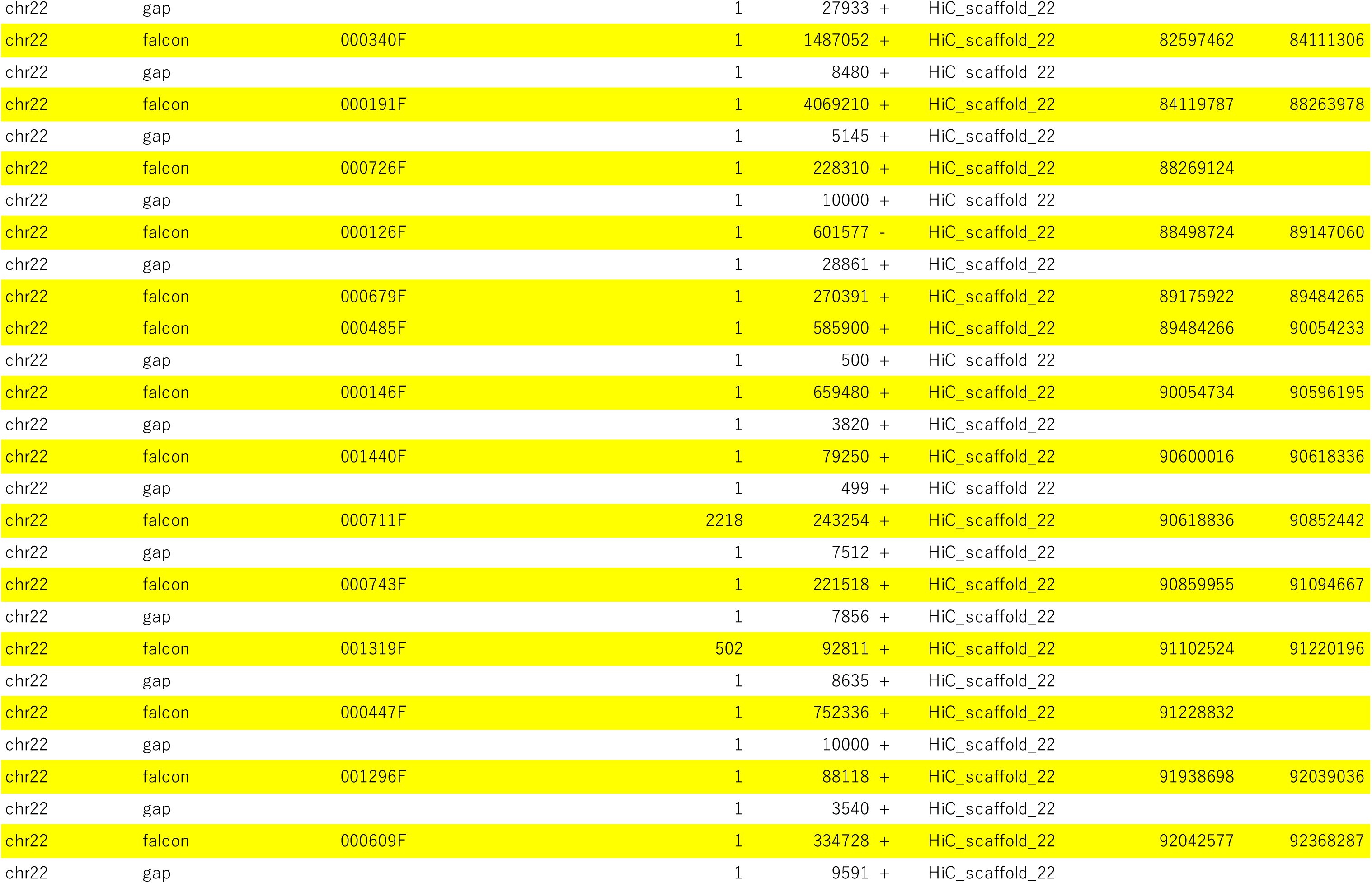
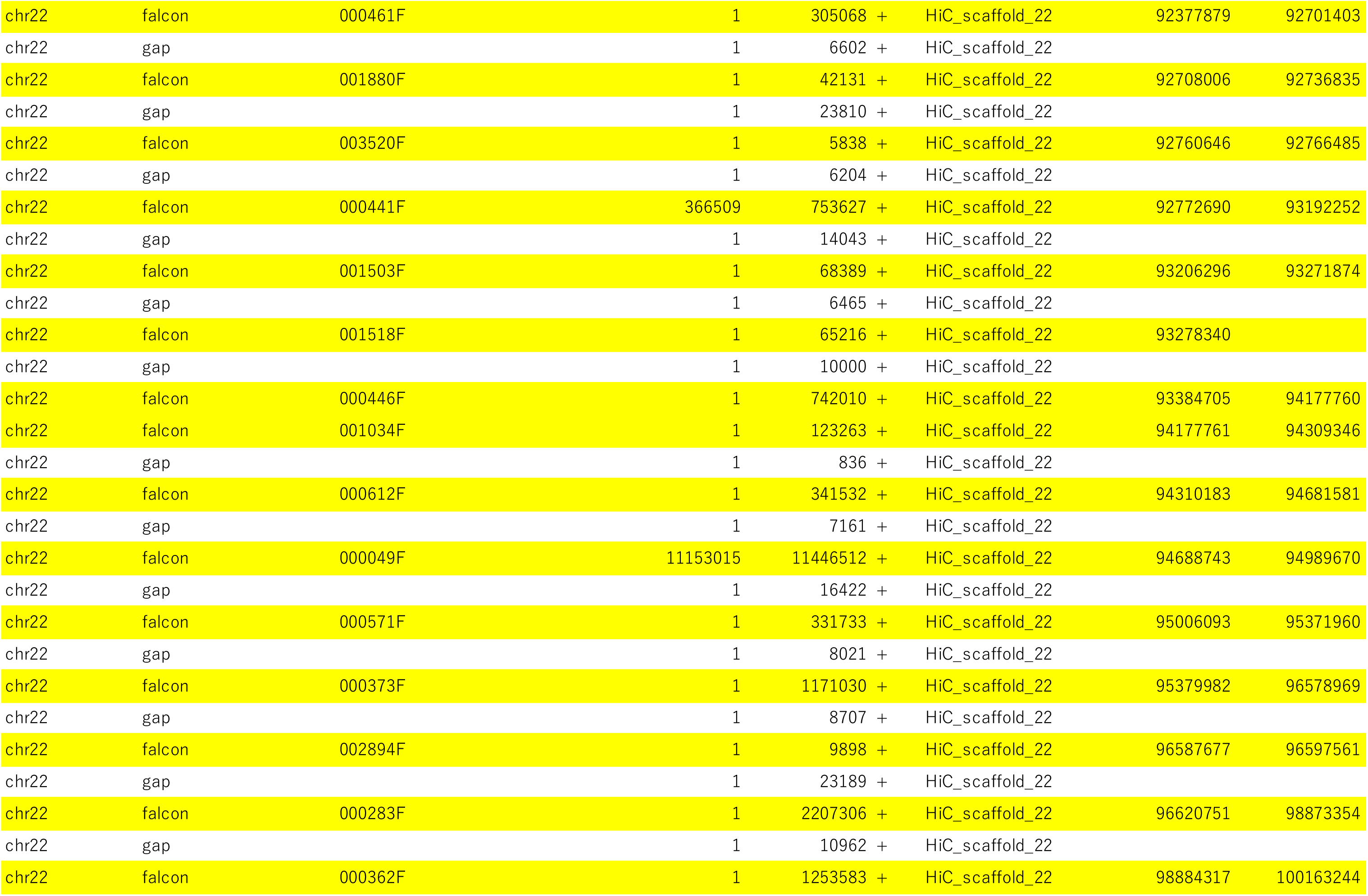
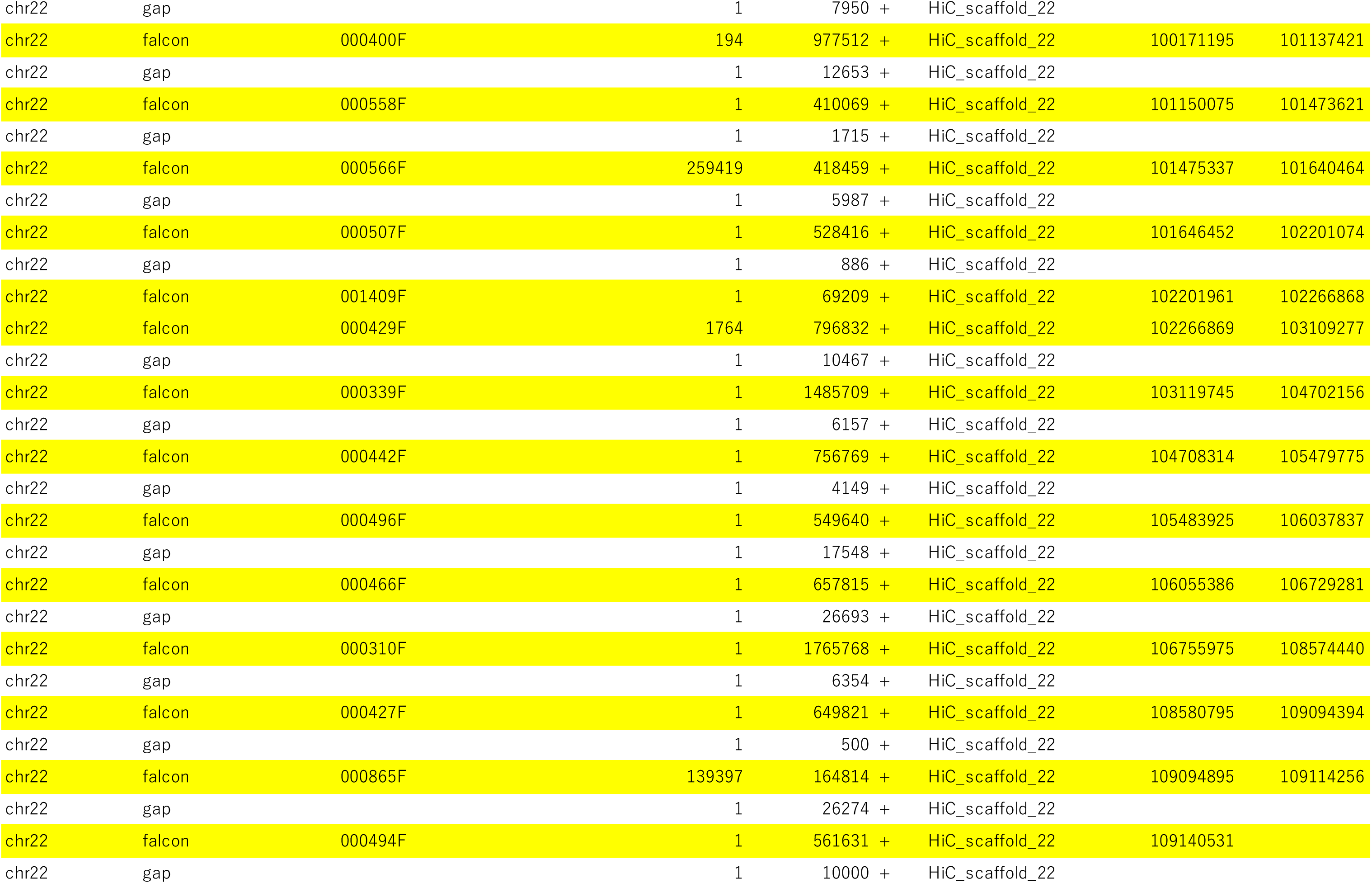
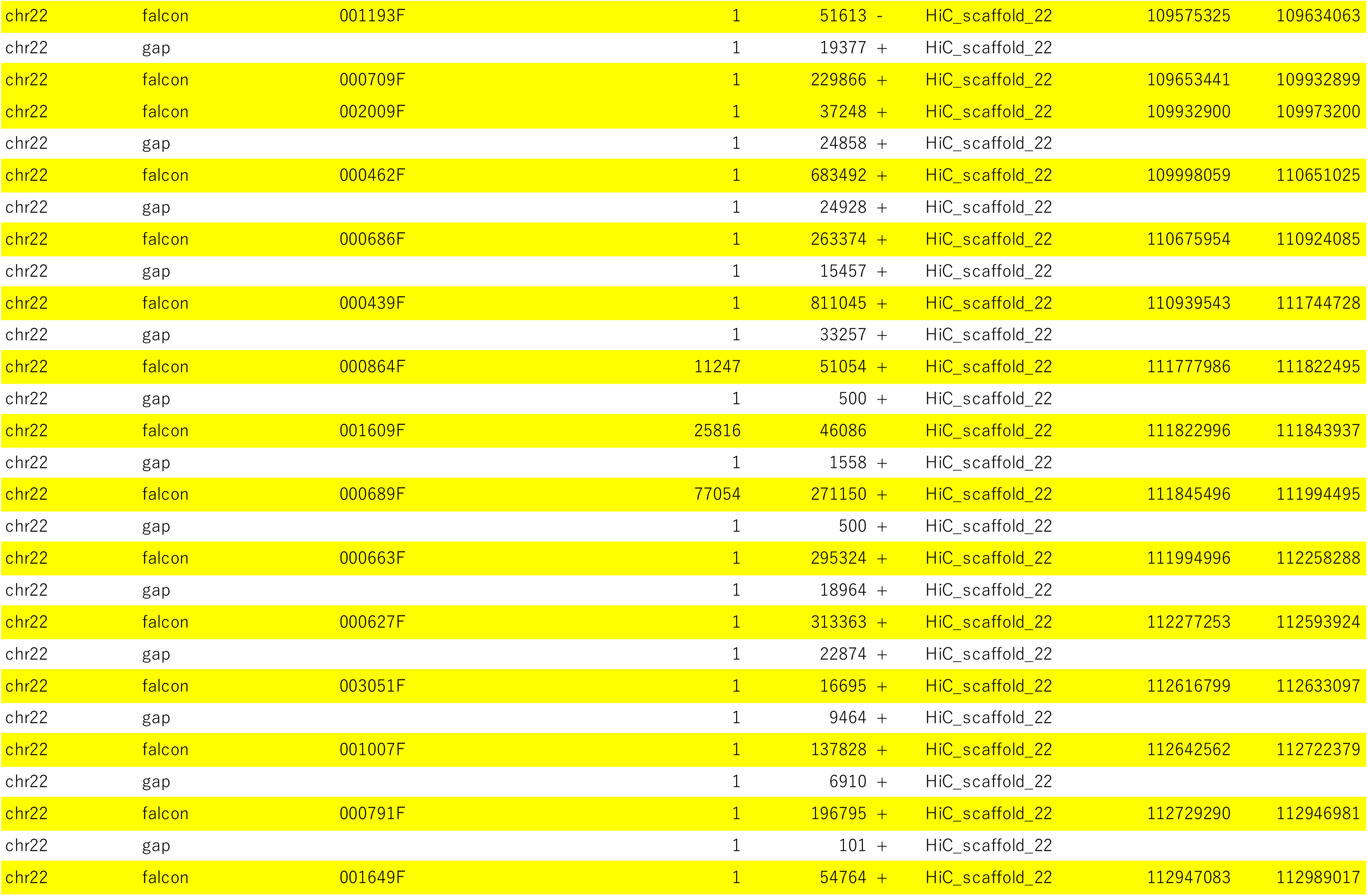
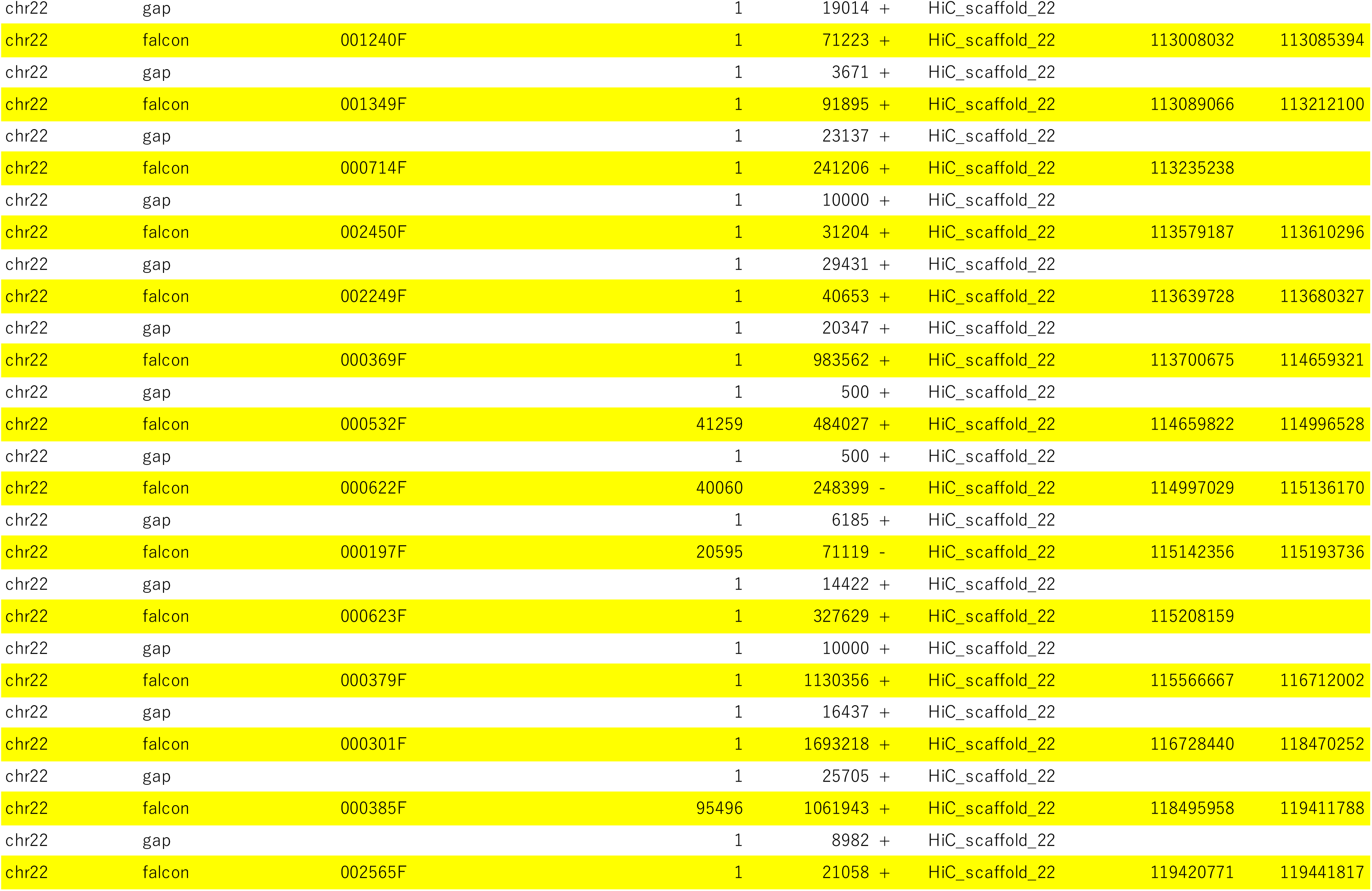
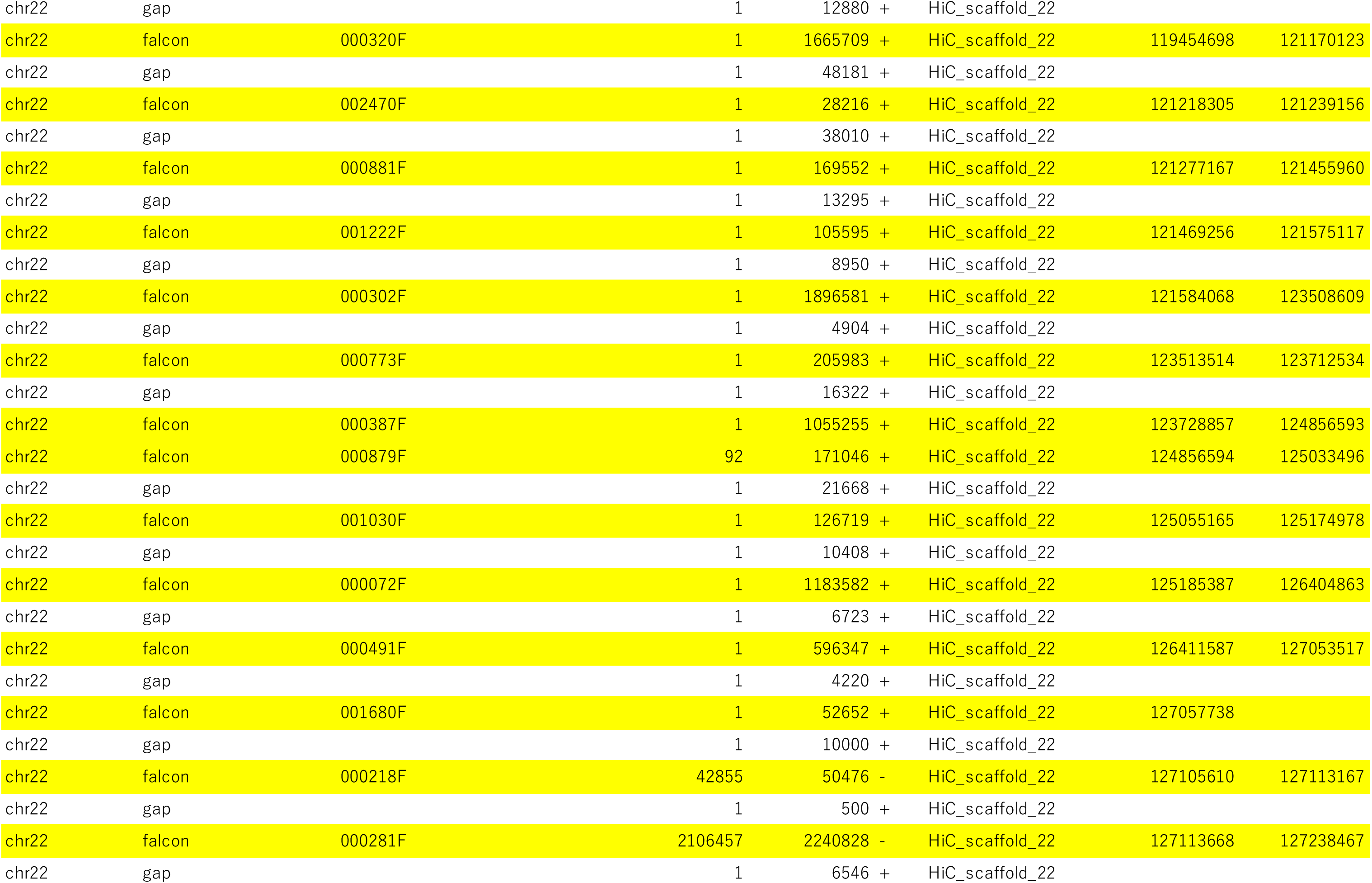
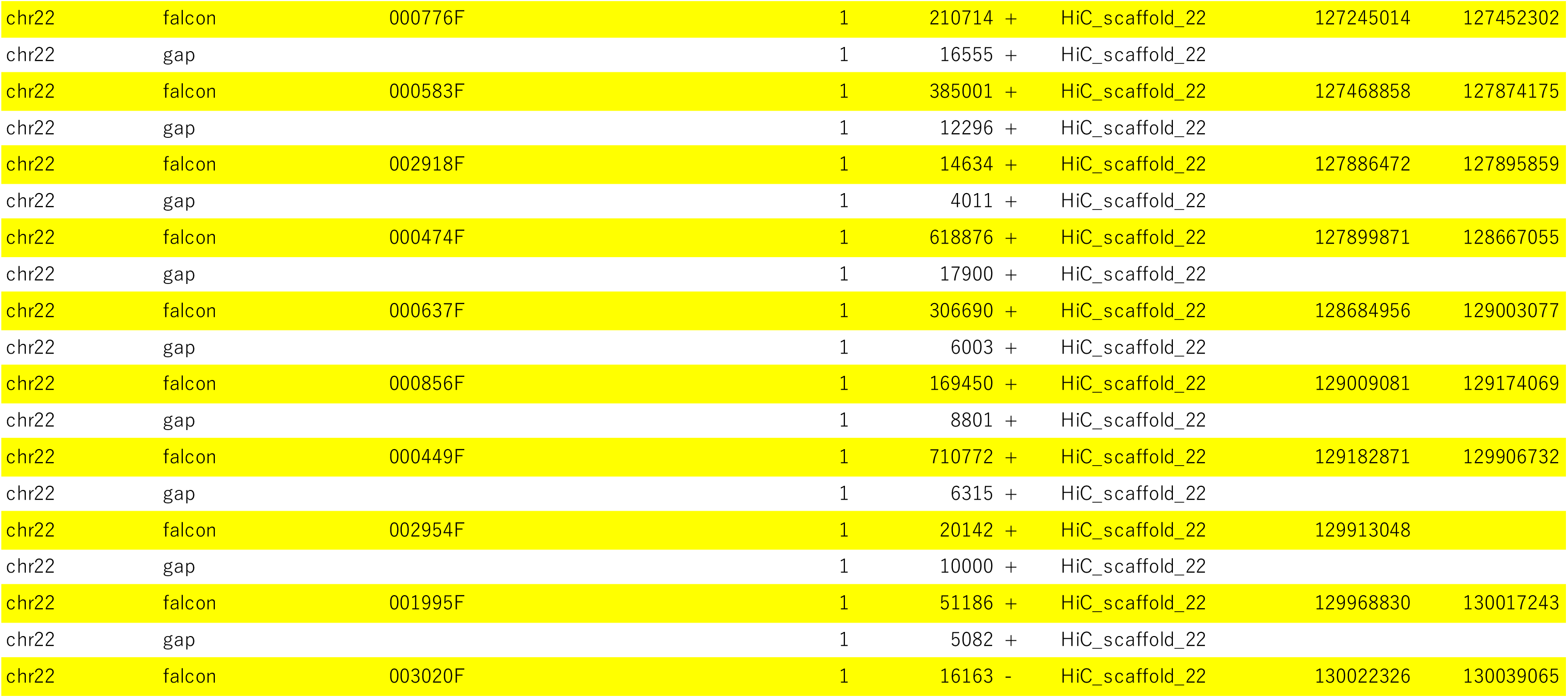
Series of connected contigs and error-corrected reads. For each chromosomal position, the first and penultimate columns list primary contigs obtained using the FALCON genome assembler (yellow), error-corrected reads (light blue), and remaining gaps (white). The fourth and fifth columns show the start and end positions for each contig, respectively.

**Table S3. Related to Table 1.** Mismatch and indel ratios of our golden hamster genome assembly before and after polishing using PacBio long reads.

Effect of polishing our golden hamster genome assembly by Arrow
|  | Before<br>polishing<br>with Arrow<br>ratio (%) | After<br>polishing<br>with Arrow<br>ratio (%) |
| --- | --- | --- |
| mismatch ratio | 0.12068% | 0.11216% |
| insertion/deletion ratio | 0.07203% | 0.02774% |
| mismatch and indel ratio | 0.19271% | 0.13990% |
| nucleotide identity | 99.80729% | 99.86010% |

**Table S4. Related to Table 2.** Proportion (%) of transposable elements in the MesAur1.0_HiC assembly.

| Class | Clade/Superfamily | MesAur1.0_HiC |
| --- | --- | --- |
| SINE | B1 | 2.71 |
|  | B2 | 1.69 |
|  | B4 | 2.27 |
|  | ID | 0.30 |
|  | MIR | 0.64 |
|  | other SINE | 0.21 |
| LINE | L1 | 11.83 |
|  | L2 | 0.50 |
|  | other LINE | 0.11 |
| LTR | ERV1 | 0.51 |
|  | ERVK | 3.18 |
|  | ERVL | 4.89 |
|  | other LTR | 0.69 |
| DNA | hAT | 0.86 |
|  | TcMar | 0.24 |
|  | other DNA | 0.03 |
| Other TE |  | 0.33 |
| Unknown TE |  | 0.08 |
| Total |  | 31.07 |

## References

Aravin, A., Gaidatzis, D., Pfeffer, S., Lagos-Quintana, M., Landgraf, P., Iovino, N., Morris, P., Brownstein, M.J., Kuramochi-Miyagawa, S., Nakano, T., et al. (2006). A novel class of small RNAs bind to MILI protein in mouse testes. Nature 442, 203–207.

Aravin, A.A., Sachidanandam, R., Bourc’his, D., Schaefer, C., Pezic, D., Toth, K.F., Bestor, T., and Hannon, G.J. (2008). A piRNA pathway primed by individual transposons is linked to de novo DNA methylation in mice. Mol Cell 31, 785–799.

Aravin, A.A., Sachidanandam, R., Girard, A., Fejes-Toth, K., and Hannon, G.J. (2007). Developmentally regulated piRNA clusters implicate MILI in transposon control. Science 316, 744–747.

Bailey, T.L., Boden, M., Buske, F.A., Frith, M., Grant, C.E., Clementi, L., Ren, J., Li, W.W., and Noble, W.S. (2009). MEME SUITE: tools for motif discovery and searching. Nucleic Acids Res 37, W202–208.

Baronti, L., Guzzetti, I., Ebrahimi, P., Friebe Sandoz, S., Steiner, E., Schlagnitweit, J., Fromm, B., Silva, L., Fontana, C., Chen, A.A., et al. (2020). Base-pair conformational switch modulates miR-34a targeting of Sirt1 mRNA. Nature 583, 139–144.

Brennecke, J., Aravin, A.A., Stark, A., Dus, M., Kellis, M., Sachidanandam, R., and Hannon, G.J. (2007). Discrete small RNA-generating loci as master regulators of transposon activity in Drosophila. Cell 128, 1089–1103.

Brind’Amour, J., Kobayashi, H., Richard Albert, J., Shirane, K., Sakashita, A., Kamio, A., Bogutz, A., Koike, T., Karimi, M.M., Lefebvre, L., et al. (2018). LTR retrotransposons transcribed in oocytes drive species-specific and heritable changes in DNA methylation. Nat Commun 9, 3331.

Carmell, M.A., Girard, A., van de Kant, H.J., Bourc’his, D., Bestor, T.H., de Rooij, D.G., and Hannon, G.J. (2007). MIWI2 is essential for spermatogenesis and repression of transposons in the mouse male germline. Dev Cell 12, 503–514.

Chin, C.S., Peluso, P., Sedlazeck, F.J., Nattestad, M., Concepcion, G.T., Clum, A., Dunn, C., O’Malley, R., Figueroa-Balderas, R., Morales-Cruz, A., et al. (2016). Phased diploid genome assembly with single-molecule real-time sequencing. Nat Methods 13, 1050–1054.

Chuong, E.B., Elde, N.C., and Feschotte, C. (2017). Regulatory activities of transposable elements: from conflicts to benefits. Nat Rev Genet 18, 71–86.

De Fazio, S., Bartonicek, N., Di Giacomo, M., Abreu-Goodger, C., Sankar, A., Funaya, C., Antony, C., Moreira, P.N., Enright, A.J., and O’Carroll, D. (2011). The endonuclease activity of Mili fuels piRNA amplification that silences LINE1 elements. Nature 480, 259–263.

Deng, W. & Lin, H. (2002). Miwi, a murine homolog of piwi, encodes a cytoplasmic protein essential for spermatogenesis. Dev. Cell 2, 819–830.

Ding D, Liu J, Dong K, Midic U, Hess RA, Xie H, Demireva EY, Chen C. (2017). PNLDC1 is essential for piRNA 3’ end trimming and transposon silencing during spermatogenesis in mice. Nat Commun. 8, 819

Elkayam, E., Kuhn, C.D., Tocilj, A., Haase, A.D., Greene, E.M., Hannon, G.J., and Joshua-Tor, L. (2012). The structure of human argonaute-2 in complex with miR-20a. Cell 150, 100–110.

Ernst, C., Odom, D.T., and Kutter, C. (2017). The emergence of piRNAs against transposon invasion to preserve mammalian genome integrity. Nat Commun 8, 1411.

Fan, Z., Li, W., Lee, S.R., Meng, Q., Shi, B., Bunch, T.D., White, K.L., Kong, I.K., and Wang, Z. (2014). Efficient gene targeting in golden Syrian hamsters by the CRISPR/Cas9 system. PLoS One 9, e109755.

Flynn, J.M., Hubley, R., Goubert, C., Rosen, J., Clark, A.G., Feschotte, C., and Smit, A.F. (2020). RepeatModeler2 for automated genomic discovery of transposable element families. Proc Natl Acad Sci U S A 117, 9451–9457.

Franke, V., Ganesh, S., Karlic, R., Malik, R., Pasulka, J., Horvat, F., Kuzman, M., Fulka, H., Cernohorska, M., Urbanova, J., et al. (2017). Long terminal repeats power evolution of genes and gene expression programs in mammalian oocytes and zygotes. Genome Res 27, 1384–1394.

Friedländer, M.R., Mackowiak, S.D., Li, N., Chen, W., and Rajewsky, N. (2012). miRDeep2 accurately identifies known and hundreds of novel microRNA genes in seven animal clades. Nucleic Acids Res 40, 37–52.

Gainetdinov, I., Colpan, C., Arif, A., Cecchini, K., and Zamore, P.D. (2018). A Single Mechanism of Biogenesis, Initiated and Directed by PIWI Proteins, Explains piRNA Production in Most Animals. Mol Cell 71, 775–790.e775.

Girard, A., Sachidanandam, R., Hannon, G.J., and Carmell, M.A. (2006). A germline-specific class of small RNAs binds mammalian Piwi proteins. Nature 442, 199–202.

Gonzalez, J., Qi, H., Liu, N., and Lin, H. (2015). Piwi is a key rgulator of both somatic and germline stem cells in the *Drosophila* testis. Cell Rep 12, 150–161.

Gunawardane, L.S., Saito, K., Nishida, K.M., Miyoshi, K., Kawamura, Y., Nagami, T., Siomi, H., and Siomi, M.C. (2007). A slicer-mediated mechanism for repeat-associated siRNA 5’ end formation in Drosophila. Science 315, 1587–1590.

Gurevich, A., Saveliev, V., Vyahhi, N., and Tesler, G. (2013). QUAST: quality assessment tool for genome assemblies. Bioinformatics 29, 1072–1075.

Haase, A. D., Fenoglio, S., Muerdter, F., Guzzardo, P. M., Czech, B., Pappin, D. J., Chen, C., Gordon, A., and Hannon, G. J. (2010). Probing the initiation and effector phases of the somatic piRNA pathway in *Drosophila*. Genes Dev 24, 2499–2504.

Han, B.W., Wang, W., Li, C., Weng, Z., and Zamore, P.D. (2015). Noncoding RNA. piRNA-guided transposon cleavage initiates Zucchini-dependent, phased piRNA production. Science 348, 817–821.

Han, J.S., and Boeke, J.D. (2005). LINE-1 retrotransposons: modulators of quantity and quality of mammalian gene expression? Bioessays 27, 775–784.

Hancks, D.C., and Kazazian, H.H., Jr. (2016). Roles for retrotransposon insertions in human disease. Mob DNA 7, 9.

Hayashi, R., Schnabl, J., Handler, D., Mohn, F., Ameres, S.L., and Brennecke, J. (2016). Genetic and mechanistic diversity of piRNA 3’-end formation. Nature 539, 588–592.

Heinz, S., Benner, C., Spann, N., Bertolino, E., Lin, Y.C., Laslo, P., Cheng, J.X., Murre, C., Singh, H., and Glass, C.K. (2010). Simple combinations of lineage-determining transcription factors prime cis-regulatory elements required for macrophage and B cell identities. Mol Cell 38, 576–589.

Hirano, T., Iwasaki, Y.W., Lin, Z.Y., Imamura, M., Seki, N.M., Sasaki, E., Saito, K., Okano, H., Siomi, M.C., and Siomi, H. (2014). Small RNA profiling and characterization of piRNA clusters in the adult testes of the common marmoset, a model primate. Rna 20, 1223–1237.

Hirose, M., Honda, A., Fulka, H., Tamura-Nakano, M., Matoba, S., Tomishima, T., Mochida, K., Hasegawa, A., Nagashima, K., Inoue, K., et al. (2020). Acrosin is essential for sperm penetration through the zona pellucida in hamsters. Proc Natl Acad Sci U S A 117, 2513–2518.

Hirose, M., and Ogura, A. (2019). The golden (Syrian) hamster as a model for the study of reproductive biology: Past, present, and future. Reprod Med Biol 18, 34–39.

Horwich, M.D., Li, C., Matranga, C., Vagin, V., Farley, G., Wang, P., and Zamore, P.D. (2007). The Drosophila RNA methyltransferase, DmHen1, modifies germline piRNAs and single-stranded siRNAs in RISC. Curr Biol 17, 1265–1272.

Ipsaro, J.J., Haase, A.D., Knott, S.R., Joshua-Tor, L., and Hannon, G.J. (2012). The structural biochemistry of Zucchini implicates it as a nuclease in piRNA biogenesis. Nature 491, 279–283.

Ishizuka, A., Siomi, M.C., and Siomi, H. (2002). A Drosophila fragile X protein interacts with components of RNAi and ribosomal proteins. Genes Dev 16, 2497–2508.

Iwasaki, Y.W., Ishino, K., and Siomi, H. (2017). Deep sequencing and high-throughput analysis of PIWI-associated small RNAs. Methods 126, 66–75.

Iwasaki, Y.W., Siomi, M.C., and Siomi, H. (2015). PIWI-Interacting RNA: Its Biogenesis and Functions. Annu Rev Biochem 84, 405–433.

Izumi, N., Shoji, K., Sakaguchi, Y., Honda, S., Kirino, Y., Suzuki, T., Katsuma, S., and Tomari, Y. (2016). Identification and Functional Analysis of the Pre-piRNA 3’ Trimmer in Silkworms. Cell 164, 962–973.

Jurka, J., Kapitonov, V.V., Pavlicek, A., Klonowski, P., Kohany, O., and Walichiewicz, J. (2005). Repbase Update, a database of eukaryotic repetitive elements. Cytogenet Genome Res 110, 462–467.

Kabayama, Y., Toh, H., Katanaya, A., Sakurai, T., Chuma, S., Kuramochi-Miyagawa, S., Saga, Y., Nakano, T., and Sasaki, H. (2017). Roles of MIWI, MILI and PLD6 in small RNA regulation in mouse growing oocytes. Nucleic Acids Res 45, 5387–5398.

Katoh, K., and Standley, D.M. (2013). MAFFT multiple sequence alignment software version 7: improvements in performance and usability. Mol Biol Evol 30, 772–780.

Kaya, E., Doxzen, K.W., Knoll, K.R., Wilson, R.C., Strutt, S.C., Kranzusch, P.J., and Doudna, J.A. (2016). A bacterial Argonaute with noncanonical guide RNA specificity. Proc Natl Acad Sci U S A 113, 4057–4062.

Kent, W.J., Baertsch, R., Hinrichs, A., Miller, W., and Haussler, D. (2003). Evolution’s cauldron: duplication, deletion, and rearrangement in the mouse and human genomes. Proc Natl Acad Sci U S A 100, 11484–11489.

Khurana, J.S., Wang, J., Xu, J., Koppetsch, B.S., Thomson, T.C., Nowosielska, A., Li, C., Zamore, P.D., Weng, Z., and Theurkauf, W.E. (2011). Adaptation to P element transposon invasion in Drosophila melanogaster. Cell 147, 1551–1563.

Kim, D., Langmead, B., and Salzberg, S.L. (2015). HISAT: a fast spliced aligner with low memory requirements. Nat Methods 12, 357–360.

Kim, D., Pertea, G., Trapnell, C., Pimentel, H., Kelley, R., and Salzberg, S.L. (2013). TopHat2: accurate alignment of transcriptomes in the presence of insertions, deletions and gene fusions. Genome Biol 14, R36.

Kirino, Y., and Mourelatos, Z. (2007). The mouse homolog of HEN1 is a potential methylase for Piwi-interacting RNAs. Rna 13, 1397–1401.

Kobayashi, H., Shoji, K., Kiyokawa, K., Negishi, L., and Tomari, Y. (2019). Iruka Eliminates Dysfunctional Argonaute by Selective Ubiquitination of Its Empty State. Mol Cell 73, 119–129.e5.

Kojima, K.K. (2018). Human transposable elements in Repbase: genomic footprints from fish to humans. Mob DNA 9, 2.

Kuramochi-Miyagawa, S., Watanabe, T., Gotoh, K., Takamatsu, K., Chuma, S., Kojima-Kita, K., Shiromoto, Y., Asada, N., Toyoda, A., Fujiyama, A., et al. (2010). MVH in piRNA processing and gene silencing of retrotransposons. Genes Dev 24, 887–892.

Kuramochi-Miyagawa, S., Watanabe, T., Gotoh, K., Totoki, Y., Toyoda, A., Ikawa, M., Asada, N., Kojima, K., Yamaguchi, Y., Ijiri, T.W., et al. (2008). DNA methylation of retrotransposon genes is regulated by Piwi family members MILI and MIWI2 in murine fetal testes. Genes Dev 22, 908–917.

Langmead, B., Trapnell, C., Pop, M., and Salzberg, S.L. (2009). Ultrafast and memory-efficient alignment of short DNA sequences to the human genome. Genome Biol 10, R25.

Lau, N.C., Seto, A.G., Kim, J., Kuramochi-Miyagawa, S., Nakano, T., Bartel, D.P., and Kingston, R.E. (2006). Characterization of the piRNA complex from rat testes. Science 313, 363–367.

Li, F., Yuan, P., Rao, M., Jin, C.H., Tang, W., Rong, Y.F., Hu, Y.P., Zhang, F., Wei, T., Yin, Q., et al. (2020). piRNA-independent function of PIWIL1 as a co-activator for anaphase promoting complex/cyclosome to drive pancreatic cancer metastasis. Nat Cell Biol 22, 425–438.

Li, H. (2018). Minimap2: pairwise alignment for nucleotide sequences. Bioinformatics 34, 3094–3100.

Li, X.Z., Roy, C.K., Dong, X., Bolcun-Filas, E., Wang, J., Han, B.W., Xu, J., Moore, M.J., Schimenti, J.C., Weng, Z., et al. (2013). An ancient transcription factor initiates the burst of piRNA production during early meiosis in mouse testes. Mol Cell 50, 67–81.

Mandal, P.K., and Kazazian, H.H., Jr. (2008). SnapShot: Vertebrate transposons. Cell 135, 192–192.e191.

Marçais, G., Delcher, A.L., Phillippy, A.M., Coston, R., Salzberg, S.L., and Zimin, A. (2018). MUMmer4: A fast and versatile genome alignment system. PLoS Comput Biol 14, e1005944.

Martin, M. (2011). <200-1885-3-PB.pdf>. EMBnet Journal 17, 2.

Matsumoto, N., Nishimasu, H., Sakakibara, K., Nishida, K.M., Hirano, T., Ishitani, R., Siomi, H., Siomi, M.C., and Nureki, O. (2016). Crystal Structure of Silkworm PIWI-Clade Argonaute Siwi Bound to piRNA. Cell 167, 484–497.e489.

Miyoshi, K., Miyoshi, T., Hartig, J.V., Siomi, H., and Siomi, M.C. (2010). Molecular mechanisms that funnel RNA precursors into endogenous small-interfering RNA and microRNA biogenesis pathways in Drosophila. Rna 16, 506–515.

Mohn, F., Handler, D., and Brennecke, J. (2015). Noncoding RNA. piRNA-guided slicing specifies transcripts for Zucchini-dependent, phased piRNA biogenesis. Science 348, 812–817.

Molaro, A., Falciatori, I., Hodges, E., Aravin, A.A., Marran, K., Rafii, S., McCombie, W.R., Smith, A.D., and Hannon, G.J. (2014). Two waves of de novo methylation during mouse germ cell development. Genes Dev 28, 1544–1549.

Nishida, K.M., Saito, K., Mori, T., Kawamura, Y., Nagami-Okada, T., Inagaki, S., Siomi, H., and Siomi, M.C. (2007). Gene silencing mechanisms mediated by Aubergine piRNA complexes in Drosophila male gonad. Rna 13, 1911–1922.

Nishida, K.M., Sakakibara, K., Iwasaki, Y.W., Yamada, H., Murakami, R., Murota, Y., Kawamura, T., Kodama, T., Siomi, H., and Siomi, M.C. (2018). Hierarchical roles of mitochondrial Papi and Zucchini in Bombyx germline piRNA biogenesis. Nature 555, 260–264.

Nishimasu, H., Ishizu, H., Saito, K., Fukuhara, S., Kamatani, M.K., Bonnefond, L., Matsumoto, N., Nishizawa, T., Nakanaga, K., Aoki, J., et al. (2012). Structure and function of Zucchini endoribonuclease in piRNA biogenesis. Nature 491, 284–287.

Nishimura T, Nagamori I, Nakatani T, Izumi N, Tomari Y, Kuramochi-Miyagawa S, Nakano T. (2018). PNLDC1, mouse pre-piRNA Trimmer, is required for meiotic and post-meiotic male germ cell development. EMBO Rep. 19, e44957

Ohara, T., Sakaguchi, Y., Suzuki, T., Ueda, H., Miyauchi, K., and Suzuki, T. (2007). The 3’ termini of mouse Piwi-interacting RNAs are 2’-O-methylated. Nat Struct Mol Biol 14, 349–350.

Ou, S., and Jiang, N. (2018). LTR_retriever: A Highly Accurate and Sensitive Program for Identification of Long Terminal Repeat Retrotransposons. Plant Physiol 176, 1410–1422.

Ozata, D.M., Gainetdinov, I., Zoch, A., O’Carroll, D., and Zamore, P.D. (2019). PIWI-interacting RNAs: small RNAs with big functions. Nat Rev Genet 20, 89–108.

Pertea, M., Pertea, G.M., Antonescu, C.M., Chang, T.C., Mendell, J.T., and Salzberg, S.L. (2015). StringTie enables improved reconstruction of a transcriptome from RNA-seq reads. Nat Biotechnol 33, 290–295.

Pezic, D., Manakov, S.A., Sachidanandam, R., and Aravin, A.A. (2014). piRNA pathway targets active LINE1 elements to establish the repressive H3K9me3 mark in germ cells. Genes Dev 28, 1410–1428.

Pillai, R.S., and Chuma, S. (2012). piRNAs and their involvement in male germline development in mice. Dev Growth Differ 54, 78–92.

Popendorf, K., Tsuyoshi, H., Osana, Y., and Sakakibara, Y. (2010). Murasaki: a fast, parallelizable algorithm to find anchors from multiple genomes. PLoS One 5, e12651.

Reuter, M., Berninger, P., Chuma, S., Shah, H., Hosokawa, M., Funaya, C., Antony, C., Sachidanandam, R., and Pillai, R.S. (2011). Miwi catalysis is required for piRNA amplification-independent LINE1 transposon silencing. Nature 480, 264–267.

Robinson, J.T., Thorvaldsdóttir, H., Winckler, W., Guttman, M., Lander, E.S., Getz, G., and Mesirov, J.P. (2011). Integrative genomics viewer. Nat Biotechnol 29, 24–26.

Romanenko, S.A., Perelman, P.L., Trifonov, V.A., and Graphodatsky, A.S. (2012). Chromosomal evolution in Rodentia. Heredity (Edinb) 108, 4–16.

Roovers, E.F., Rosenkranz, D., Mahdipour, M., Han, C.T., He, N., Chuva de Sousa Lopes, S.M., van der Westerlaken, L.A., Zischler, H., Butter, F., Roelen, B.A., et al. (2015). Piwi proteins and piRNAs in mammalian oocytes and early embryos. Cell Rep 10, 2069–2082.

Rosenkranz, D., and Zischler, H. (2012). proTRAC--a software for probabilistic piRNA cluster detection, visualization and analysis. BMC Bioinformatics 13, 5.

Saito, K., Inagaki, S., Mituyama, T., Kawamura, Y., Ono, Y., Sakota, E., Kotani, H., Asai, K., Siomi, H., and Siomi, M.C. (2009). A regulatory circuit for piwi by the large Maf gene traffic jam in Drosophila. Nature 461, 1296–1299.

Saito, K., Sakaguchi, Y., Suzuki, T., Suzuki, T., Siomi, H., and Siomi, M.C. (2007). Pimet, the Drosophila homolog of HEN1, mediates 2’-O-methylation of Piwi-interacting RNAs at their 3’ ends. Genes Dev 21, 1603–1608.

Sasaki, T., Shiohama, A., Minoshima, S., and Shimizu, N. (2003). Identification of eight members of the Argonaute family in the human genome. Genomics 82, 323–330.

Shen, W., Le, S., Li, Y., and Hu, F. (2016). SeqKit: A Cross-Platform and Ultrafast Toolkit for FASTA/Q File Manipulation. PLoS One 11, e0163962.

Sheng, G., Zhao, H., Wang, J., Rao, Y., Tian, W., Swarts, D.C., van der Oost, J., Patel, D.J., and Wang, Y. (2014). Structure-based cleavage mechanism of Thermus thermophilus Argonaute DNA guide strand-mediated DNA target cleavage. Proc Natl Acad Sci U S A 111, 652–657.

Shi, S., Yang, Z.Z., Liu, S., Yang, F., and Lin, H. (2020). PIWIL1 promotes gastric cancer via a piRNA-independent mechanism. Proc Natl Acad Sci U S A 117, 22390–22401.

Sia, S.F., Yan, L.M., Chin, A.W.H., Fung, K., Choy, K.T., Wong, A.Y.L., Kaewpreedee, P., Perera, R., Poon, L.L.M., Nicholls, J.M., et al. (2020). Pathogenesis and transmission of SARS-CoV-2 in golden hamsters. Nature 583, 834–838.

Simon, B., Kirkpatrick, J.P., Eckhardt, S., Reuter, M., Rocha, E.A., Andrade-Navarro, M.A., Sehr, P., Pillai, R.S., and Carlomagno, T. (2011). Recognition of 2’-O-methylated 3’-end of piRNA by the PAZ domain of a Piwi protein. Structure 19, 172–180.

Siomi, M.C., Higashijima, K., Ishizuka, A., and Siomi, H. (2002). Casein kinase II phosphorylates the fragile X mental retardation protein and modulates its biological properties. Mol Cell Biol 22, 8438–8447.

Smibert, P., Yang, J. S., Azzam, G., Liu, J. L., and Lai, E. C. (2013). Homeostatic control of Argonaute stability by microRNA availability. Nat Struct Mol Biol 20, 789–795.

Speir, M.L., Zweig, A.S., Rosenbloom, K.R., Raney, B.J., Paten, B., Nejad, P., Lee, B.T., Learned, K., Karolchik, D., Hinrichs, A.S., et al. (2016). The UCSC Genome Browser database: 2016 update. Nucleic Acids Res 44, D717–725.

Thomson, T., and Lin, H. (2009). The biogenesis and function of PIWI proteins and piRNAs: progress and prospect. Annu Rev Cell Dev Biol 25, 355–376.

Trapnell, C., Williams, B.A., Pertea, G., Mortazavi, A., Kwan, G., van Baren, M.J., Salzberg, S.L., Wold, B.J., and Pachter, L. (2010). Transcript assembly and quantification by RNA-Seq reveals unannotated transcripts and isoform switching during cell differentiation. Nat Biotechnol 28, 511–515.

Treiber, T., Treiber, N., and Meister, G. (2019). Regulation of microRNA biogenesis and its crosstalk with other cellular pathways. Nat Rev Mol Cell Biol 20, 5–20.

Vagin, V.V., Sigova, A., Li, C., Seitz, H., Gvozdev, V., and Zamore, P.D. (2006). A distinct small RNA pathway silences selfish genetic elements in the germline. Science 313, 320–324.

Vargiu, L., Rodriguez-Tomé, P., Sperber, G.O., Cadeddu, M., Grandi, N., Blikstad, V., Tramontano, E., and Blomberg, J. (2016). Classification and characterization of human endogenous retroviruses; mosaic forms are common. Retrovirology 13, 7.

Vaser, R., Sović, I., Nagarajan, N., and Šikić, M. (2017). Fast and accurate de novo genome assembly from long uncorrected reads. Genome Res 27, 737–746.

Vasiliauskaitė L, Berrens RV, Ivanova I, Carrieri C, Reik W, Enright AJ, and O’Carroll D. (2018). Defective germline reprogramming rewires the spermatogonial transcriptome. Nat Struct Mol Biol. 25, 394–404

Vourekas, A., Zheng, Q., Alexiou, P., Maragkakis, M., Kirino, Y., Gregory, B.D., and Mourelatos, Z. (2012). Mili and Miwi target RNA repertoire reveals piRNA biogenesis and function of Miwi in spermiogenesis. Nat Struct Mol Biol 19, 773–781.

Wang, Y., Sheng, G., Juranek, S., Tuschl, T., and Patel, D.J. (2008). Structure of the guide-strand-containing argonaute silencing complex. Nature 456, 209–213.

Waterhouse, R.M., Seppey, M., Simão, F.A., Manni, M., Ioannidis, P., Klioutchnikov, G., Kriventseva, E.V., and Zdobnov, E.M. (2018). BUSCO Applications from Quality Assessments to Gene Prediction and Phylogenomics. Mol Biol Evol 35, 543–548.

Williams, Z., Morozov, P., Mihailovic, A., Lin, C., Puvvula, P.K., Juranek, S., Rosenwaks, Z., and Tuschl, T. (2015). Discovery and Characterization of piRNAs in the Human Fetal Ovary. Cell Rep 13, 854–863.

Yamaguchi, S., Oe, A., Nishida, K.M., Yamashita, K., Kajiya, A., Hirano, S., Matsumoto, N., Dohmae, N., Ishitani, R., Saito, K., et al. (2020). Crystal structure of Drosophila Piwi. Nat Commun 11, 858.

Yang, Q., Li, R., Lyu, Q., Hou, L., Liu, Z., Sun, Q., Liu, M., Shi, H., Xu, B., Yin, M., et al. (2019). Single-cell CAS-seq reveals a class of short PIWI-interacting RNAs in human oocytes. Nat Commun 10, 3389.

